# The joint evolution of separate sexes and sexual dimorphism

**DOI:** 10.1101/2024.05.31.596835

**Authors:** Thomas Lesaffre, John R. Pannell, Charles Mullon

## Abstract

Dioecious plants are frequently sexually dimorphic. Such dimorphism, which reflects responses to selection acting in opposite directions for male and female components of fitness, is commonly thought to emerge after separate sexes evolved from hermaphroditism. But associations between allocation to male and female function and traits under sexual conflict may well also develop in hermaphroditic ancestors. Here, we show that variation in sex allocation and a trait under sexual conflict inevitably generates an advantage to sexual specialisation, fueling the transition to dioecy. In the absence of constraints, this leads to the joint evolution of separate sexes and sexual dimorphism through the build-up of an association between sex allocation and the conflict trait, such that eventually the population consists of unisexuals expressing their sex-specific optima. We then investigate how such association might materialise genetically, either via recombination suppression or via sex-dependent expression, and show that the genetic architecture of sex allocation and the conflict trait readily evolves to produce the association favoured by selection. Finally and in agreement with previous theory, we demonstrate that limited dispersal and self-fertilisation, which are pervasive ecological characteristics of flowering plants, can offset the advantage of sexual specialisation generated by sexual conflict and thus maintain hermaphroditism. Taken together, our results indicate that advantages to sexual specialisation are inevitable when there is conflict between sexual functions in hermaphrodites, but these advantages can be counterbalanced by ecological benefits of hermaphroditism.

## 1 Introduction

The vast majority of flowering plant species are hermaphroditic, but dioecy (separate sexes) has evolved many times from this ancestral state (Charlesworth, 1985; Renner and Ricklefs, 1995; Barrett, 2002a; Renner, 2014; Käfer et al., 2017). Indeed, dioecious species are rare (about 6% of flowering plants) but widely distributed across the Angiosperm phylogeny, suggesting that hundreds -possibly thousands -of independent transitions to dioecy have taken place in flowering plants (Charlesworth, 1985; Renner and Ricklefs, 1995; Renner, 2014). Evolutionary transitions from hermaphroditism to dioecy are thought to occur along two main pathways (Lloyd, 1980; Charlesworth, 1999; Pannell and Jordan, 2022). First, dioecy may evolve in a stepwise process involving gynodioecy (the coexistence of females and hermaphrodites in a population) as an intermediate state. Theory for this pathway is based on population genetics models that consider the spread of male and female sterility mutations in an initially hermaphroditic population (Charlesworth and Charlesworth, 1978a,b). These models show that partial selfing and inbreeding depression can facilitate the spread of a male-sterility mutation, leading to gynodioecy. In turn, the presence of females in the population can favour the invasion of one or more mutations turning hermaphrodites into males, thereby establishing dioecy.

Second, dioecy may evolve via the gradual divergence of increasingly male- and female-biased morphs that eventually become pure males and females (Lloyd, 1980; Charlesworth, 1999). This pathway, which is sometimes referred to as the ‘monoecy-paradioecy’ pathway (Lloyd, 1980), has been studied through the evolution of sex allocation, i.e., the division of limited reproductive resources between male and female functions (Charnov et al., 1976; Charnov, 1982; West, 2009). In sex allocation theory, how resource allocation translates into fitness depends on the shapes of the male and female ‘gain curves’, which are functions that relate the amount of resources allocated to one sex to fitness gained through that sex. When a curve is linear, one additional unit of resources invested in one sex results in an increase of one unit of fitness through that sex; when a curve is accelerating, the fitness gained increases at an increasing rate with additional investment, whereas when it is saturating, the fitness gained increases at a decreasing rate with additional investment. The central prediction of this theory is that saturating gain curves allow the maintenance of hermaphroditism, because such curves lead individuals to accrue diminishing fitness returns as they allocate more resources to a given sex function, whereas accelerating gain curves promote the evolution of dioecy, because individuals then accrue increasing fitness returns from specialising into one sex (Charnov et al., 1976).

Several ecological mechanisms have been proposed to explain when we should see accelerating gain curves and thus fitness benefits to sexual specialisation (Thomson and Brunet, 1990; Charlesworth and Morgan, 1991; Freeman et al., 1997; Charlesworth, 1999; Pannell and Jordan, 2022). For example, individuals allocating more to their male function may be able to substantially increase their floral display. This increase could in turn enhance their attractiveness to pollinators, and result in more efficient pollen dispersal. Such coupling between pollen dispersal and production could generate multiplicative benefits to specialising as a male, i.e., an accelerating male gain curve (Bawa, 1980; Ohashi and Yahara, 2001; see also Chap. 3 in De Jong and Klinkhamer, 2005). For female function, there are three main ideas to account for possible accelerating gain curves. First, in plants with animal-dispersed fruits, individuals allocating more heavily to their female function may enjoy more efficient dispersal of their seeds if animals are more attracted to plants bearing larger fruit crops (Bawa, 1980; Givnish, 1980). This coupling of seed dispersal with sex allocation could cause the female gain curve to accelerate by reducing kin competition among seeds of more female individuals (Vamosi et al., 2007; Biernaskie, 2010). Second, seeds of individuals producing larger crops may sometimes enjoy a lower predation risk due to ‘predator satiation’ (Janzen, 1971; Lloyd, 1982), which may cause the female gain curve to accelerate through the coupling of seed survival and sex allocation. Third, if the selfing rate in hermaphrodites increases with pollen production, so that selfing is coupled with sex allocation, then high inbreeding depression may lead to multiplicative benefits of specialising in female function, because more female individuals would then produce seeds that are both more numerous and more outcrossed (and thus more fit, Charlesworth and Charlesworth, 1978b; De Jong and Geritz, 2001; Lesaffre et al., 2024).

Previous ecological explanations for accelerating gain curves allow for features of plant ecology, such as the efficiency of pollen and seed dispersal, to vary as a direct consequence of a plant’s allocation to male versus female function, but all such explanations assume that sex allocation is the only evolving trait. However, transitions to dioecy are typically accompanied by the emergence of trait differences between male and female individuals that go beyond the production of pollen or ovules. Indeed, sexual dimorphism is widespread in dioecious plants, with males and females typically differing in several life-history (e.g. lifespan, growth rate or phenology) and reproductive traits (e.g. flowering time or flower size, position, shape and number, Lloyd and Webb, 1977; Geber, 1999; Barrett and Hough, 2013). Sexual dimorphism is commonly thought to be the result of the resolution of conflicts over trait expression between males and females, and therefore to arise after separate sexes have evolved (Barrett and Hough, 2013; Charlesworth, 2018). But these conflicts were likely already at work in the hermaphroditic ancestor, especially over reproductive traits, because what is optimal for reproduction through male function might often not be for reproduction through female function (Geber, 1999; Delph and Ashman, 2006). In wind-pollinated plants, for instance, efficient pollen dispersal by stamens and pollen capture by stigmas require floral traits with opposite biophysical properties. Pollen dispersal requires that flowers intermittently interrupt the airstream by shaking in the wind, allowing pollen to escape, whereas pollen capture requires flowers to act as fixed obstacles on which pollen can get stuck (Niklas, 1985; Freeman et al., 1997; Barrett, 2002b; Timerman and Barrett, 2021). Another example is floral display in animal-pollinated plants (Barrett and Hough, 2013). Pollinators are typically attracted to plants with higher floral displays, and both sexual functions need to attract pollinators. But whereas increased attractiveness is expected to benefit male function through increased pollen dispersal, female function might eventually suffer from increased pollinator attraction if attractiveness increases risks of geiotonogamous selfing and of flower damage, which could compromise seed development (Ohashi and Yahara, 2001; De Jong and Klinkhamer, 2005). Conflicts over the expression of traits between sexual functions in hermaphrodites could thus promote the evolution of separate sexes as a means of conflict resolution. In other words, the joint evolution of sex allocation and traits under sexual conflict could generate benefits to sexual specialisation, and lead to the simultaneous, rather than sequential, emergence of separate sexes and sexual dimorphism (Willson, 1979; Geber, 1999).

The joint emergence of separate sexes and sexual dimorphism has so far been studied in the context of the gynodioecy pathway, i.e., of the spread of unisexuals in a population of hermaphrodites. Several models have investigated the effect of a joint change in sexual phenotype together with another trait such as floral display, allocation to growth vs. reproduction, or to attractive structures vs. gamete production, and have shown that it can facilitate invasion by unisexuals (Morgan, 1992; Seger and Eckhart, 1996; Sato, 2002). However, these models assume that invading unisexuals already express the trait value that is optimal for their sex, which is biologically equivalent to assuming either that sterility mutations are pleiotropic in a way that is phenotypically optimal, or that the trait is plastic and expressed conditional on the sexual phenotype. More recently, Olito and Connallon (2019) studied the effect of sexually antagonistic alleles segregating at a partially linked locus on the ability of male and female sterility mutations to invade a hermaphroditic population (see also Blackburn et al. (2010) for the effect of sexually antagonistic variation on the evolution of sex-ratio adjustment in dioecious species). They found that sexually antagonistic variation at a nearby locus could facilitate the spread of sterility mutations through hitchhiking, demonstrating that polymorphism maintained by sexual conflict can aid the establishment of unisexuals. Their approach assumes a constant recombination rate between the two loci concerned and does not consider the emergence of a phenotypic association between sex and a trait, so that sexual dimorphism cannot emerge jointly with separate sexes in their model.

Here, we investigate the joint emergence of separate sexes and sexual dimorphism along the monoecy-paradioecy pathway. We consider the joint evolution of sex allocation and a continuous trait with sex-specific optima (so that it is under sexual conflict). We first use a baseline model to show that selection favours sex allocation and the conflict trait to become associated, thereby generating an advantage to sexual specialisation and promoting the evolution of separate sexes. We then extend our baseline model to explore how this association might come about at a genetic level. Our analyses reveal that the genetic architecture of sex allocation and the conflict trait readily evolves to produce the association favoured by selection, suggesting that genetic constraints are unlikely to constitute a long-term barrier to the joint evolution of separate sexes and sexual dimorphism. Finally, we show that ecological characteristics that are pervasive among flowering plants, namely limited dispersal and self-fertilisation, can offset the advantage of sexual specialisation driven by variation in sex allocation and a trait under sexual conflict, and maintain hermaphroditism. Taken together, our results indicate that traits under sexual conflict in hermaphrodites can favour the gradual emergence of separate sexes and sexual dimorphism, shedding further light on the possible basis and dynamics of transitions to dioecy in flowering plants.

## 2 Sexual conflict in hermaphrodites always favours sexual specialisation

We first propose a simple baseline model to investigate the joint evolution of separate sexes and sexual dimorphism.

### 2.1 Baseline model

Our baseline model considers a large population of haploid hermaphrodites with the following life cycle. *(i) Sexual development:* First, individuals undergo sexual development and produce a large number of female and male gametes. *(ii) Mating:* Female and male gametes then fuse randomly, leading to the production of a large number of diploid zygotes that divide to give rise to two haploid seeds each. *(iii) Recruitment:* Finally, adults die and are replaced by juveniles sampled uniformly from the produced seeds. The assumption that individuals are haploid is made mainly for mathematical convenience, but also because it allows us to focus on the interaction between loci rather than within loci for now. It also means that our baseline model readily applies to haplontic species, such as algae (which are discussed in section 5). We consider the effects of diploidy in section 3.3.

Each individual is characterised by two continuous traits, *x* ∈ [0, 1] and *z* ∈ ℝ, which determine their fecundity through female and male functions. The first trait *x* is sex allocation, i.e., *x* is the proportion of its reproductive resources that an individual allocates to its female function during sexual development (so that the remaining 1 − *x* is allocated to its male function). Meanwhile the second trait, which we will refer to as ‘trait *z*’ or the ‘conflict trait’ for short, is a continuous trait that affects reproduction differently through female and male function. This conflict trait could be a trait that enhances reproductive success through one sexual function at the expense of the other, such as floral stem length in wind-pollinated plants, or traits related to pollinator attraction in animal-pollinated plants (Delph, 1996; Friedman and Barrett, 2009); it could also represent a physiological trade-off underlying the expression of distinct male- and female-beneficial phenotypes.

Overall, the effective fecundity of individual *i* with traits **y**_*i*_ = (*x*_*i*_, *z*_*i*_) through its female and male function are given by

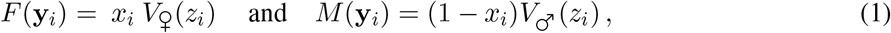

respectively, where *V*♀ (*z*_*i*_) and *V*♂ (*z*_*i*_) are functions capturing the effect of trait *z* on reproduction through female and male functions. We assume that *V*♀ (*z*_*i*_) and *V*♂ (*z*_*i*_) are positive, smooth and continuous functions, each with a unique maximum. Without loss of generality, we assume that this maximum is larger for female than for male function, so that the optimal trait value is greater for female than for male function. We derive most of our results with these general functions, but where necessary we assume that *V*♀(*z*_*i*_) and *V* ♂ (*z*_*i*_) are Gaussian, i.e.,

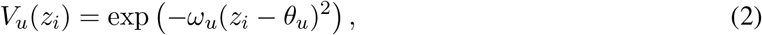

with *u* ∈ {♀, ♂}, and where *θ*_*u*_ ∈ ℝ and *ω*_*u*_ *>* 0 denote the optimal trait value and the strength of selection in sex *u*, respectively.

We assume that sex allocation *x* and trait *z* are each determined by a quantitative trait locus that evolves via the input of rare mutations with weak phenotypic effects (‘continuum-of-alleles’ model; Kimura, 1965, p. 883 in Walsh and Lynch, 2018). We assume for now that the loci encoding *x* and *z* are fully-linked (i.e., no recombination; we later relax this assumption). We analyse this model in detail in Appendix A and summarise our main results below.

### 2.2 Joint evolutionary dynamics of sex allocation and a trait under sexual conflict

When mutations are rare and have weak phenotypic effects, trait evolution can be decomposed into two phases (Geritz et al., 1998). First, the population evolves under directional selection, whereby positively selected mutations sweep to fixation so that the population transits from being largely monomorphic for one trait value to being monomorphic for another. Through the sequential fixation of mutations altering sex allocation *x* and trait *z* in the direction favoured by selection, we show in Appendix A.1.1 that the population eventually converges to the equilibrium **y**^*^ = (*x*^*^, *z*^*^) where

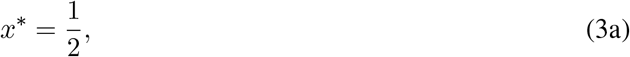

and *z*^*^ is such that,

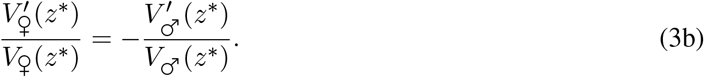

Therefore, at this equilibrium, individuals divide their reproductive resources equally between their female and male functions (eq. 3a) and express a trait value *z*^*^ such that the marginal fitness gained through male and female functions compensate each other (eq. 3b).

Once the population expresses **y**^*^, it either remains monomorphic or becomes polymorphic in a process known as ‘evolutionary branching’ in which two discrete morphs gradually diverge from one another (Geritz et al., 1998; Paásztor et al., 2016). Whether or not this occurs depends on the so-called Hessian matrix,

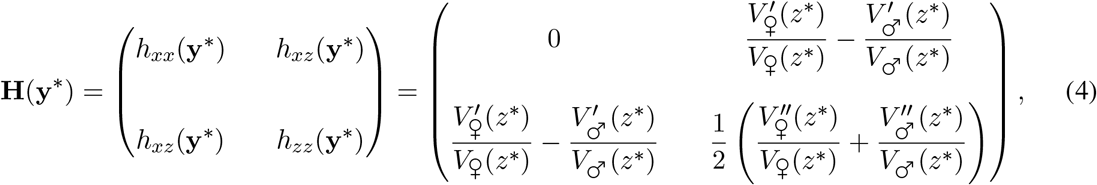

with the emergence of polymorphism requiring that the leading eigenvalue of **H**(**y**^*^) is positive (Leimar, 2009; Deébarre et al., 2014; Geritz et al., 2016, see eq. A12 in Appendix A.1.2 for the mathematical definition of **H**(**y**^*^)). Before determining the conditions for polymorphism from this matrix, we note that each entry of **H**(**y**^*^) captures a different aspect of selection (Lande and Arnold, 1983). Diagonal entries *h*_*xx*_(**y**^*^) and *h*_*zz*_(**y**^*^), which are the quadratic selection coefficients on sex allocation *x* and trait *z*, respectively, tell us whether selection on each trait is disruptive (if positive) or stabilising (if non-positive) when traits evolve independently from one another. Meanwhile, the off-diagonal entry *h*_*xz*_(**y**^*^) is the coefficient of correlational selection acting on *x* and *z*. A positive (resp. negative) *h*_*xz*_(**y**^*^) indicates that selection favours a positive (negative) association between the two traits when they evolve jointly. Using the fact the **H**(**y**^*^) is symmetric, we can identify two ways for its leading eigenvalue to be positive and thus for selection to favour polymorphism: *(i)* either one of the diagonal entries is positive and thus one trait is independently experiencing disruptive selection, i.e.,

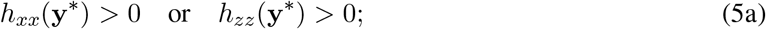

or, *(ii)* if condition (5a) does not hold, then polymorphism is favoured when

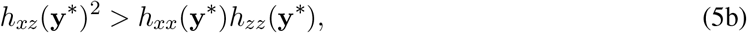

i.e., when correlational selection is sufficiently strong to overcome stabilising selection acting on each trait independently.

Because *h*_*xx*_(**y**^*^) = 0 for our baseline model (top-left element in eq. 4), selection is stabilising on sex allocation when *x* evolves independently (and *z* is fixed). This is because we assumed a linear trade-off between male and female allocation, such that there is no inherent advantage to sexual specialisation (Charnov et al., 1976). When trait *z* evolves independently (i.e., when sex allocation is fixed at *x*^*^ = 1*/*2), inspection of *h*_*xx*_(**y**^*^) reveals that experiences disruptive selection when 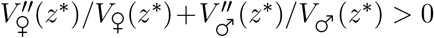, i.e., when *V*♀ (*z*) and *V*♂ (*z*) are on average accelerating when *z* = *z*^*^. In this case, conflict between female and male trait expression is so strong that it favours the emergence of two hermaphroditic morphs that allocate equally into their female and male functions (*x*^*^ = 1*/*2), with one morph expressing the optimal-female trait value (such that 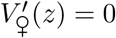) and the other morph expressing the male-optimum (such that 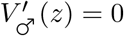).

Plugging the entries of **H**(**y**^*^) (eq. 4) into condition (5b), we see that this condition always holds in our baseline model. This indicates that selection always favours polymorphism when both sex allocation *x* and trait *z* evolve jointly, even when each trait experiences stabilising selection in isolation. In other words, hermaphroditism is never evolutionarily stable here. Using numerical analyses and individual-based simulations to investigate the polymorphism that results from correlational selection (Appendices A.2 and A.3 for details), we find that, eventually, the population consists of two types: *(i)* pure females that express the optimal trait value for their reproduction (i.e., *x* = 1 and *z* such that 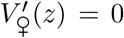; and *(ii)* pure males that express the optimal trait value for their reproduction (i.e., *x* = 0 and *z* such that 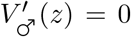, as shown in Figure 1. Selection thus always leads to the joint emergence of separate sexes and sexual dimorphism in our baseline model.

**Figure 1.**
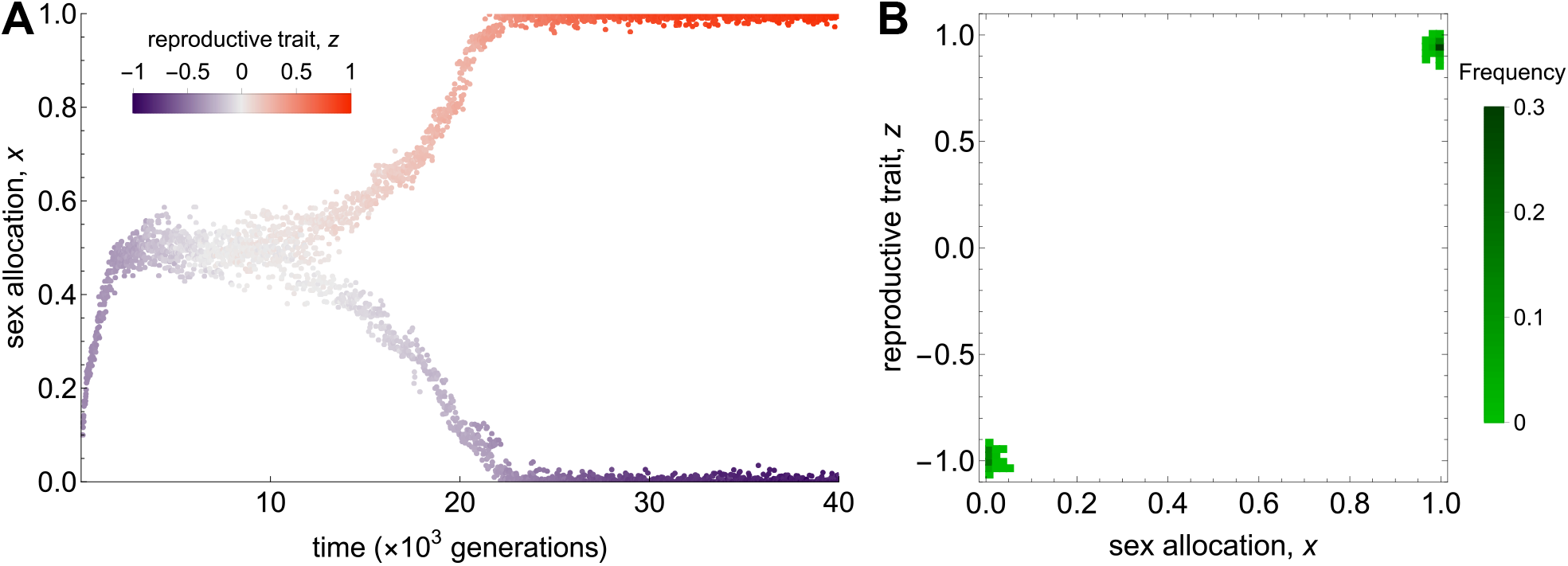
Joint evolutionary dynamics of sex allocation *x* and trait *z* in the baseline model leading to the simultaneous emergence of separate sexes and sexual dimorphism. Simulations assume Gaussian fecundity functions (eq. 2, Appendix A.3 for details on the simulation process). **A** Sex allocation *x* (y-axis) and trait *z* (colour scale) of three randomly sampled individuals every 10 generations as a function of time. Orange indicates more female-beneficial trait values, whereas purple indicates more male-beneficial values. The population first converges to the equilibrium **y**^*^ = (*x*^*^, *z*^*^), where *x*^*^ = 1*/*2 and *z*^*^ = 0, and then sees the gradual emergence of two morphs, a female morph (*x* = 1) expressing the female-optimal trait value *θ*♀ = 1, and a male morph expressing the male-optimal trait value *θ*♂ = −1. **B** Distribution of phenotypes averaged over the last 1, 000 generations of the simulation. Darker colours indicate higher frequencies. Parameters used in simulations: *θ*♀ = 1, *θ*♂ = −1, *ω* ♀= *ω*♂= 1*/*6, *N* = 5 *×*10^3^, *µ* = 5 *×*10^−3^, and standard deviation *σ* = 10^−2^.

For the rest of our manuscript, we extend our baseline model to explore potential factors that may maintain hermaphroditism against this selective advantage of sexual specialisation. These factors are broadly classified as to whether they are genetic (Section 3) or ecological (Section 4).

## 3 Genetic constraints and their resolution

The transition to dioecy due to correlational selection described in the preceding section relies on the build-up of an association between sex allocation *x* and trait *z*. In our baseline model (Section 2.1), this is achieved by linkage between alleles at the two respective loci that improve reproductive success through the same sexual function. These genetic associations between co-adapted alleles are free to build up here, because loci influencing *x* and *z* do not recombine.

We ran simulations allowing for recombination at a fixed rate *r* ∈ [0, 1*/*2] between the loci during the diploid zygotic stage. We found that low recombination may still allow the emergence of polymorphism, albeit leading to the production of unfit recombinant females and males that express a trait *z* value that is optimal for the sex opposite to theirs (Fig. 2A). However, higher recombination rates prevent polymorphism from emerging entirely (Figure 2B-C). This is because recombination breaks the positive genetic association on which correlational selection acts. Nevertheless, correlational selection should favour the evolution of genetic architectures that increase the heritability of beneficial associations between *x* and *z*. We investigate two potential routes for this to occur. First, we model the evolution of recombination between the two loci influencing *x* and *z* (Section 3.1). Second, we study the evolution of sex allocation-dependent expression of trait *z* (Section 3.2). Finally, we include diploidy in the adult stage in our model, and explore how the additional genetic constraint this poses can be resolved through the evolution of dominance (Section 3.3).

**Figure 2.**
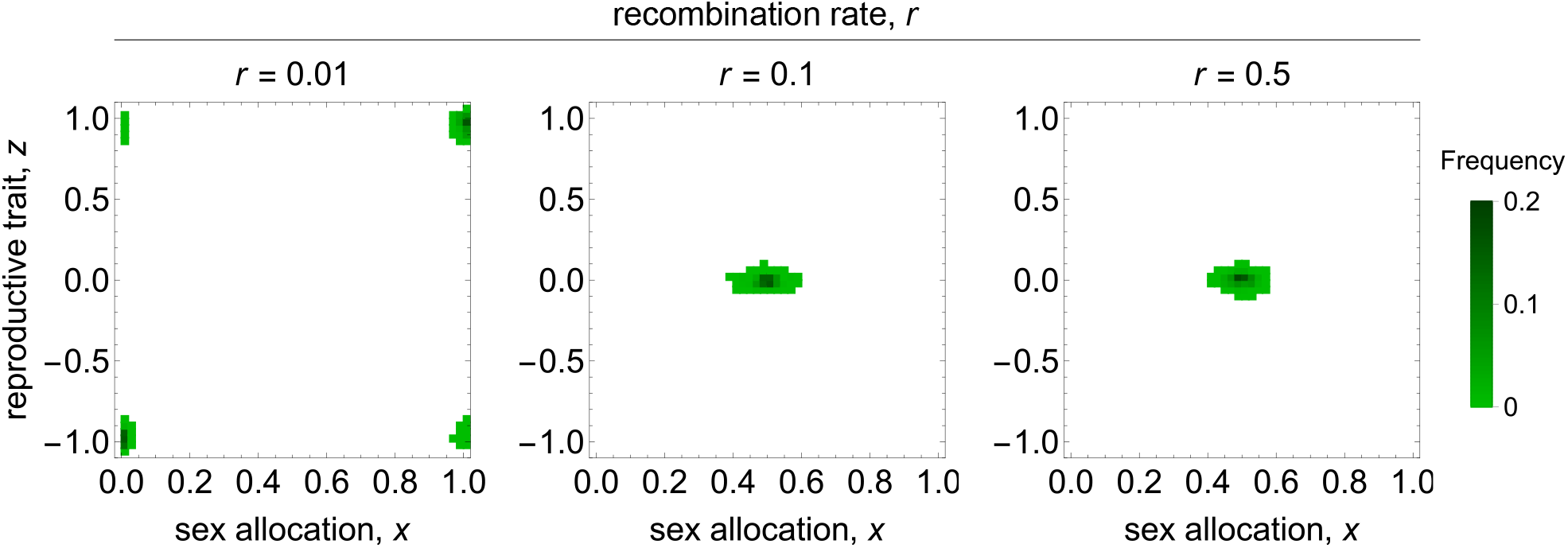
Recombination breaks the necessary genetic associations for the joint evolution of separate sexes and sexual dimorphism. Panels show the distribution of phenotypes expressed in simulations populations averaged over 25, 000 generations after a 50, 000 generations burn-in period, with higher densities indicated by darker colours. Results are shown for three recombination rates ranging from *r* = 0.01 on the left-hand side to *r* = 0.5 (free recombination) on the right-hand side. Parameters used in simulations: *θ* ♀ = 1, *θ*♂ = −1, *ω*♀ = *ω*♂ = 0.4, *N* = 5 *×* 10^3^, *µ* = 10^−3^, *σ* = 10^−2^.

### 3.1 Evolution of recombination

We extend our baseline model (Section 2.1) to include recombination evolution. We assume that the recombination rate in a zygote is controlled by an unlinked modifier locus at which alleles are additive and subject to rare, small-effect mutations, so that *r* evolves gradually (Appendix B.1.1 for details).

We first investigate recombination evolution by considering the fate of a rare mutant allele coding for a recombination rate *r*_m_ at the modifier, in a population otherwise fixed for rate *r*. We assume that two alleles segregate at the sex allocation locus. We label these alleles by their phenotypic effect *x*♀ and *x*♂, where *x*♂ *< x*♀. Similarly, two alleles *z*♀ and *z*♂ (*z*♂ *< z*♀) segregate at the locus encoding trait *z*. Therefore, there are four possible haplotypes in the population: two ‘adapted’ haplotypes carrying matching alleles, with one haplotype adapted to female reproduction **y**_1_ = (*x*♀, *z*♀), and the other to male reproduction **y**_2_ = (*x*♂, *z*♂); and two ‘maladapted’ haplotypes that carry mismatched alleles **y**_3_ = (*x*♀, *z*♂) and **y**_4_ = (♂ ♀).

We show in Appendix B.1.2 that the selection gradient on *r*, which gives the direction and intensity of selection on recombination in a population expressing *r*, can be written in the same form as more general theory on the evolution of recombination (eq. A1.5e in Barton, 1995), i.e., as

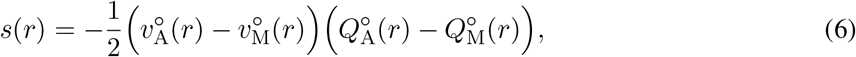

where 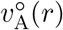 and 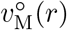 are the mean reproductive values of adapted and maladapted haplotypes, respectively (eq. B9b), which measure the asymptotic demographic contribution of individuals carrying a given haplotype to the future population (for a mathematical definition, see e.g. p. 153 in Rousset, 2004), and 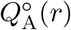 and 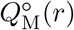 are the frequencies of double-heterozygous zygotes carrying adapted and maladapted haplotypes, i.e., of zygotes with genotypes **y**_1_*/***y**_2_ and **y**_3_*/***y**_4_, respectively (only double-heterozygous zygotes appear in eq. 6, as these zygotes are those in which recombination has an effect on the frequency of haplotypes in the produced seeds; see eqs. B9c-B9d in Appendix B.1.2.2). The first term in brackets in eq. (6) is the difference in reproductive values between adapted and maladapted haplotypes, which quantifies epistasis between the loci encoding sex allocation and trait *z* (in fact, this term is analogous to the epistasis term *ϵ*_*jk*_ in eq. A1.5e of Barton, 1995). Given that adapted haplotypes have higher fitness, this term is always positive (i.e., there is positive epistasis). The second term in brackets in eq. (6) measures the difference in frequency between double-heterozygous zygotes carrying adapted vs. maladapted haplotypes, i.e., linkage disequilibrium between the loci encoding sex allocation and the trait (*C*_*jk*_ in eq. A1.5e of Barton, 1995). This is also positive, as adapted haplotypes are maintained at higher frequencies and so form more zygotes. Therefore, eq. (6) is always negative, and, as expected from classical theory on the evolution of recombination (Barton, 1995), selection always favours a decrease in the recombination rate between the loci encoding *x* and *z* here. This should eventually lead to complete recombination suppression (*r* → 0).

We checked this prediction with individual-based simulations in which sex allocation *x*, trait *z* and recombination rate *r* all evolve. Loci for *x* and *z* are initially freely recombining (i.e., 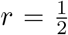 at the beginning of each simulation run; the program is described in detail in Appendix B.1.3). As predicted, recombination suppression always evolves in these simulations (Fig. 3A). This allows for positive genetic linkage to build up between the loci encoding *x* and *z*, and thus for the simultaneous evolution of separate sexes and sexual dimorphism (Fig. 3B-C). However, full recombination suppression can take a long time to evolve. This is because selection on the recombination rate is proportional to the divergence between alleles at the loci encoding *x* and *z*, and so is initially weak. Recombination can thus delay transitions to dioecy in the shortterm, but it is unlikely to prevent them from taking place in the long-term, as mutations and chromosomal rearrangements leading to recombination suppression are always favoured by selection.

**Figure 3.**
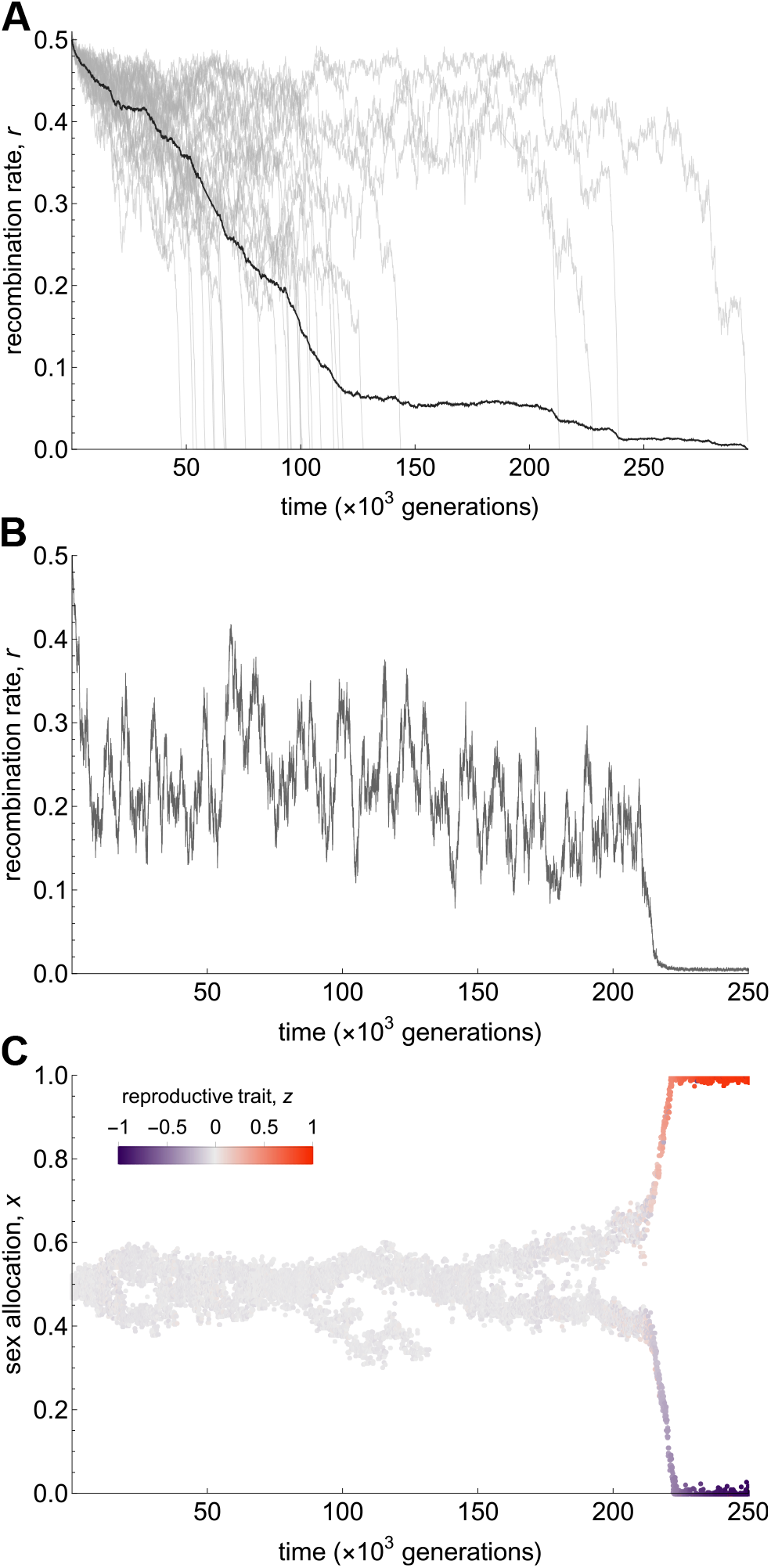
Evolution of recombination suppression. **A** Recombination rate as a function of time in 30 replicate populations where the recombination rate *r* evolves jointly with sex allocation *x* and trait *z* (Appendix B.1.3 for details on the simulation process). Each grey line corresponds to one replicate. The black line indicates the average trajectory. Parameters used in simulations: *θ*♀ = 1, *θ*♂ =− 1, *ω*♀ = *ω*♂ = 0.4, *N* = 10^4^, *µ* = 5 *×* 10^−3^, *σ* = 10^−2^ for all three evolving traits (*x, z* and *r*). **B-C** Recombination rate (panel B) and phenotypes expressed (panel C) in a single replicate as a function of time. In panel C, sex allocation is given on the y-axis, and trait *z* by the colour scale (as in Fig. 1). This shows that recombination suppression allows the simultaneous emergence of separate sexes and sexual dimorphism. Parameters used in simulations: *θ*♀ = 1, *θ*♂ = −1, *ω*♀ = *ω*♂ = 0.4, *N* = 10^4^, *µ* = 10^−3^, *σ* = 10^−2^ for *x* and *z*, and *µ* = 5 *×* 10^−3^, *σ* = 5 *×* 10^−2^ for *r*).

### 3.2 Evolution of conditional trait expression

Another potential route for sex allocation *x* and trait *z* to become associated is through the evolution of conditional expression of *z* on *x*. This would allow individuals that allocate more to a given sexual function to express traits that are better suited to this function, but does not require physical linkage between the loci encoding *x* and *z*.

To investigate this possibility, we now let trait *z* of an individual with sex allocation *x* be given by

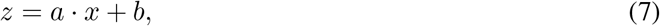

where *b* ∈ ℝ is the component of *z* that is independent of sex allocation, and *a* ∈ ℝ is the sex allocation-dependent component of *z*. When *a* = 0, *z* is determined by *b* and is therefore independent of sex allocation. In contrast, *z* increases in more female individuals when *a >* 0 (and decreases when *a <* 0). We assume that there is a cost to plasticity, which could for instance be due to the cost of maintaining the additional physiological and regulatory machinery required for conditional trait expression (DeWitt et al., 1998). This cost decreases fecundity for individuals with larger sex allocation-dependent trait components (i.e., larger |*a*|). Specifically, we assume that female and male fecundities of an individual expressing **y**_*i*_ = (*x*_*i*_, *a*_*i*_, *b*_*i*_) are now given by

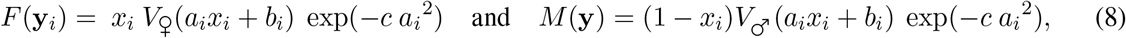

where *c >* 0. Sex allocation *x* and trait *z* components *a* and *b* are assumed to be encoded by unlinked loci that each evolve according to a ‘continuum-of-alleles’ model.

We show in Appendix B.2.1 that, first, directional selection leads the population to express the equilibrium **y**^*^ = (*x*^*^, *a*^*^, *b*^*^), where

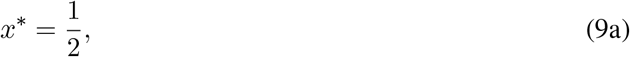

so that individuals divide their reproductive resources equally between their male and female function. Mean-while

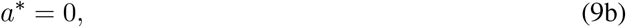

and *b*^*^ is such that

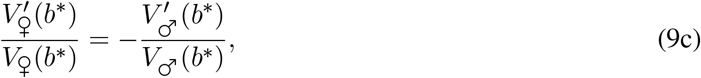

i.e., such that individuals express a trait value *z*^*^ = *b*^*^ that is independent of sex allocation (*a*^*^ = 0) and sits between the female and male optima, as in our baseline model (eq. 3).

Once the population expresses **y**^*^, we find that two distinct cases need to be considered. The first is when functions *V*♀(*z*) and *V*♂(*z*) are saturating on average, i.e., when

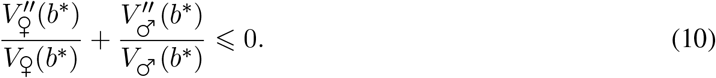

In this case, quadratic selection is stabilising on all three traits, and so the population should remain uni-modally distributed around **y**^*^ (recall that because loci are unlinked here, genetic associations are constantly being removed by recombination so that correlational selection cannot act). Nevertheless, standing variation for sex allocation around *x*^*^ maintained at mutation-selection-drift balance may generate selection on trait components *a* and *b*. To investigate this, we assume that two sex allocation alleles weakly diverging from *x*^*^, denoted *x*♀ = *x*^*^ + Δ*x* and *x*♂ = *x*^*^ − Δ*x*, where Δ*x >* 0 is small, segregate. We study selection on *a* and *b* assuming Gaussian fecundity functions for simplicity (eq. 2; see Appendix B.2.2 for details on the analysis).

We find that selection leads the polymorphic population to evolve sex allocation-dependent trait expression, with the sex-allocation dependent and independent components of *z* evolving to

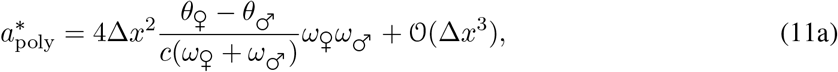

and

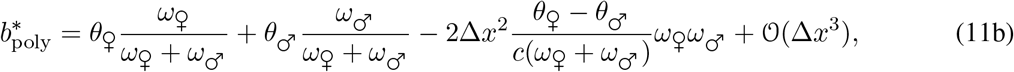

so that individuals that are more female (*x x*^*^) express larger female-beneficial *z* values, and individuals that are more male (*x < x*^*^) express smaller male-beneficial *z* values. As indicated by eq. (11), the dependence of *z* on sex allocation that evolves is stronger when sex-specific selection is stronger (i.e., as *ω*♀ and *ω*♂ increase, or the distance between male and female optima |*θ*♀ − *θ*♂ | increases), and when the cost of plasticity is small (*c* decreases), as individuals then pay a smaller fecundity cost from expressing a larger *a*.

We show that, in turn, sex allocation-dependent expression of *z* generates selection for alleles *x*♀ and *x*♂ to diverge further (eq. B67 in Appendix B.2.2). Specifically, we show that the selection gradients *s*♀(**y**♀, **y**♂) and *s* (**y**♀, **y**) on each sex allocation allele are given by

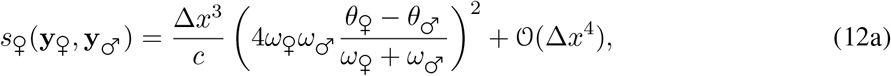

and

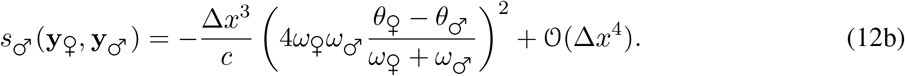

These gradients are positive and negative, respectively, indicating that selection favours the female-biased allele *x*♀ to become more female-biased (i.e., to code for larger values of *x*) and the male-biased allele *x*♂ to become more male-biased (i.e., to code for smaller values of *x*).

Selection on conditional trait expression and sex allocation should therefore reinforce each other, leading to the joint emergence of separate sexes and sexual dimorphism. To check this, we used individual-based simulations (Appendix B.2.3 for details on the simulation program). These simulations always show the emergence of separate sexes and sexual dimorphism (Figure 4), with trait components *a* and *b* evolving to

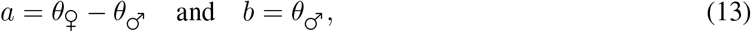

such that females (*x* = 1) express the female optimum *z* = *θ*♀, and males (*x* = 0) express the male optimum *z* = *θ*♂ (recall eq. 7).

**Figure 4.**
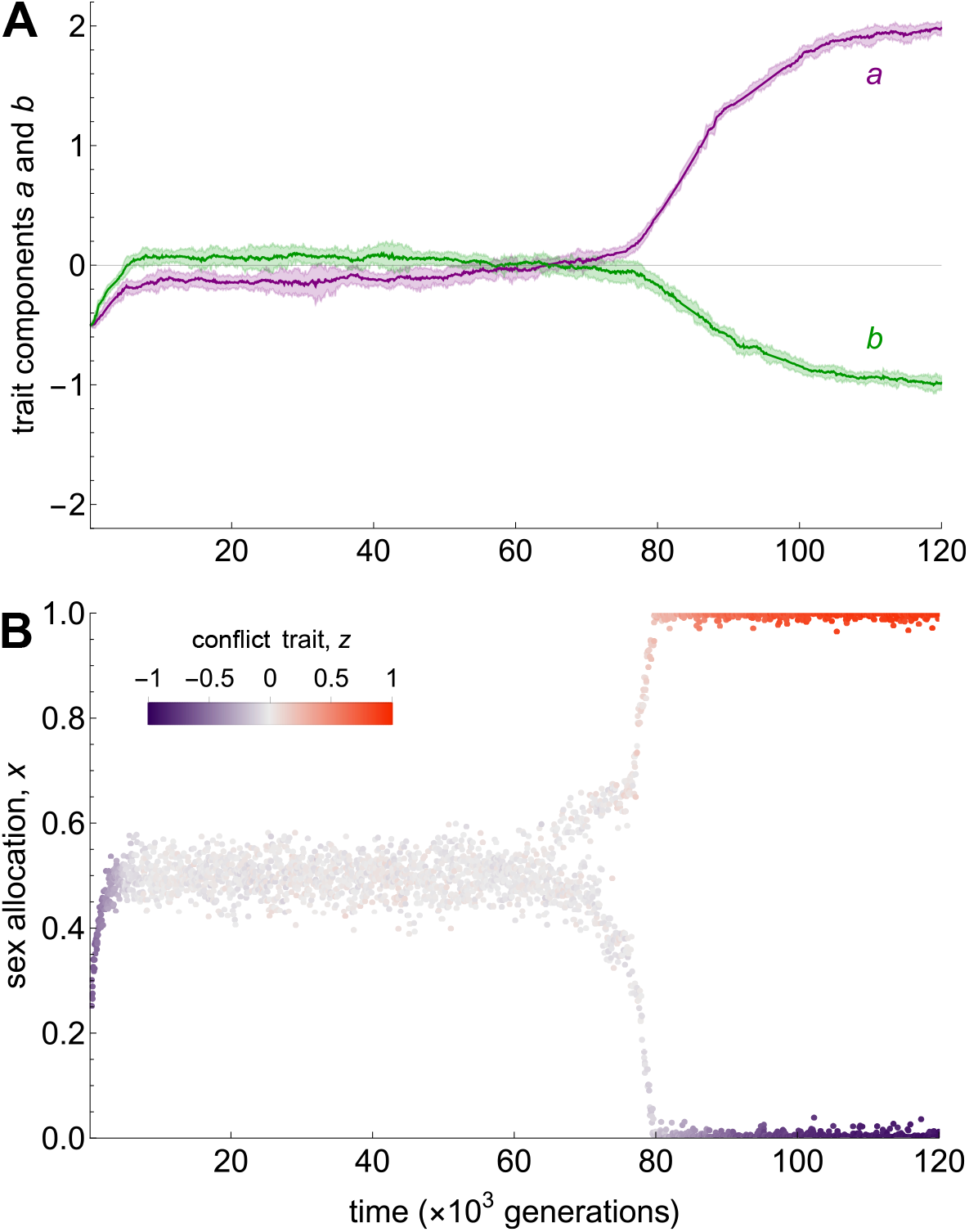
Evolution of conditional trait expression. **A** Sex allocation-dependent *a* (in purple) and -independent *b* (in green) components of trait *z* as a function of time. Solid lines indicate mean values in the population, and shaded areas indicate two standard deviations around these means. **B** Sex allocation *x* (y-axis) and trait *z* = *a*·*x* + *b* (colour scale) during the same simulation. The population first evolves to an intermediate equilibrium **y**^*^ = (*x*^*^, *a*^*^, *b*^*^) such that *x*^*^ = 1*/*2 and *a*^*^ = *b*^*^ = 0 (so that *z*^*^ = 0), and then undergoes the simultaneous emergence of conditional expression of *z* and polymorphism in sex allocation, ultimately leading to the stable coexistence of females and males expressing their sex-specific optima. Parameters used in the simulation: *θ* ♀= 1, *θ*♂ = − 1, *ω*♀ = *ω* ♂= 1*/*3, *N* = 5 *×*10^3^, *µ* = 10^−3^, *σ* = 10^−2^ for all traits (*x, a* and *b*).

The second case to consider is when

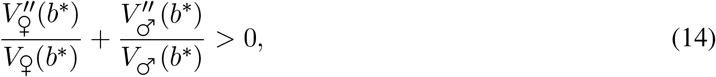

so that selection on trait components *a* and *b* is disruptive when population expresses **y**^*^ (eq. 9). Under these conditions, individual-based simulations show the joint evolution of separate sexes and sexual dimorphism occurring most of the time, with conditional trait expression given by eq. (13). However, in some simulations where 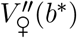 and 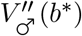 are particularly large, conditional expression fails to evolve and evolutionary branching instead occurs for *b*, the component of trait *z* that is independent of sex allocation. This leads to a hermaphroditic population where female-adapted individuals expressing *b* = *θ*♀ coexist with male-adapted individuals expression *b* = *θ*♂ (while the population remains fixed for *x*^*^ = 1*/*2 and *a*^*^ = 0). Such a population is evolutionarily stable, as there is no longer selection for conditional expression of *z*. Nevertheless, polymorphism for *b* generates selection for recombination suppression between the loci encoding sex allocation *x* and trait *b*, as seen in Section 3.1. We therefore expect that if recombination between loci influencing *x* and *b* could evolve, separate sexes would readily emerge.

### 3.3 Diploidy and the evolution of dominance

Our results above assume that individuals are haploid, which facilitates the build-up of genetic associations among traits. In diploids, such build-up may be more difficult, as an allele’s phenotypic effect is diluted in heterozygotes unless it is dominant. To see this, we extended our model to consider individuals that are diploid (haploid gametes are produced through meiosis that then fuse randomly to give rise to diploid zygotes that mature into diploid seeds). We assume that sex allocation and trait *z* are encoded by fully linked loci evolving under a ‘continuum-of-alleles’ model, as in our baseline model, and that alleles are additive at these loci, meaning that they contribute equally to the expressed phenotype. For instance, an individual carrying alleles *x*_1_ and *x*_2_ at the sex allocation locus will express *x* = (*x*_1_ + *x*_2_)*/*2.

We show in Appendix E.1.1 that the strength of directional selection on sex allocation and trait *z* is half of that found under haploidy, but it takes the population to the same equilibrium (given by eq. 3). Similarly, the Hessian matrix, which determines the conditions under which polymorphism is favoured, is equal to the one obtained under haploidy multiplied by a factor 1*/*4, so that selection for polymorphism is weaker but occurs under the same conditions for diploids and haploids. Therefore, diploidy affects the strength but not the nature of selection here (we show in Appendix E.3 that this result also holds under partial selfing, which we consider in section 4.2 below).

When polymorphism is favoured, evolutionary branching leads to the emergence of a female and a male haplotype, with alleles encoding pure femaleness (*x* = 1) and pure maleness (*x* = 0) at the sex allocation locus found in complete linkage with female- and male-beneficial alleles at the conflict trait locus, respectively. Because alleles are additive at both loci, the coexistence of these two haplotypes leads to the production of intermediate heterozygotes. These heterozygotes are unfit but remain at high frequency due to matings among homozygotes, and this situation should favour the evolution of dominance (Billiard et al., 2021). We investigated this possibility using simulations in which the sex allocation and the conflict trait locus each have two components, a gene at which alleles encode phenotypic values (as before) and a promoter sequence. The promoter is characterised by its affinity with transcription factors, which we model as a continuous trait subject to mutation. Variation in promoter affinity determines allelic expression and so encodes dominance relationships between alleles at the associated gene through *cis* regulation (Van Dooren, 1999; see Appendix E.1.2 for details).

Simulations show that dominance always evolves in this diploid model, such that the population eventually consists of males and females expressing the trait value that is optimal for their sex at equilibrium, as in our haploid model. Sex is determined either by an XY or by a ZW system of sex determination, depending on which of the male or female allele becomes dominant, respectively (Fig. S2). We also ran simulations where trait *z* could evolve conditional expression on sex allocation (as in section 3.2), and found that dominance readily evolves in this case too (Appendix E.2). Overall, we therefore observe that while diploidy diminishes the intensity of selection due to a reduced association among alleles and traits, it does not alter the evolutionary outcomes in our model, as the evolution of dominance effectively resolves the genetic constraints posed by diploidy. However, this reduced selection could present challenges in small populations, where genetic drift may become a more dominant evolutionary force.

## 4 The role of ecology in maintaining hermaphroditism

We now explore the effects of two prevalent ecological factors in flowering plants on the joint evolution of separate sexes and sexual dimorphism: limited dispersal (Section 4.1) and self-fertilisation (Section 4.2).

### 4.1 Limited dispersal and kin competition

Plants are immobile and have limited dispersal abilities. This leads to pervasive kin competition. Indeed, most pollen is dispersed over short distances, so that related pollen grains often compete with one another to fertilise neighbouring ovules (“Local Mate/Sperm Competition”; Hamilton, 1967; Schärer, 2009). Similarly, seeds are often dispersed in the close vicinity of maternal plants, so that related offspring compete with one another for recruitment (“Local Resource Competition”, Clark, 1978; Charnov, 1982). Competition among related pollen grains and related seeds increases as a focal parent allocates more heavily into its male and female functions, respectively, which in turn generates diminishing fitness returns to sexual specialisation, and is thus expected to favour hermaphroditism (Chap. 8 in Maynard Smith, 1978; Lloyd and Bawa, 1984; Fromhage and Kokko, 2010; Biernaskie, 2010). Here, we extend our baseline model (Section 2) to consider the effect of limited dispersal on the joint evolution of sex allocation and a trait under sexual conflict.

In this extended model, the population is subdivided into an effectively infinite number of patches that each carries *N* individuals, and that are uniformly connected by seed dispersal (the ‘Island Model’, Wright, 1931). At the beginning of each generation, individuals undergo sexual development, expressing sex allocation *x* and trait *z*, and then mate randomly within each patch, leading to the production of a large number of haploid seeds. A fraction *d*_s_ ∈ [0, 1] of the produced seeds is dispersed to other patches in the population, while the remaining fraction 1 − *d*_s_ stays in their natal patch. Following seed dispersal, all adults die and seeds present in each patch compete for recruitment. Due to limited seed dispersal, individuals thus compete with relatives through male function for siring ovules in their patch, and through female function for recruitment of non-dispersed seeds. To focus on the effects of kin competition on correlational selection, we assume that individuals are haploid and that sex allocation *x* and trait *z* are encoded by fully-linked quantitative trait loci evolving according to a ‘continuum-of-alleles’ model (as in our baseline model).

We show in Appendix C that the population first converges to the equilibrium **y**^*^ = (*x*^*^, *z*^*^) where

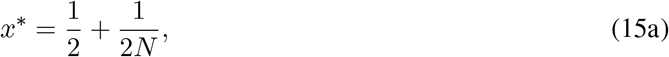

and *z*^*^ such that,

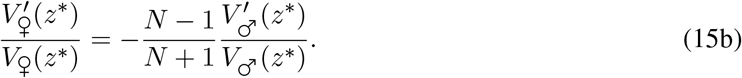

The equilibrium for sex allocation *x*^*^ corresponds exactly to Hamilton (1967)’s result for the effect of local mate competition on sex-ratio evolution in dioecious species (see also eq. 12 in Roux et al., 2024 for the diploid additive case with selfing). This allocation becomes increasingly female-biased as patch size *N* decreases, owing to an intensification of kin competition in the pollen pool (Fig. 5A). The equilibrium *z*^*^ is affected by limited dispersal in a similar way to sex allocation, with decreasing patch size selecting for more female-biased trait values (Fig. 5B). Neither *x*^*^ nor *z*^*^ depends on the seed dispersal rate *d*_s_. This is because decreasing *d*_s_ causes an increase in kin competition through male and female function that cancel out exactly (Frank, 1986; Biernaskie, 2010).

**Figure 5.**
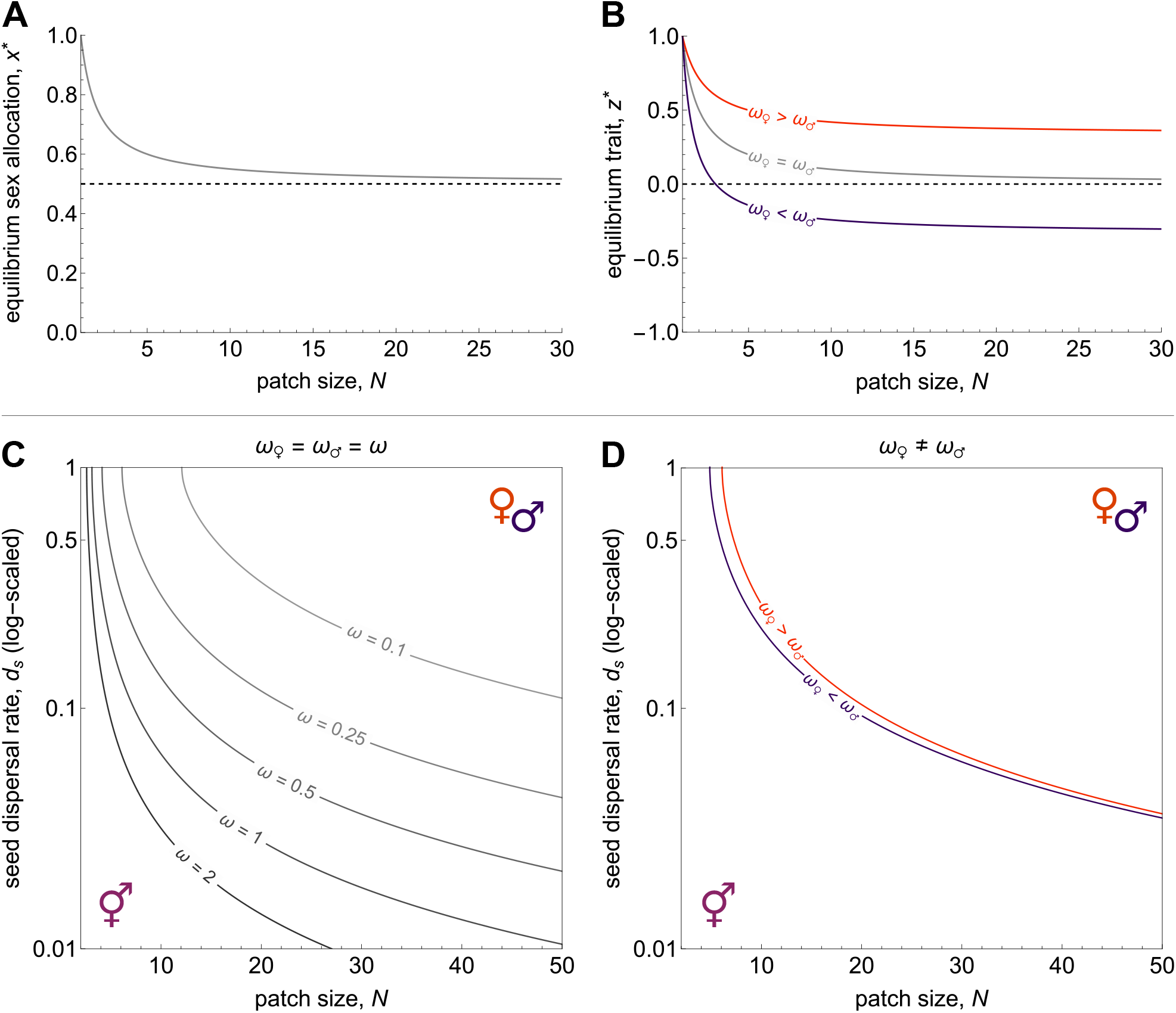
Effect of limited dispersal on the joint evolution of sex allocation *x* and trait *z*. Gaussian fecundity functions with sex-specific optima set to *θ*♀ = 1 and *θ*♂ = −1 are assumed throughout (eq. 2). **A** Equilibrium sex allocation *x*^*^ favoured by directional selection under limited dispersal (eq. 15a). The population becomes increasingly female-biased as patch size *N* decreases due to local mate competition. **B** Equilibrium trait *z*^*^ as a function of patch size *N* (eq. 15b with Gaussian functions). The grey line shows a case where selection is equally strong between sexual functions (*ω*♀ = *ω*♂ = 1*/*4). The orange line shows a case where selection is stronger through female function (*ω*♀ = 1*/*2 and *ω*♂ = 1*/*4), and the purple line shows the symmetrical case where selection is stronger through female function (*ω*♀ = 1*/*4 and *ω*♂ = 1*/*2). **C-D** Patch size *N* and seed dispersal *d*_s_ (log-scaled) leading to polymorphism when selection is equally strong through female and male function (*ω*♀ = *ω*♂ = *ω*, panel C), and when the strength of selection differs between the two sexual functions (*ω*♀ ≠ *ω*♂, panel D). For a given strength of selection *ω* polymorphism is favoured above the line (as indicated by the female and male symbols shown in the top-right corner), whereas hermaphroditism is maintained below. In panel D, the orange line shows a case where selection is twice as strong through female function (*ω*♀ = 1*/*2 and *ω*♂ = 1*/*4), and the purple line shows the opposite case where *ω*♀ = 1*/*4 and *ω*♂ = 1*/*2.

Once the population expresses **y**^*^, we find that the effect of limited dispersal on the Hessian matrix (eq. 4) depends on patch size *N* and seed dispersal *d*_s_ in a complex way (Appendix C.1.2 for derivation and Mathematica Notebook for full expression). Key insights can however be gained from considering the case of large patches connected by low seed dispersal, i.e., *N* → ∞ and *d*_s_ → 0, such that the number of immigrants *Nd*_s_ remains constant. In this case, the entries of the Hessian matrix (eq. 4) read as

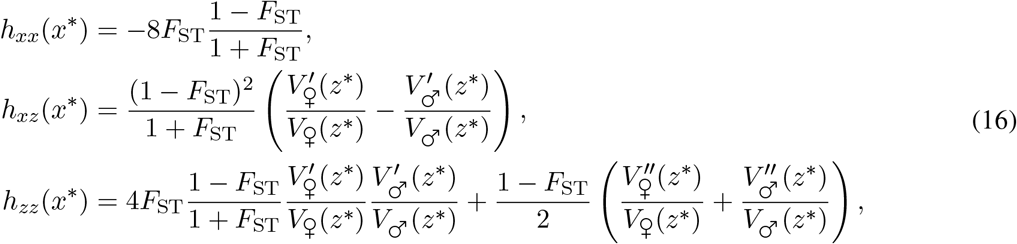

where *F*_ST_ = 1*/*(1 + 2*Nd*_s_) is Wright’s (e.g., p. 28 in Rousset, 2004), and the equilibrium **y**^*^ = (*x*^*^, *z*^*^) is identical to the well-mixed case (as eq. 15 reduces to eq. 3 when *N* → ∞). The first thing to notice from eq. (16) is that each selection coefficient is factored by 1 − *F*_ST_, reflecting the well-known effect that the intensity of selection is weaker under limited dispersal, as an individual’s reproductive success comes at the expense of genetic relatives (Frank, 1998; Rousset, 2004). Once this first effect is factored out, the second thing to notice is that quadratic selection on *x* and *z* is more negative in populations that are more structured (since 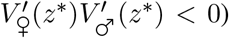. This is because kin competition among related pollen grains and among seeds leads to diminishing returns to specialising into one sexual function (also a well-known result for sex allocation *x*; Lloyd and Bawa, 1984; Biernaskie, 2010). Overall, this means that condition (5b) is harder to fulfill, indicating that limited dispersal tends to stabilise hermaphroditism against correlational selection on *x* and *z*. To see this more clearly, we can substitute eq. (16) into condition (5b) assuming Gaussian fecundity functions (eq. 2). Re-arrangements then yield the following necessary condition for correlational selection to lead to evolutionary branching

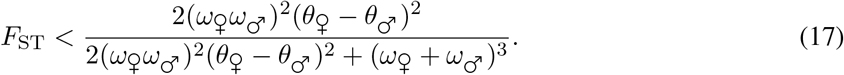

In other words, *F*_ST_ must be sufficiently low compared to the strength of sexual conflict, which here is measured by (*ω*♀ *ω*♂)^2^(*θ*♀ − *θ*♂)^2^ (i.e., the distance between the male and female optima weighted by the product of selection strength through male and female function).

We also computed numerically the eigenvalue of the Hessian matrix for arbitrary values of patch size *N* and seed dispersal *d*_s_ assuming Gaussian functions (eq. 2). We first set the intensity of selection to be equal through female and male function (*ω* = *ω*♀ = *ω*♂). In line with condition (17), we see in Figure 5C that polymorphism occurs more readily when patches are large, seed dispersal is high, and selection is strong (i.e., larger *ω*). Figure 5D also shows that increasing selection on male function is more conducive to polymorphism than increasing selection on female function (i.e., increasing *ω*♂ compared to increasing *ω* ♀). To understand this result, it is useful to first note that sexual conflict over trait *z*, which here drives sexual specialisation, is strongest when the population expresses an intermediate value of *z* that is far from both the male and female optima. All else being equal, local mate competition makes the equilibrium *z*^*^ female-biased (i.e., closer to the female optimum) when *N* is finite (recall eq. 15b and Fig. 5B). When selection is stronger over male traits (*ω*♀ *< ω*♂), this can offset the effects of local mate competition and thus bring the equilibrium *z*^*^ closer to an intermediate value where sexual conflict is stronger, and so make selection for sexual specialisation more intense.

### 4.2 Self-fertilisation and inbreeding depression

Hermaphroditic plants produce both pollen and ovules, which grants them the ability to self-fertilise. Although self-fertilisation (selfing for short) can incur a fitness cost due to inbreeding depression (Charlesworth and Charlesworth, 1987; Charlesworth and Willis, 2009), it allows hermaphrodites to produce offspring even when mating opportunities are scarce or absent (i.e., it brings “reproductive assurance”), a situation that may be particularly frequent in plants, which cannot move in search of mates (Lloyd, 1992; Kalisz et al., 2004; Busch and Delph, 2012). Moreover, selfing has a transmission advantage because a parent transmits twice as many gene copies to its selfed relative to its outcrossed progeny (Fisher, 1941). As a result, selfing yields higher fitness returns than outcrossing when inbreeding depression is weak, which further favours hermaphroditism (Charlesworth and Charlesworth, 1978b; De Jong and Geritz, 2001). In this section, we extend our baseline model (Section 2) to investigate the effect of the interplay between the transmission advantage of selfing and inbreeding depression on transitions to dioecy via correlational selection. We do not consider the benefits of reproductive assurance for simplicity.

As in our baseline model, we consider a large population of haploid hermaphrodites in which, each generation, individuals first undergo sexual development, dividing their reproductive resources between their female and male functions in proportions *x* and 1 − *x*, respectively, and expressing a trait value *z*. In contrast to our baseline model, individuals now retain a fraction *α* ∈ [0, 1] of their pollen for self-fertilisation and disperse the remainder for outcrossing, distributing it equally among other individuals in the population. The selfing rate of an individual is determined by the amount of pollen it retained relative to the total pollen it received, following a mass-action model (Holsinger, 1991). We assume throughout that enough pollen is produced for all ovules to be fertilised. This ‘competing’ mode of selfing (Lloyd, 1979) implies that the selfing rate of an individual depends on its sex allocation, with more male individuals having higher selfing rates. Mating results in the formation of a large number of outcrossed and selfed diploid zygotes. Outcrossed zygotes are viable with probability 1. Selfed zygotes suffer from inbreeding depression, so that they are viable and give rise to haploid seeds with probability 1 − δ, where δ ∈ [0, 1] measures inbreeding depression (Charlesworth and Charlesworth, 1987). Finally, all adults die, and the next generation is sampled from the produced seeds.

Again, to focus on the effects of self-fertilisation on correlational selection, we assume that the genetic basis of sex allocation *x* and trait *z* is the same as in our baseline model (Section 2).

We show in Appendix D that the population first converges to the equilibrium **y**^*^ = (*x*^*^, *z*^*^) where

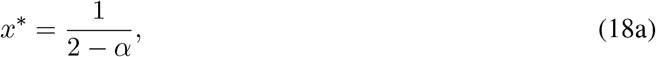

and *z*\* is such that

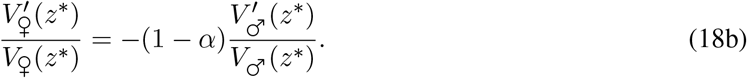

Eq. (18a) reduces to eq. (3) in the absence of selfing (*α* = 0) and increases with *α*, indicating that populations in which selfing is more common should evolve more female-biased sex allocations (Fig. 6A). This is because selfing reduces the number of ovules available for siring as an outcrossing male, which reduces opportunities to gain fitness through male function and selects for female-biased strategies (Charlesworth and Charlesworth, 1981; Lesaffre et al., 2024). For the same reason, populations where selfing is more common should also express more female-beneficial trait values, as seen from eq. (18b) and Figure 6B.

**Figure 6.**
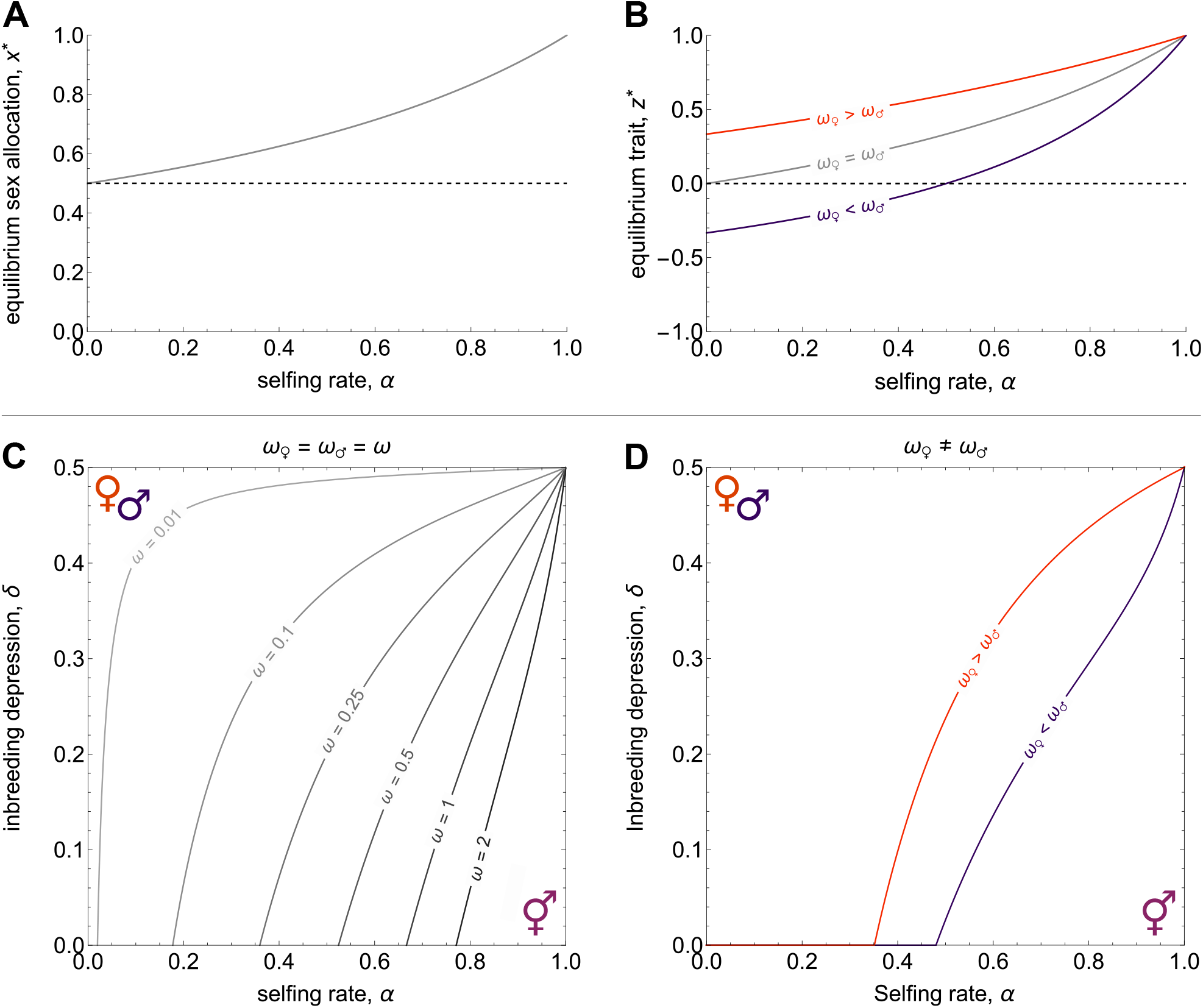
Effect of selfing on the joint evolution of sex allocation *x* and trait *z*. We assume Gaussian fecundity functions throughout, with sex-specific optima set to ♀*θ* = 1 and *θ*♂ =− 1 (eq. 2). **A** Equilibrium sex allocation *x*^*^ favoured by selection under partial selfing (eq. 18a). The population becomes increasingly female-biased as the selfing rate (*α*) increases because selfing reduces the relative contribution of male function to fitness. **B** Equilibrium trait *z*^*^ as a function of the selfing rate (eq. 18b with Gaussian functions, eq. 2). The grey line shows a case where selection is equally strong between sexual functions (*ω*♀ = *ω*♂ = 1*/*4). The orange line shows a case where selection is stronger through female function (*ω*♀ = 1*/*2 and *ω*♂ = 1*/*4), and the purple line shows the symmetrical case where selection is stronger through female function (*ω*♀ = ¼ and *ω*♂ = 1*/*2).**C-D** Selfing rate *α* and inbreeding depression δ leading to polymorphism when selection is equally strong through female and male function (*ω*♀ = *ω*♂ = *ω*, panel C), and when the strength of selection differs between the two sexual functions (*ω*♀ *≠ω*♂, panel D). For a given strength of selection *ω* polymorphism is favoured above the line (as indicated by the female and male symbols shown in the top-right corner), whereas hermaphroditism is maintained below. In panel D, the orange line shows a case where selection is twice as strong through female function (*ω*♀ = 1*/*2 and *ω*♂ = 1*/*4), and the purple line shows the opposite case where *ω*♀ = 1*/*4 and *ω*♂ = 1*/*2.

Once the population expresses **y**^*^, we show in Appendix D.2 that the entries of the Hessian matrix (eq. 4) are given by

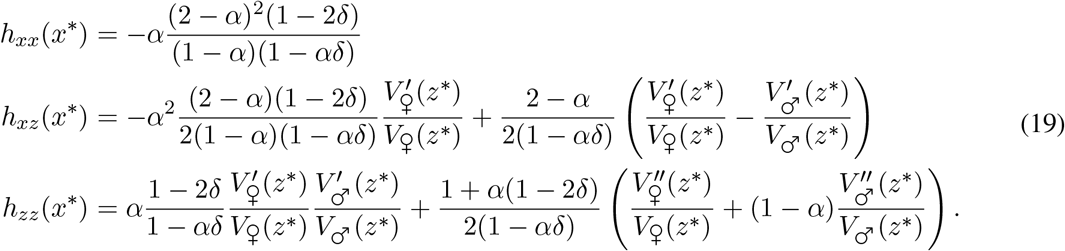

The term *h*_*xx*_(*x*^*^) in eq. (19) shows that quadratic selection on sex allocation can either promote or oppose polymorphism under partial selfing, depending on the magnitude of inbreeding depression (e.g., Charlesworth and Charlesworth, 1978b; De Jong and Geritz, 2001; Lesaffre et al., 2024). When inbreeding depression is too weak to offset the twofold transmission advantage of selfing (i.e., when outbred zygotes have less than twice the chance to survive than selfed zygotes, δ *<* 1*/*2), selfing leads *h*_*xx*_(**y**^*^) to become more negative and thus opposes the emergence of polymorphism. In contrast, when inbreeding depression is strong enough to offset the twofold transmission advantage of selfing (i.e., when δ *>* 1*/*2), selfing generates increasing fitness returns to becoming more female and thus leads quadratic selection to favour polymorphism in sex allocation (see Lesaffre et al., 2024 for further explanations).

The effects of selfing on quadratic selection on *z* and correlational selection on *x* and *z* (*h*_*zz*_(*x*^*^) and *h*_*xz*_(*x*^*^)) are not immediately clear from eq. (19), nor is the overall effect of selfing on sexual specialisation through correlational selection. To gain insights into this question, we substitute eq. (19) into condition (5b) assuming Gaussian fecundity functions, with equal strength of selection on female and male traits (eq. 2 with *ω* = *ω*♀ = *ω*♂). After some re-arrangements, we find that for polymorphism to occur due to correlational selection, the strength of selection must be greater than some threshold, specifically

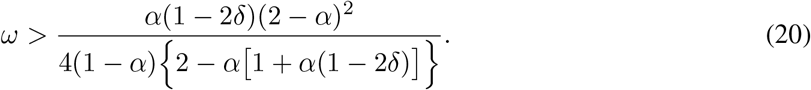

This threshold increases with the selfing rate when δ *<* 1*/*2. This result shows that, where inbreeding depression is weak enough, selfing opposes the emergence of polymorphism, which requires greater sexual conflict over *z* as *α* increases and δ decreases (i.e., higher *ω* values, Fig. 6C). We analyse the effect of asymmetrical selection through male and female function numerically in Figure 6D (i.e., *ω*♀ *> ω*♂ or *ω*♀ *< ω*♂). Similarly to what we observed under limited dispersal, an increase of selection on male traits favours polymorphism to a greater extent than an increase of selection on female traits. Again, this is because selfing favours an equilibrium *z*^*^ that is closer to the female optimum (eq. 18b, Fig. 5B), so that an increase of selection in males intensifies sexual conflict over *z* more significantly than an increase in females, thus promoting sexual specialisation.

## 5 Discussion

Sex allocation theory predicts that transitions to dioecy should occur when male and female fitness gain curves are accelerating, whereas hermaphroditism should be maintained when gain curves are saturating (Charnov et al., 1976; Charnov, 1982). Because hermaphroditism is so prevalent among flowering plants (more than 90% of species; Renner, 2014), it is generally thought that gain curves should be saturating in the absence of ecological mechanisms causing them to become accelerating (Charlesworth and Morgan, 1991; Charlesworth, 1999; Pannell and Jordan, 2022). However, previous theory has largely assumed that sex allocation evolves in isolation. Here, we have shown that the joint evolution of sex allocation with a trait that affects fitness gained through female and male function differently generates an inevitable advantage to sexual specialisation. When ecological or genetic constraints are absent, as in our baseline model, this advantage readily leads to the simultaneous emergence of separate sexes and sexual dimorphism. Given that many life-history and reproductive traits may have conflicting effects on fitness gained through female and male function (Freeman et al., 1997; Barrett, 2002b; Barrett and Hough, 2013), our results therefore suggest that fitness gain curves may in fact be accelerating in the absence of ecological mechanisms stabilising hermaphroditism. Dioecy should thus evolve whenever these mechanisms are insufficient to offset the advantage of sexual specialisation brought about by variation in traits under sexual conflict.

The advantage of sexual specialisation in our model relies on the build-up of an association between sex allocation and a trait under sexual conflict. This association is free to arise under haploidy and complete linkage between the loci encoding the two phenotypes, as in our baseline model. In that case, correlational selection generates linkage disequilibrium between female-beneficial alleles on the one hand and male-beneficial alleles on the other hand, so that the population eventually consists of sexually dimorphic females and males that are each encoded by a different haplotype. Several genetic mechanisms may interfere with the build-up of an association between sex allocation and a trait under sexual conflict. Recombination between the loci encoding the two traits can prevent polymorphism from emerging, as it breaks up the genetic associations favoured by selection (Fig. 2). However, selection readily leads to the suppression of recombination (Fig. 3), as individuals with lower recombination rates are more likely to produce offspring that express the fittest haplotypes that associate one sexual function with its optimal trait value (Fisher, 1930; Crow, 1970; Barton, 1995; Barton and Charlesworth, 1998). Furthermore, physical linkage is not the only way sex allocation and traits under sexual conflict can become associated. In fact, the associations between sex and other traits that lead to sexual dimorphism are commonly achieved through sex-biased and sex-specific gene expression, which does not rely on any physical linkage (Ellegren and Parsch, 2007; Barrett and Hough, 2013; Mank, 2017). Our model shows how correlational selection favours traits under sexual conflict to evolve conditional expression on sex allocation, which in turn promotes the emergence of separate sexes (Fig. 4). This suggests that sexual dimorphism resulting from differences in gene regulation can easily begin to evolve in hermaphrodites, and in turn fuel the transition to dioecy.

Our baseline model assumes haploid individuals with a reduced diploid phase. Sex expression in the haploid phase of the life cycle is common in many haplontic organisms such as algae, mosses or ferns, and transitions between combined and separate sexes have occurred many times in some of these groups (Villarreal and Renner, 2013; Haig, 2016; Coelho et al., 2018; Charlesworth, 2022). Our model describes a possible path along which such evolutionary transitions may occur, and also suggests how they might unfold at the genetic level. Indeed, correlational selection leads to the gradual divergence of female- and male-coding haplotypes in our haploid model, which form the basis of UV sex chromosomes, a common system of sex determination in haplontic species (Coelho et al., 2018).

Although our baseline model considers haploid individuals, our results readily apply to the many transitions to dioecy in diploid flowering plants. We investigated how diploidy affects our main results, and found correlational selection favours dioecy under identical conditions irrespective of ploidy in our baseline model and under partial selfing. Although we did not model the stabilising effect of limited dispersal on hermaphroditism under diploidy, we expect results would be qualitatively similar to the haploid case, with quantitative differences due to the influence of ploidy on relatedness (Rousset, 2004). The main effect of diploidy is seen once polymorphism has emerged, as it leads to the production of heterozygotes expressing intermediate trait values under additivity. We showed that the production of unfit heterozygotes favours the evolution of dominance, which lifts the genetic constraint posed by diploidy, both in our baseline model and when the conflict trait evolved conditional expression on sex allocation. Dominance evolution at the sex allocation locus leads to the emergence of either an XY or a ZW sex determination system, depending on whether the male or female allele becomes dominant, respectively, as seen in most dioecious plants (Bachtrog et al., 2014; Lesaffre et al., 2024). We did not consider the evolution of recombination in diploids. Presumably, diploidy would weaken selection on recombination but it would not change its direction. This is because haplotypes carrying matching alleles would still be involved in fitter genotypes than those carrying mismatched alleles when averaged over genetic backgrounds. There would thus still be positive epistasis between loci, favouring recombination suppression (Barton, 1995). Moreover, our results on conditional expression evolution demonstrate that physical linkage between the loci encoding sex allocation and the conflict trait is not necessary for correlational selection to act in diploids. Overall, our results thus indicate that diploidy should not impede the joint emergence of separate sexes and sexual dimorphism from hermaphroditism in the long-term, consistent with the numerous such transitions that have occurred in diploid organisms.

A potential genetic constraint that we did not consider is polygenicity, which could also hamper the joint evolution of separate sexes and sexual dimorphism, as the expression of alleles segregating at many loci would prevent the production of discrete, well-differentiated sexual morphs. However, selection for discrete, highly-differentiated morphs on a trait with an initially polygenic basis is expected to favour the concentration of its genetic basis, such that it ends up being encoded by one or few loci (because such concentration allows greater heritability of the discrete phenotypes favoured by selection; van Doorn and Dieckmann, 2006; Kopp and Hermisson, 2006; Yeaman and Whitlock, 2011). Indeed, Lesaffre et al. (2024) showed that when sex allocation evolves alone, disruptive selection for dioecy favours the concentration of an initially polygenic basis of sex to a single locus at which complete dominance evolves, resulting in the emergence of nascent sex chromosomes.

When sex allocation evolves jointly with a trait under sexual conflict, we might expect the genetic architecture of sex determination to depend on the way sex allocation and the other trait become genetically associated. If the association comes about from sex-dependent expression (e.g., via gene regulation), then the concentration of the genetic basis of sex should unfold in a similar way to when sex allocation evolves alone (Lesaffre et al., 2024), with the genetic basis of the trait under sexual conflict remaining polygenic. The resulting sex-determining region could therefore contain as little as a single gene, and so initially show no sign of recombination suppression around the sex-determining locus. In contrast, if sex allocation and the trait under sexual conflict become associated through recombination suppression, then the resulting sex-determining region might consist of a non-recombining chromosomal segment capturing the loci encoding sex and (at least) one conflict trait, which could constitute a first step towards the extension of recombination suppression along sex chromosomes (Rice, 1984). This recombination suppression scenario shows interesting parallels with the scenario proposed in Muralidhar and Veller (2018) for evolutionary transitions from environmental sex determination, where sex is determined by abiotic conditions via a threshold trait with a typically polygenic basis, to genetic sex determination, where sex is determined by an individual’s genotype at a single Mendelian locus. Muralidhar and Veller (2018) showed that alleles modifying the threshold such that their carriers are more likely to develop into a given sex can increase in frequency when linked with a sexually antagonistic mutation benefiting that sex. The establishment of such an allelic combination in turn favours alleles elsewhere in the genome that bias the threshold back in the other direction as a result of sex-ratio selection. Through the sequential spread and capture of such allelic combinations into an increasingly large non-recombining haplotype, carriers of this haplotype eventually end up developing exclusively into one sex, and non-carriers exclusively into the other, so that sex determination has effectively become genetic and encoded by a large non-recombining region. In the context of our model, a similar scenario could unfold if sex allocation and the conflict trait initially have a polygenic basis, such that alleles encoding a sex allocation biased towards a given sex gradually become linked with alleles beneficial to that sex, leading to the emergence of a sex-determining haplotype.

Our results indicate that although genetic constraints can hinder the joint emergence of separate sexes and sexual dimorphism in the short-term, they are unlikely to prevent it in the long-term. The maintenance of hermaphroditism may thus be better explained by ecological benefits of hermaphroditism, which would have to offset the selective advantage of sexual specialisation brought about by jointly evolving traits. Here we have explored the effects of limited dispersal, which characterises many flowering plant species (Vekemans and Hardy, 2004). Our analyses show that kin competition caused by limited dispersal stabilises hermaphroditism against correlational selection for dioecy and sexual dimorphism, because local mate competition (Hamilton, 1967) and local resource competition (Clark, 1978) generate diminishing returns to sexual specialisation (Fig. 5). We also considered the effect of partial selfing, another common feature of hermaphroditic plants (Goodwillie et al., 2005), and found that selfing can stabilise hermaphroditism as well when inbreeding depression is weak relative to the inherent transmission advantage of selfing (Fig. 6). These results are in line with previous theoretical work that assumed that sex allocation evolves alone (e.g., Charlesworth and Charlesworth, 1978b; Lloyd and Bawa, 1984; De Jong and Geritz, 2001; Biernaskie, 2010). They imply that ecology can readily explain the maintenance of hermaphroditism in spite of inevitable correlational selection for sexual specialisation. In this context, it is interesting to note that dioecy appears to be more common in species in which the ecological benefits associated with hermaphroditism are expected to be weakest, such as long-lived plant species (Friedman, 2020), in which inbreeding depression is typically high and selfing rates low (Duminil et al., 2009; Angeloni et al., 2011; Munoz et al., 2016), as well as in species with effective dispersal mechanisms, such as wind-pollination and seed dispersal by animals (Bawa, 1980; Renner and Ricklefs, 1995).

## Supporting information

Supplementary Mathematica notebooks

## Acknowledgements

The authors would like to thank the Swiss National Science Foundation for funding (SNF grants 310030 215135 to JRP and PCEFP3181243 to CM).

