## Appendix for "The joint evolution of separate sexes and sexual dimorphism"

**A.3 Individual-based simulations** **11**

**B Evolution of phenotypic associations via the evolution of genetic architecture** 13

**B.1 Evolution of recombination suppression** 13

**D The effects of self-fertilisation and inbreeding depression** 37

**D.1 Directional selection** 38

**D.2 Disruptive selection** 39

**E Diploidy and the evolution of dominance** **40**

**E.1 Baseline model under diploidy** **40**

#### Appendix A

### Analysis of the baseline model

In this Appendix, we study the joint evolution of sex allocation $x$ and trait $z$ in the baseline model described in Section 2.1 of the main text, and derive the results presented in Section 2.2.

#### A.1 Invasion analysis

We study the evolutionary dynamics of $x$ and $z$ using invasion analysis. As a basis for this analysis, we characterise the invasion fitness $W(\mathbf{y}_\mathrm{m}, \mathbf{y})$ of a rare mutant expressing $\mathbf{y}_\mathrm{m} = (x_\mathrm{m}, z_\mathrm{m})$ in a resident population otherwise fixed for $\mathbf{y} = (x, z)$, i.e., the number of successful mutant offspring contributed by the focal mutant to the next generation. This is given by

$$\begin{aligned} W(\mathbf{y}_\mathrm{m}, \mathbf{y}) &= \frac{1}{F(\mathbf{y})}\left(\frac{1}{2}F(\mathbf{y}_\mathrm{m}) + \frac{1}{2}F(\mathbf{y})\frac{M(\mathbf{y}_\mathrm{m})}{M(\mathbf{y})}\right) \\ &= \frac{1}{2}\left(\frac{F(\mathbf{y}_\mathrm{m})}{F(\mathbf{y})} + \frac{M(\mathbf{y}_\mathrm{m})}{M(\mathbf{y})}\right) = \frac{1}{2}\left(\frac{x_\mathrm{m} V_{♀}(z_\mathrm{m})}{x\, V_{♀}(z)} + \frac{(1-x_\mathrm{m})V_{♂}(z_\mathrm{m})}{(1-x)V_{♂}(z)}\right). \end{aligned} \tag{A1}$$

Equation (A1) can be understood as follows. The term between brackets on the first line gives the number of mutant seeds produced by the focal mutant. A mutant individual produces $F(\mathbf{y}_\mathrm{m})$ ovules, which are all fertilised by resident pollen because individuals are strictly outcrossing and the mutant is rare, so that mating among mutants can be neglected. This generates diploid zygotes that are heterozygous for the mutant haplotype, so that half the seeds they produce will carry the mutant haplotype (hence $F(\mathbf{y}_\mathrm{m})$ is multiplied by $1/2$). The mutant can also transmit its genes by siring resident ovules ($F(\mathbf{y})$), which it does in proportion $M(\mathbf{y}_\mathrm{m})/M(\mathbf{y})$ assuming the population is large ($N \to \infty$). This also leads to zygotes that are heterozygous for the mutant haplotype, and therefore also to half of the produced seeds being mutant. The mutant seeds then compete for recruitment with all the seeds produced in the population, and succeed with probability $1/F(\mathbf{y})$.

##### A.1.1 Directional selection

When mutations are rare with small effects, the population first evolves gradually under directional selection, while remaining largely monomorphic (Geritz et al., 1998). It may then attain a ‘convergence stable strategy’, which is an attractor of directional selection. The strength and direction of selection is given by the selection gradients on traits $x$ and $z$, $s_x(\mathbf{y})$ and $s_z(\mathbf{y})$, which are defined as

$$s_x(\mathbf{y}) = \left.\frac{\partial W(\mathbf{y}_\mathrm{m}, \mathbf{y})}{\partial x_\mathrm{m}}\right|_{\mathbf{y}_\mathrm{m}=\mathbf{y}} \quad \text{and} \quad s_z(\mathbf{y}) = \left.\frac{\partial W(\mathbf{y}_\mathrm{m}, \mathbf{y})}{\partial z_\mathrm{m}}\right|_{\mathbf{y}_\mathrm{m}=\mathbf{y}}. \tag{A2}$$

Candidate attractors of directional selection are singular strategies $\mathbf{y}^* = (x^*, z^*)$ at which directional selection vanishes, i.e.,

$$\mathbf{y}^* = (x^*, z^*) \quad \text{such that} \quad s_x(\mathbf{y}^*) = s_z(\mathbf{y}^*) = 0. \tag{A3}$$

Using eq. (A2), we find that selection gradients $s_x(\mathbf{y})$ and $s_z(\mathbf{y})$ are given by

$$s_x(\mathbf{y}) = \frac{1-2x}{2x(1-x)} \quad \text{and} \quad s_z(\mathbf{y}) = \frac{1}{2}\left(\frac{V'_♀(z)}{V_♀(z)} + \frac{V'_♂(z)}{V_♂(z)}\right). \tag{A4}$$

**Equilibrium phenotype.** Using eq. (A4), we can solve eq. (A3) to find that the selection gradient on $s_x(\mathbf{y})$ cancels for a unique sex allocation

$$x^* = \frac{1}{2}, \tag{A5a}$$

so that selection favours equal allocation to male and female function in this baseline model. The equilibrium trait value $z^*$, meanwhile, must satisfy

$$\frac{V'_♀(z^*)}{V_♀(z^*)} = -\frac{V'_♂(z^*)}{V_♂(z^*)}, \tag{A5b}$$

i.e., it must be such that the marginal fitness gain from expressing a more female-favorable trait value is exactly compensated by the marginal fitness loss it causes through male function (and vice-versa). Eq. (A5) corresponds to eq. (3) in the main text. With Gaussian $V_u(z)$ functions (eq. 2), this becomes

$$z^* = \theta_♀\frac{\omega_♀}{\omega_♂ + \omega_♀} + \theta_♂\frac{\omega_♂}{\omega_♂ + \omega_♀}. \tag{A6}$$

Eq. (A6) shows that the singular trait value assuming Gaussian functions lies somewhere between the male and female optima, $\theta_{♂}$ and $\theta_{♀}$. When selection is equally intense in both sexual functions (i.e., when $\omega_{♀} = \omega_{♂}$), the singular value $z^*$ is equidistant from $\theta_{♂}$ and $\theta_{♀}$, i.e.,

$$z^* = \frac{\theta_{♀} + \theta_{♂}}{2}. \tag{A7}$$

Otherwise, $z^*$ is biased towards the sexual function in which selection is the strongest, i.e., where $\omega_u$ is the largest, so that it is male-biased when $\omega_{♂} > \omega_{♀}$ and female-biased when $\omega_{♂} < \omega_{♀}$.

**Convergence stability.** To be an attractor of directional selection, the singular strategy $\mathbf{y}^* = (x^*, z^*)$ must be *convergence stable*, which requires that all eigenvalue of the Jacobian matrix $\mathbf{J}(\mathbf{y}^*)$ have negative real parts at this point provided that both traits mutate independently (Leimar, 2009). This matrix is given by

$$\mathbf{J}(\mathbf{y}^*) = \begin{pmatrix} \left.\dfrac{\partial s_x(\mathbf{y})}{\partial x}\right|_{\mathbf{y}=\mathbf{y}^*} & \left.\dfrac{\partial s_x(\mathbf{y})}{\partial z}\right|_{\mathbf{y}=\mathbf{y}^*} \\ \left.\dfrac{\partial s_z(\mathbf{y})}{\partial x}\right|_{\mathbf{y}=\mathbf{y}^*} & \left.\dfrac{\partial s_z(\mathbf{y})}{\partial z}\right|_{\mathbf{y}=\mathbf{y}^*} \end{pmatrix} = \begin{pmatrix} -4 & 0 \\ 0 & \dfrac{1}{2}\left(\dfrac{V''_{♀}(z^*)}{V_{♀}(z^*)} + \dfrac{V''_{♂}(z^*)}{V_{♂}(z^*)} + 2\dfrac{V'_{♀}(z^*)}{V_{♀}(z^*)}\dfrac{V'_{♂}(z^*)}{V_{♂}(z^*)}\right) \end{pmatrix}, \tag{A8}$$

such that $\mathbf{y}^* = (x^*, z^*)$ is convergence stable when

$$\frac{1}{2}\left(\frac{V''_{♀}(z^*)}{V_{♀}(z^*)} + \frac{V''_{♂}(z^*)}{V_{♂}(z^*)} + 2\frac{V'_{♀}(z^*)}{V_{♀}(z^*)}\frac{V'_{♂}(z^*)}{V_{♂}(z^*)}\right) < 0, \tag{A9}$$

which means that functions $V_u(z)$ must not accelerate too strongly relative to their rate of increase in the vicinity of $z^*$. This is always the case with Gaussian functions (eq. 2), as this becomes

$$-\left(\omega_{♂} + \omega_{♀}\right) < 0, \tag{A10}$$

which always holds.

More broadly, condition (A9) should always be satisfied under the assumptions we made on the shapes of functions $V_u(z)$, $u \in \{♀, ♂\}$. Our assumptions that the functions are smooth with a single local and global maximum each guarantee that the equilibrium trait $z^*$ *(i)* lies somewhere between their two maxima, *(ii)* is unique and *(iii)* always convergence stable. This is because these assumptions entail that functions $V_u(z)$ are both increasing functions of $z$ when $z$ is smaller than both maxima, and both decreasing functions of $z$ above both maxima, i.e.

$$\lim_{z\to-\infty} V_u'(z) > 0 \quad \text{and,} \quad \lim_{z\to+\infty} V_u'(z) < 0, \tag{A11}$$

so that eq. (A5b), which defines the singular strategy $z^*$ can only be satisfied between the two maxima, where the derivatives of $V_u(z)$ functions have opposite signs (which shows that *(i)* must hold). Furthermore, the derivatives of $V_u(z)$ functions are monotonous between their respective optima, as $V_{♂}(z)$ always decreases as $z$ increases beyond its optimum and $V_{♀}(z)$ always increases as $z$ increases towards its optimum. This implies that the derivatives cross only once, and thus that *(ii)* $z^*$ is unique. Thus, the selection gradient cancels at $z^*$ only, is positive below and negative above (eq. A11), and so $z^*$ must be *(iii)* convergence stable.

##### A.1.2 Disruptive selection

Once the population expresses the equilibrium $\mathbf{y}^*$, it may experience stabilising selection and remain monomorphic for this $\mathbf{y}^*$, or it may experience negative frequency-dependent disruptive selection ('disruptive selection' for short) and become polymorphic via evolutionary branching (Geritz et al., 1998). Which of these two outcomes unfolds is dictated by the sign of the eigenvalues of the Hessian matrix $\mathbf{H}(\mathbf{y}^*)$. Specifically, $\mathbf{y}^*$ is evolutionarily stable if all the eigenvalues of $\mathbf{H}(\mathbf{y}^*)$ are negative, whereas polymorphism emerges if just one eigenvalue is positive. The $\mathbf{H}(\mathbf{y}^*)$ matrix is defined as

$$\mathbf{H}(\mathbf{y}^*) = \begin{pmatrix} h_{xx}(\mathbf{y}^*) & h_{xz}(\mathbf{y}^*) \\ h_{xz}(\mathbf{y}^*) & h_{zz}(\mathbf{y}^*) \end{pmatrix} = \begin{pmatrix} \left.\dfrac{\partial^2 W(\mathbf{y}_\mathrm{m},\mathbf{y})}{\partial {x_\mathrm{m}}^2}\right|_{\mathbf{y}_\mathrm{m}=\mathbf{y}=\mathbf{y}^*} & \left.\dfrac{\partial^2 W(\mathbf{y}_\mathrm{m},\mathbf{y})}{\partial x_\mathrm{m} z_\mathrm{m}}\right|_{\mathbf{y}_\mathrm{m}=\mathbf{y}=\mathbf{y}^*} \\ \left.\dfrac{\partial^2 W(\mathbf{y}_\mathrm{m},\mathbf{y})}{\partial x_\mathrm{m} z_\mathrm{m}}\right|_{\mathbf{y}_\mathrm{m}=\mathbf{y}=\mathbf{y}^*} & \left.\dfrac{\partial^2 W(\mathbf{y}_\mathrm{m},\mathbf{y})}{\partial {z_\mathrm{m}}^2}\right|_{\mathbf{y}_\mathrm{m}=\mathbf{y}=\mathbf{y}^*} \end{pmatrix}. \tag{A12}$$

In our model, this yields

$$\mathbf{H}(\mathbf{y}^*) = \begin{pmatrix} 0 & \dfrac{V_{♀}'(z^*)}{V_{♀}(z^*)} - \dfrac{V_{♂}'(z^*)}{V_{♂}(z^*)} \\ \dfrac{V_{♀}'(z^*)}{V_{♀}(z^*)} - \dfrac{V_{♂}'(z^*)}{V_{♂}(z^*)} & \dfrac{1}{2}\left(\dfrac{V_{♀}''(z^*)}{V_{♀}(z^*)} + \dfrac{V_{♂}''(z^*)}{V_{♂}(z^*)}\right). \end{pmatrix}, \tag{A13}$$

which corresponds to eq. (4) in the main text.

#### A.2 Stable polymorphism

In the main text, we show that condition (5) is always satisfied, such that polymorphism is always favoured by selection once the population converges to the equilibrium $\mathbf{y}^*$. To study the nature of the polymorphism maintained in the long run, we consider a population in which two types of individuals (denoted type 1 and type 2) expressing phenotypes $\mathbf{y}_1 = (x_1, z_1)$ and $\mathbf{y}_2 = (x_2, z_2)$, respectively, coexist. Our aim is to compute the selection gradients acting on their traits, $x_1$, $x_2$, $z_1$ and $z_2$, which we denote as $s_x^1(\mathbf{y}_1, \mathbf{y}_2)$, $s_x^2(\mathbf{y}_1, \mathbf{y}_2)$, $s_z^1(\mathbf{y}_1, \mathbf{y}_2)$ and $s_z^2(\mathbf{y}_1, \mathbf{y}_2)$, to determine the polymorphic equilibrium favoured by selection. To this end, we consider the invasion of a rare mutant expressing $\mathbf{y}_\mathrm{m} = (x_\mathrm{m}, z_\mathrm{m})$ in a resident population where individuals expressing $\mathbf{y}_1 = (x_1, z_1)$ and $\mathbf{y}_2 = (x_2, z_2)$ coexist at an equilibrium.

##### A.2.1 Equilibrium in the resident population

We first characterise the equilibrium reached by the resident population. Since the population is assumed to be of constant size $N$, this equilibrium can be simply characterised by the equilibrium frequency of type 1, $f_1^*$, as the frequency of type 2 is then given by $1 - f_1^*$. Given a frequency $f_1^t$ at time $t$, the frequency of type 1 in the next generation is given by

$$\begin{aligned} f_1^{t+1} =& \frac{f_1^t F(\mathbf{y}_1)}{f_1^t F(\mathbf{y}_1) + (1 - f_1^t) F(\mathbf{y}_2)} \frac{f_1^t M(\mathbf{y}_1) + \dfrac{1 - f_1^t}{2} M(\mathbf{y}_2)}{f_1^t M(\mathbf{y}_1) + (1 - f_1^t) M(\mathbf{y}_2)} \\ &+ \frac{(1 - f_1^t) F(\mathbf{y}_2)}{f_1^t F(\mathbf{y}_1) + (1 - f_1^t) F(\mathbf{y}_2)} \frac{\dfrac{f_1^t}{2} M(\mathbf{y}_1)}{f_1^t M(\mathbf{y}_1) + (1 - f_1^t) M(\mathbf{y}_2)}, \end{aligned} \tag{A14}$$

and the equilibrium frequency is defined as

$$f_1^* \quad \text{such that} \quad f_1^{t+1} = f_1^t. \tag{A15}$$

Solving for $f_1^*$, we obtain

$$f_1^* = \frac{F(\mathbf{y}_1) M(\mathbf{y}_2) + F(\mathbf{y}_2) M(\mathbf{y}_1)}{F(\mathbf{y}_1) M(\mathbf{y}_2) + F(\mathbf{y}_2) M(\mathbf{y}_1) - 2 F(\mathbf{y}_1) M(\mathbf{y}_1)}. \tag{A16}$$

##### A.2.2 Invasion fitness and selection gradients

We now study the fate of a rare mutant $\mathbf{y}_\mathrm{m}$ in the resident population at equilibrium. The invasion fitness of this mutant, $W_\mathrm{P}(\mathbf{y}_\mathrm{m}|\mathbf{y}_1, \mathbf{y}_2)$, where the P subscript stands for 'Polymorphic', is obtained following the same logic as for eq. (A1), and is given by

$$W_\mathrm{P}(\mathbf{y}_\mathrm{m}|\mathbf{y}_1, \mathbf{y}_2) = \frac{1}{2}\left(\frac{F(\mathbf{y}_\mathrm{m})}{\overline{F}(\mathbf{y}_1, \mathbf{y}_2)} + \frac{M(\mathbf{y}_\mathrm{m})}{\overline{M}(\mathbf{y}_1, \mathbf{y}_2)}\right), \tag{A17}$$

where

$$\overline{F}(\mathbf{y}_1, \mathbf{y}_2) = f_1^* F(\mathbf{y}_1) + (1 - f_1^*)F(\mathbf{y}_2) \quad \text{and} \quad \overline{M}(\mathbf{y}_1, \mathbf{y}_2) = f_1^* M(\mathbf{y}_1) + (1 - f_1^*)M(\mathbf{y}_2) \tag{A18}$$

are the average number of female and male gametes produced in the population.

Using eq. (A17), the selection gradients on traits $x_1$, $x_2$, $z_1$ and $z_2$ can be computed as

$$\mathbf{S}(\mathbf{y}_1, \mathbf{y}_2) = \begin{pmatrix} s_x^1(\mathbf{y}_1, \mathbf{y}_2) \\ s_x^2(\mathbf{y}_1, \mathbf{y}_2) \\ s_z^1(\mathbf{y}_1, \mathbf{y}_2) \\ s_z^2(\mathbf{y}_1, \mathbf{y}_2) \end{pmatrix} = \begin{pmatrix} \left.\dfrac{\partial W_\mathrm{P}(\mathbf{y}_\mathrm{m}|\mathbf{y}_1, \mathbf{y}_2)}{\partial x_\mathrm{m}}\right|_{\mathbf{y}_\mathrm{m}=\mathbf{y}_1} \\ \left.\dfrac{\partial W_\mathrm{P}(\mathbf{y}_\mathrm{m}|\mathbf{y}_1, \mathbf{y}_2)}{\partial x_\mathrm{m}}\right|_{\mathbf{y}_\mathrm{m}=\mathbf{y}_2} \\ \left.\dfrac{\partial W_\mathrm{P}(\mathbf{y}_\mathrm{m}|\mathbf{y}_1, \mathbf{y}_2)}{\partial z_\mathrm{m}}\right|_{\mathbf{y}_\mathrm{m}=\mathbf{y}_1} \\ \left.\dfrac{\partial W_\mathrm{P}(\mathbf{y}_\mathrm{m}|\mathbf{y}_1, \mathbf{y}_2)}{\partial z_\mathrm{m}}\right|_{\mathbf{y}_\mathrm{m}=\mathbf{y}_2} \end{pmatrix}, \tag{A19}$$

which gives

$$\mathbf{S}(\mathbf{y}_1, \mathbf{y}_2) = \begin{pmatrix} \dfrac{1}{2}\left(\dfrac{F_x(\mathbf{y}_1)}{\overline{F}(\mathbf{y}_1, \mathbf{y}_2)} + \dfrac{M_x(\mathbf{y}_1)}{\overline{M}(\mathbf{y}_1, \mathbf{y}_2)}\right) \\ \dfrac{1}{2}\left(\dfrac{F_x(\mathbf{y}_2)}{\overline{F}(\mathbf{y}_1, \mathbf{y}_2)} + \dfrac{M_x(\mathbf{y}_2)}{\overline{M}(\mathbf{y}_1, \mathbf{y}_2)}\right) \\ \dfrac{1}{2}\left(\dfrac{F_z(\mathbf{y}_1)}{\overline{F}(\mathbf{y}_1, \mathbf{y}_2)} + \dfrac{M_z(\mathbf{y}_1)}{\overline{M}(\mathbf{y}_1, \mathbf{y}_2)}\right) \\ \dfrac{1}{2}\left(\dfrac{F_z(\mathbf{y}_2)}{\overline{F}(\mathbf{y}_1, \mathbf{y}_2)} + \dfrac{M_z(\mathbf{y}_2)}{\overline{M}(\mathbf{y}_1, \mathbf{y}_2)}\right) \end{pmatrix}, \tag{A20}$$

where subscripts $x$ and $z$ denote partial differentiation of the function with respect to $x$ and $z$, respectively, so that we have for instance

$$F_x(\mathbf{y}_i) = \left.\frac{\partial F(\mathbf{y}_\mathrm{m})}{\partial x_\mathrm{m}}\right|_{\mathbf{y}_\mathrm{m}=\mathbf{y}_i} \quad \text{or} \quad M_z(\mathbf{y}_i) = \left.\frac{\partial F(\mathbf{y}_\mathrm{m})}{\partial z_\mathrm{m}}\right|_{\mathbf{y}_\mathrm{m}=\mathbf{y}_i}, \tag{A21}$$

with $i \in \{1, 2\}$.

##### A.2.3 Numerical analysis

**Algorithm.** The selection gradients given in eq. (A20) can be used to infer the evolutionary dynamics of the population numerically. Let us define vector $\mathbf{Y}_t$ whose entries are the phenotypes encoded by the two haplotypes at some time-step $t$, i.e.,

$$\mathbf{Y}_t = \begin{pmatrix} x_1^t \\ x_2^t \\ z_1^t \\ z_2^t \end{pmatrix}. \tag{A22}$$

Assuming mutations of weak and independent effects on each of the four traits, the change in phenotypes over evolutionary time can be computed using the recursion

$$\mathbf{Y}_{t+1} = \mathbf{Y}_t + \kappa\, \mathbf{S}(\mathbf{y}_1^t, \mathbf{y}_2^t), \tag{A23}$$

where $\kappa > 0$ is a small constant. From an initial point $\mathbf{Y}_0$ close to the singular strategy $\mathbf{y}^*$, i.e.,

$$\mathbf{Y}_0 = \begin{pmatrix} x^* + \epsilon \\ x^* - \epsilon \\ z^* + \epsilon \\ z^* - \epsilon \end{pmatrix}, \tag{A24}$$

with $\epsilon > 0$ a small deviation, we compute the vector of phenotypes in the next time-step using eq. (A23), and correct the obtained values in order to keep all traits within their definition domain (namely, we correct $x_1$ and $x_2$ such that they remain in $[0, 1]$). We iterate this algorithm until we reach an equilibrium combination of traits

$$\mathbf{Y}^* = \begin{pmatrix} x_1^* \\ x_2^* \\ z_1^* \\ z_2^* \end{pmatrix}, \quad \text{such that} \quad \mathbf{Y}_{t+1} = \mathbf{Y}_t. \tag{A25}$$

**Results.** Repeating the analysis described above for many combinations of parameters assuming Gaussian functions (eq. 2), we find that selection always leads to the same polymorphic equilibrium, with one type allocating all its resource to female function ($x_1^* = 1$) and expressing the female optimal trait value, i.e., $\mathbf{y}_1^* = (1,\ \theta_{♀})$, and the other allocating all its resource to male function ($x_2^* = 0$) and expressing the male optimal $z$ value, $\mathbf{y}_2^* = (0,\ \theta_{♂})$, so that we have

$$\mathbf{Y}^* = \begin{pmatrix} 1 \\ 0 \\ \theta_{♀} \\ \theta_{♂} \end{pmatrix}. \tag{A26}$$

#### A.3 Individual-based simulations

To complement our numerical analysis and validate our findings, we ran individual-based simulations of our baseline model. The simulation program is coded in C++.

It simulates a population of haploid individuals with a constant size $N$, in which the $i^{\text{th}}$ individual is characterised by its genotype at the sex allocation and trait locus $x_i$ and $z_i$. The population is initially fixed for arbitrary sex allocation and trait values $x_0 \in (0, 1)$ and $z_0 \in \mathbb{R}$. Each generation proceeds as follows. We first determine the female and male fecundities of individuals. For individual $i$, these are given by

$$F(x_i, z_i) = x_i V_{♀}(z_i) \quad \text{and} \quad M(x_i, z_i) = (1 - x_i) V_{♂}(z_i)\,, \tag{A27}$$

respectively, where we use Gaussian functions for the effect of $z$ (eq. 2). To create the next generation, we generate $N$ haploid offspring by directly sampling $N$ parents with replacement from the previous generation, which we can do because neither recombination nor selection occurs during the diploid phase (Section 2.1). Parents are sampled from the female or male gamete pool with equal probability (i.e., each haploid offspring is equally likely to carry maternally or paternally inherited alleles). Within each sex, parents are sampled in proportion to their sex-specific fecundity (eq. A27). Each time an offspring is produced, the alleles it carries undergo mutation with probability $\mu$, in which case the new value encoded by the mutated allele is sampled in a Gaussian distribution centred on the parental value with standard deviation $\sigma$, truncated such that allelic values are kept within bounds (i.e., between zero and one for sex allocation). We let simulations run for $t_{\mathrm{max}}$ generations. Every $t_{\mathrm{mes}}$ generations, we record the sex allocation $x$ and trait $z$ values of $n_{\mathrm{mes}}$ randomly sampled individuals in the population.

Figure 1 was generated using this program.

### Appendix B

### Evolution of phenotypic associations via the evolution of genetic architecture

#### B.1 Evolution of recombination suppression

##### B.1.1 The model

We consider the same life cycle as in the baseline model, which is detailed in Section 2.1 of the main text, except that upon diploid zygote formation, we assume that the two haplotypes carried by a zygote undergo recombination between the loci encoding sex allocation $x$ and trait $z$ with probability $r$, in which case they exchange alleles, as illustrated in Figure S1.

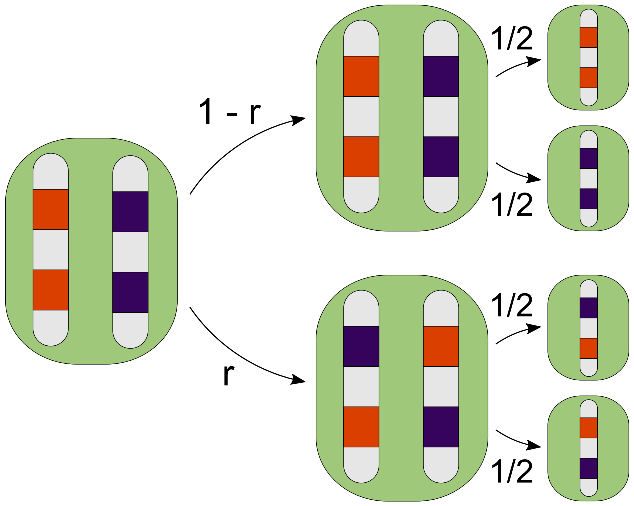

Figure S1: Zygotic recombination. The maternally and paternally inherited chromosomes carry orange and purple alleles at each locus, respectively. With probability $1-r$, no recombination event takes place between the two loci, and the haplotypes transmitted to haploid seeds are identical to the parental haplotypes. Conversely with probability $r$, recombination occurs and the two haplotypes exchange genetic material, so that the transmitted haplotypes are mixtures of the parental ones.

We assume that a zygote's recombination rate is determined by its diploid genotype at an unlinked recombination rate modifier (say, a modifier found on a different chromosome). Alleles at the modifier are assumed to have additive effects, such that the recombination rate $r_i$ of individual $i$, carrying alleles $r_{i,1}$ and $r_{i,2}$ at the modifier, is given by

$$r_i = \frac{r_{i,1} + r_{i,2}}{2}.$$

We assume that the modifier also follows a 'continuum-of-alleles' model, such that alleles can encode any recombination rate between zero and a half (i.e., $r_{i,j} \in [0, 1/2]$ for all $i \in 1, ..., N$ and $j \in \{1, 2\}$). Alleles at the modifier undergo recurrent mutations of small-effect, so that the recombination rate $r$ evolves gradually, and jointly with sex allocation $x$ and trait $z$.

##### B.1.2 Evolutionary analysis

To better understand how selection operates at the recombination modifier, we begin by studying a version of the model in which only recombination evolves. We assume the sex allocation and trait loci to each be polymorphic for two alleles, denoted $x_{♀}$ and $x_{♂}$, and $z_{♀}$ and $z_{♂}$, which encode fixed female- and male-biased sex allocation and trait values, respectively. In this setup, we consider the invasion of a mutant allele at the modifier encoding recombination rate $r_{\mathrm{m}}$ in a resident population fixed for rate $r$.

###### B.1.2.1 Invasion matrix

From the point-of-view of the modifier locus, the population is structured into four classes, each corresponding to one of the four possible haplotypes found in adults. We number these classes (haplotypes) from one to four as follows

$$\begin{aligned} \mathbf{y}_1 &= (x_{♀}, z_{♀}), \\ \mathbf{y}_2 &= (x_{♂}, z_{♂}), \\ \mathbf{y}_3 &= (x_{♀}, z_{♂}), \\ \mathbf{y}_4 &= (x_{♂}, z_{♀}). \end{aligned} \tag{B1}$$

Haplotypes 1 and 2 carry matching alleles at the two loci, as haplotype 1 codes for a female-biased sex allocation in association with a female-biased trait value, and haplotype 2 codes for male-biased sex allocation and a male-biased trait value. Haplotypes 3 and 4, meanwhile, carry mismatched alleles and code for female- or male-biased sex allocation associated with male- and female-biased trait values, respectively. We refer to haplotypes 1 and 2 as 'adapted' haplotypes and haplotypes 3 and 4 as 'maladapted' haplotypes hereafter, for brevity.

The invasion dynamics of a mutant at the modifier can be described by the following recursion,

$$\mathbf{n}_{t+1} = \mathbf{W}(r_{\mathrm{m}}, r) \cdot \mathbf{n}_t, \tag{B2}$$

where $\mathbf{n}_t = \big(n_{1,t},\ n_{2,t},\ n_{3,t},\ n_{4,t}\big)$ is a vector whose $i^{\text{th}}$ entry $n_{i,t}$ gives the number of mutant individuals of class $i$ present in the population at time $t$, and $\mathbf{W}(r_{\mathrm{m}}, r)$ is a $4 \times 4$ invasion matrix whose $(i, j)$-entry $w_{ij}(r_{\mathrm{m}}, r)$ denotes the number of successful mutant offspring of class $i$ (i.e., carrying haplotype $\mathbf{y}_i$) produced by a focal mutant of class $j$ (i.e., carrying haplotype $\mathbf{y}_j$). We refrain from giving the full $\mathbf{W}(r_{\mathrm{m}}, r)$ matrix here, as it is large, but it is available in the supplementary *Mathematica* notebook "AppendixB1-Recombination_Evolution.nb".

###### B.1.2.2 Selection gradient

From standard theory on selection in class-structured population (eq. A2 in Taylor and Frank, 1996; see also Caswell, 2001; Taylor, 1990; Avila and Mullon, 2023), the selection gradient on recombination can be computed as

$$s(r) = \mathbf{v}^{\circ}(r) \cdot \mathbf{D}(r) \cdot \mathbf{q}^{\circ}(r), \tag{B3}$$

where

$$\mathbf{q}^{\circ}(r) = \Big(q_1^{\circ}(r),\ q_2^{\circ}(r),\ q_3^{\circ}(r),\ q_4^{\circ}(r)\Big) \tag{B4}$$

is the vector of asymptotic class-frequencies under neutrality (i.e., when $r_{\mathrm{m}} = r$), which corresponds to the right eigenvector of $\mathbf{W}^{\circ}(r) = \mathbf{W}(r, r)$, the invasion matrix under neutrality, normalised such that class-frequencies sum to one, i.e.,

$$\sum_{i=1}^{4} q_i^{\circ}(r) = 1, \tag{B5}$$

matrix $\mathbf{D}(r)$ is the matrix of first derivatives of $\mathbf{W}(r_{\mathrm{m}}, r)$, whose $(i, j)$-entry $d_{ij}(r)$ is given by

$$d_{ij}(r) = \left.\frac{\partial w_{ij}(r_{\mathrm{m}}, r)}{\partial r_{\mathrm{m}}}\right|_{r_{\mathrm{m}}=r}, \tag{B6}$$

and

$$\mathbf{v}^{\circ}(r) = \Big(v_1^{\circ}(r),\ v_2^{\circ}(r),\ v_3^{\circ}(r),\ v_4^{\circ}(r)\Big) \tag{B7}$$

is the vector of class-specific reproductive values under neutrality (when $r_{\mathrm{m}} = r$), which is given by the left eigenvector of $\mathbf{W}^{\circ}(r)$ normalised such that the mean reproductive value in the population is one, i.e.,

$$\mathbf{v}^{\circ}(r) \cdot \mathbf{q}^{\circ}(r) = 1, \tag{B8}$$

and measures the asymptotic demographic contribution of individuals carrying a given haplotype to the future population. Applying eq. (B3) to $\mathbf{W}(r_{\mathrm{m}}, r)$, we obtain after some straightforward rearrangements that the selection gradient on recombination is given by

$$s(r) = -\frac{1}{2}\Big(v^{\circ}_{\mathrm{A}}(r) - v^{\circ}_{\mathrm{M}}(r)\Big)\Big(Q^{\circ}_{\mathrm{A}}(r) - Q^{\circ}_{\mathrm{M}}(r)\Big), \tag{B9a}$$

where,

$$v^{\circ}_{\mathrm{A}}(r) = \frac{v^{\circ}_{1}(r) + v^{\circ}_{2}(r)}{2} \quad \text{and} \quad v^{\circ}_{\mathrm{M}}(r) = \frac{v^{\circ}_{3}(r) + v^{\circ}_{4}(r)}{2} \tag{B9b}$$

are the reproductive values of adapted haplotypes $\mathbf{y}_1$ and $\mathbf{y}_2$, and maladapted haplotypes $\mathbf{y}_3$ and $\mathbf{y}_4$, respectively; and $Q^{\circ}_{\mathrm{A}}(r)$ and $Q^{\circ}_{\mathrm{M}}(r)$ are the frequencies of double-heterozygous zygotes in the population, which are given by

$$Q^{\circ}_{\mathrm{A}}(r) = q^{\circ}_{1}(r)q^{\circ}_{2}(r)\left(\frac{F(\mathbf{y}_1)}{\overline{F}(\mathbf{Y})}\frac{M(\mathbf{y}_2)}{\overline{M}(\mathbf{Y})} + \frac{F(\mathbf{y}_2)}{\overline{F}(\mathbf{Y})}\frac{M(\mathbf{y}_1)}{\overline{M}(\mathbf{Y})}\right) \tag{B9c}$$

and

$$Q^{\circ}_{\mathrm{M}}(r) = q^{\circ}_{3}(r)q^{\circ}_{4}(r)\left(\frac{F(\mathbf{y}_3)}{\overline{F}(\mathbf{Y})}\frac{M(\mathbf{y}_4)}{\overline{M}(\mathbf{Y})} + \frac{F(\mathbf{y}_4)}{\overline{F}(\mathbf{Y})}\frac{M(\mathbf{y}_3)}{\overline{M}(\mathbf{Y})}\right), \tag{B9d}$$

where $\mathbf{Y} = (\mathbf{y}_1, \mathbf{y}_2, \mathbf{y}_3, \mathbf{y}_4)$, and

$$\overline{F}(\mathbf{Y}) = \sum_{i=1}^{4} q^{\circ}_{i}(r)F(\mathbf{y}_i) \quad \text{and} \quad \overline{M}(\mathbf{Y}) = \sum_{i=1}^{4} q^{\circ}_{i}(r)M(\mathbf{y}_i) \tag{B9e}$$

are the mean female and male fecundities in the population. Eq. (B9a) correspond to eq. (6) in the main text, see Section 3.1 for interpretation.

##### B.1.3 Individual-based simulations

To confirm the intuition gained from the mathematical analyses described above, we ran individual-based simulations of the full model, where allelic values at both loci encoding $x$ and $z$ evolve jointly with alleles at the recombination modifier.

The program is coded in C++. It simulates a population of haploid individuals with a constant size $N$, in which the $i^{\text{th}}$ individual is characterised by its genotype at the sex allocation and trait locus $x_i$ and $z_i$, and at the recombination modifier $r_i$. The population is initially fixed for arbitrary sex allocation and trait values $x_0 \in (0, 1)$ and $z_0 \in \mathbb{R}$, with free recombination between the two loci ($r_0 = 1/2$). Each generation proceeds as follows. We first determine the female and male fecundities of individuals. For individual $i$, these are given by

$$F(\mathbf{y}_i) = x_i V_{♀}(z_i) \quad \text{and} \quad M(\mathbf{y}_i) = (1 - x_i)V_{♂}(z_i)\,, \tag{B10}$$

respectively, where $\mathbf{y}_i = (x_i, x_i)$, and we use Gaussian functions for the effect of $z$ (eq. 2). To create the next generation, we then generate $N$ haploid offspring. For each offspring, we sample with replacement a maternal parent and a paternal parent with probabilities weighted by individual's fecundity in each sex (eq. B10). The resulting diploid zygote then undergoes recombination between the sex allocation and trait $z$ locus (Fig. S1), with a probability given by its diploid genotype at the recombination rate modifier at which alleles are assumed to be co-dominant, i.e., the recombination rate of the $j^{\text{th}}$ zygote produced is given by

$$r_j = \frac{r_{j,1} + r_{j,2}}{2}, \tag{B11}$$

with $r_{j,1}$ and $r_{j,2}$ the recombination rates encoded by their paternally and maternally inherited alleles at the modifier, respectively. Following recombination, one of the two resulting haplotypes is selected to be transmitted to the haploid offspring, and the other is discarded. Alleles carried by the haploid offspring at the two loci and the modifier then each undergo mutation with probability $\mu$ for the $x$ and $z$ loci, and $\mu_n mr$ for the modifier, in which case the new value encoded by the mutated allele is sampled in a Gaussian distribution centred on the parental value with standard deviation $\sigma$ for $x$ and $z$, and $\sigma_{\text{r}}$ for $r$, truncated such that allelic values are kept within bounds (i.e., between zero and one for sex allocation, and between zero and a half for the recombination rate). We let simulations run for $t_{\text{max}}$ generations. Every $t_{\text{mes}}$ generations, we record the sex allocation $x$ and trait $z$ values of $n_{\text{mes}}$ randomly sampled individuals in the population, and the proportion of recombinant zygotes produced in the population at this generation as a proxy of mean recombination rate.

Figures 2 and 3 in the main text were generated using this program.

#### B.2 Evolution of conditional expression

##### B.2.1 Invasion analysis

We study the model described in Section 3.2 assuming the population is initially monomorphic. Because the loci encoding $x$, $a$ and $b$ are unlinked, the evolutionary dynamics of each of these traits can be considered independently of the others, by studying the fate of a rare mutant at each locus when the others are fixed for a resident strategy, as individuals that are mutant at more than one locus at a time will (almost surely) never be formed when mutations are rare. We denote as $\mathbf{y}_\tau$ the phenotype of an individual that carries a mutant allele for trait $\tau \in \{x,\ a,\ b\}$, so that we have

$$\mathbf{y}_x = (x_\mathrm{m},\ a,\ b), \quad \mathbf{y}_a = (x,\ a_\mathrm{m},\ b), \quad \text{and} \quad \mathbf{y}_b = (x,\ a,\ b_\mathrm{m}), \tag{B12}$$

where the "m" subscript denotes a mutant allele. Using this notation, the invasion fitness of a mutant for trait $\tau$ is given by

$$W(\mathbf{y}_\tau, \mathbf{y}) = \frac{1}{2}\left(\frac{F(\mathbf{y}_\tau)}{F(\mathbf{y})} + \frac{M(\mathbf{y}_\tau)}{M(\mathbf{y})}\right). \tag{B13}$$

**Directional selection.** Using eq. (B13), the selection gradient $s_\tau(\mathbf{y})$ on trait $\tau$ can be computed as

$$s_\tau(\mathbf{y}) = \left.\frac{\partial W(\mathbf{y}_\tau, \mathbf{y})}{\partial \tau_\mathrm{m}}\right|_{\mathbf{y}_\tau = \mathbf{y}}. \tag{B14}$$

Applying eq. (B14) to $x$, $a$ and $b$, we find that the selection gradient on each trait is given by

$$s_x(\mathbf{y}) = \frac{1}{2}\left[\frac{1}{x} - \frac{1}{1-x} + a\left(\frac{V'_{♀}(ax+b)}{V_{♀}(ax+b)} + \frac{V'_{♂}(ax+b)}{V_{♂}(ax+b)}\right)\right], \tag{B15a}$$

$$s_a(\mathbf{y}) = \frac{x}{2}\left(\frac{V'_{♀}(ax+b)}{V_{♀}(ax+b)} + \frac{V'_{♂}(ax+b)}{V_{♂}(ax+b)}\right) - 2ac, \tag{B15b}$$

and

$$s_b(\mathbf{y}) = \frac{1}{2}\left(\frac{V'_{♀}(ax+b)}{V_{♀}(ax+b)} + \frac{V'_{♂}(ax+b)}{V_{♂}(ax+b)}\right). \tag{B15c}$$

Solving for the equilibrium $\mathbf{y}^* = (x^*,\ a^*,\ b^*)$ defined as

$$\mathbf{y}^* = (x^*,\ a^*,\ b^*), \quad \text{such that} \quad s_\tau(\mathbf{y}^*) = 0 \quad \text{for all} \quad \tau \in \{x,\ a,\ b\}, \tag{B16}$$

we obtain

$$x^* = \frac{1}{2},\ a^* = 0, \text{ and } b^* \text{ such that } \frac{V'_{♀}(b^*)}{V_{♀}(b^*)} = -\frac{V'_{♂}(b^*)}{V_{♂}(b^*)}, \tag{B17}$$

which corresponds to eq. (9) in the main text. The equilibrium $\mathbf{y}^*$ can be shown to be convergence stable following the same line of argument as in our baseline model (Appendix A.1.1). Thus, directional selection leads individuals to allocate equally to their male and female functions ($x^* = 1/2$, as before), and to express a trait value $z^* = b^*$ that is independent of sex allocation ($a^* = 0$) and corresponds to the equilibrium trait $z^*$ obtained in our baseline model (eq. A6).

**Disruptive selection.** Once the population expresses $\mathbf{y}^*$, each trait may experience stabilising selection or negative frequency-dependent disruptive selection. The nature of selection at $\mathbf{y}^*$ on each trait independently is determined by the sign of the disruptive selection coefficient on trait $\tau$, $h_{\tau\tau}(\mathbf{y}^*)$, which is given by

$$h_{\tau\tau}(\mathbf{y}^*) = \left.\frac{\partial^2 W(\mathbf{y}_\tau, \mathbf{y})}{\partial {\tau_{\mathrm{m}}}^2}\right|_{\mathbf{y}_\tau = \mathbf{y} = \mathbf{y}^*}. \tag{B18}$$

When $h_{\tau\tau}(\mathbf{y}^*) \leqslant 0$, $\tau$ is under stabilising selection. Conversely, when $h_{\tau\tau}(\mathbf{y}^*) > 0$, $\tau$ is under disruptive selection. Applying eq. (B18) to sex allocation $x$, we obtain

$$h_{xx}(\mathbf{y}^*) = 0, \tag{B19}$$

indicating that the singular sex allocation $x^* = 1/2$ is under stabilising selection. As for the sex allocation-dependent and -independent components of trait $z$, $a$ and $b$, we have

$$h_{aa}(\mathbf{y}^*) = \frac{1}{8}\left(\frac{V''_{♀}(b^*)}{V_{♀}(b^*)} + \frac{V''_{♂}(b^*)}{V_{♂}(b^*)}\right) - 2c \quad \text{and} \quad h_{bb}(\mathbf{y}^*) = \frac{1}{2}\left(\frac{V''_{♀}(b^*)}{V_{♀}(b^*)} + \frac{V''_{♂}(b^*)}{V_{♂}(b^*)}\right). \tag{B20}$$

Eq. (B20) reveals that $a$ and $b$ may be under disruptive selection and undergo the gradual emergence of polymorphism if $V_{♀}(z)$ and $V_{♂}(z)$ are accelerating at $z = b^*$. This case will not be given further consideration analytically, as it is more complicated, but we investigate it using simulations later.

##### B.2.2 Amplification of existing polymorphism in sex allocation

Assuming that trait $z$ is under stabilising selection, the above analysis shows that an initially monomorphic population converges to an equilibrium $\mathbf{y}^*$ and should remain monomorphic in the weak mutation limit, such that there can be no standing variation for either sex allocation or trait $z$. To relax this assumption, we consider a population in which two alleles encoding sex allocations $x_{♀}$ and $x_{♂}$ segregate at the sex allocation locus (e.g., owing to mutation-selection balance). We study the evolution of $a$ and $b$ in the presence of such sex allocation polymorphism, and then consider the consequences of this evolution for selective pressures acting on allele $x_{♀}$ and $x_{♂}$. Throughout, sex allocations $x_{♀}$ and $x_{♂}$ are assumed to deviate from the equilibrium sex allocation $x^*$ according to

$$x_{♀} = x^* + \Delta x \quad \text{and} \quad x_{♂} = x^* - \Delta x, \tag{B21}$$

where $\Delta x$ is a small deviation.

**Coexistence of $x_{♀}$ and $x_{♂}$.** We first characterise the coexistence of the two alleles at the sex allocation locus, and determine the frequencies $\hat{p}$ and $1-\hat{p}$ at which allele $x_{♀}$ and $x_{♂}$ are maintained at equilibrium, respectively. Denoting as $\mathbf{y}_{♀} = (x_{♀},\ a,\ b)$ and $\mathbf{y}_{♂} = (x_{♂},\ a,\ b)$ the phenotype of individuals carrying allele $x_{♀}$ and $x_{♂}$, respectively, we obtain that the change in frequency of allele $x_{♀}$ is

$$\Delta p = p\frac{F(\mathbf{y}_{♀})}{\overline{F}(\mathbf{y}_{♀},\mathbf{y}_{♂})}\left(p\frac{M(\mathbf{y}_{♀})}{\overline{M}(\mathbf{y}_{♀},\mathbf{y}_{♂})} + \frac{1-p}{2}\frac{M(\mathbf{y}_{♂})}{\overline{M}(\mathbf{y}_{♀},\mathbf{y}_{♂})}\right) + (1-p)\frac{F(\mathbf{y}_{♂})}{\overline{F}(\mathbf{y}_{♀},\mathbf{y}_{♂})}\frac{p}{2}\frac{M(\mathbf{y}_{♀})}{\overline{M}(\mathbf{y}_{♀},\mathbf{y}_{♂})} - p, \tag{B22}$$

where

$$\overline{F}(\mathbf{y}_{♀},\mathbf{y}_{♂}) = pF(\mathbf{y}_{♀}) + (1-p)F(\mathbf{y}_{♂}) \quad \text{and} \quad \overline{M}(\mathbf{y}_{♀},\mathbf{y}_{♂}) = pM(\mathbf{y}_{♀}) + (1-p)M(\mathbf{y}_{♂}) \tag{B23}$$

are the mean female and male fecundities in the population, respectively. Eq. (B22) can be understood as follows. Individuals carrying $x_{♀}$ are present in frequency $p$ in the population, and each produces $F(\mathbf{y}_{♀})$ ovules, which will compete for recruitment with the seeds produced by the whole population ($\overline{F}(\mathbf{y}_{♀},\mathbf{y}_{♂})$). An ovule carrying $x_{♀}$ is fertilised by pollen carrying $x_{♀}$ with probability $pM(\mathbf{y}_{♀})/\overline{M}(\mathbf{y}_{♀},\mathbf{y}_{♂})$, which leads to an $x_{♀}/x_{♀}$ diploid zygote and thus only to seeds carrying allele $x_{♀}$; and by pollen carrying $x_{♂}$ with probability $(1-p)M(\mathbf{y}_{♂})/\overline{M}(\mathbf{y}_{♀}, \mathbf{y}_{♂})$, which leads to a $x_{♀}/x_{♂}$ zygote that produces a proportion $1/2$ of seeds carrying $x_{♀}$. Individuals carrying $x_{♂}$ are present in frequency $1-p$ and each produce $F(\mathbf{y}_{♂})$ ovules. These ovules give rise to a proportion of $1/2$ of $x_{♀}$ seeds when they are fertilised by pollen carrying $x_{♀}$, which occur with probability $pM(\mathbf{y}_{♀})/\overline{M}(\mathbf{y}_{♀}, \mathbf{y}_{♂})$.

The equilibrium frequency of allele $x_{♀}$ is defined as

$$\hat{p}, \quad \text{such that} \quad \Delta p = 0. \tag{B24}$$

Solving for $\hat{p}$ using eq. (B22), we find

$$\hat{p} = \frac{2F(\mathbf{y}_{♂})M(\mathbf{y}_{♂}) - F(\mathbf{y}_{♂})M(\mathbf{y}_{♀}) - F(\mathbf{y}_{♀})M(\mathbf{y}_{♂})}{2\big[F(\mathbf{y}_{♀}) - F(\mathbf{y}_{♂})\big]\big[M(\mathbf{y}_{♀}) - M(\mathbf{y}_{♂})\big]}. \tag{B25}$$

**Invasion matrix.** We consider the spread of a mutant expressing sex allocation-dependent and independent trait components $a_{\text{m}}$ and $b_{\text{m}}$ in a resident population fixed for $a$ and $b$ at an ecological equilibrium (i.e., where allele $x_{♀}$ is present at frequency $\hat{p}$). Denoting as $\mathbf{y}_{♀}^{\text{m}} = (x_{♀},\ a_{\text{m}},\ b_{\text{m}})$ and $\mathbf{y}_{♂}^{\text{m}} = (x_{♂},\ a_{\text{m}},\ b_{\text{m}})$ the phenotypes expressed by a mutant carrying allele $x_{♀}$ and $x_{♂}$ at the sex allocation locus, respectively, the dynamics of the mutant sub-population can be described by the recursion

$$\mathbf{n}_{t+1} = \mathbf{W}(\mathbf{y}_{♀}^{\text{m}}, \mathbf{y}_{♂}^{\text{m}} | \mathbf{y}_{♀}, \mathbf{y}_{♂}) \cdot \mathbf{n}_t, \tag{B26}$$

where $\mathbf{n}_t = (n_{1,t}, n_{2,t})$ gives the number of mutants carrying sex allocation alleles $x_{♀}$ and $x_{♂}$ (which form two classes of mutants), and $\mathbf{W}(\mathbf{y}_{♀}^{\text{m}}, \mathbf{y}_{♂}^{\text{m}} | \mathbf{y}_{♀}, \mathbf{y}_{♂})$ is a $2 \times 2$ invasion matrix, whose $(i,j)$-entry $w_{ij}(\mathbf{y}_{♀}^{\text{m}}, \mathbf{y}_{♂}^{\text{m}} | \mathbf{y}_{♀}, \mathbf{y}_{♂})$ gives the number of successful mutant offspring of class $i$ produced by a mutant carrying of class $j$. The elements of the invasion matrix are given by

$$w_{11}(\mathbf{y}^{\mathrm{m}}_{♀},\mathbf{y}^{\mathrm{m}}_{♂}|\mathbf{y}_{♀},\mathbf{y}_{♂}) = \frac{1}{2}\Bigg[\frac{F(\mathbf{y}^{\mathrm{m}}_{♀})}{\overline{F}(\mathbf{y}_{♀},\mathbf{y}_{♂})}\left(\hat{p}\frac{M(\mathbf{y}_{♀})}{\overline{M}(\mathbf{y}_{♀},\mathbf{y}_{♂})}+\frac{1-\hat{p}}{2}\frac{M(\mathbf{y}_{♂})}{\overline{M}(\mathbf{y}_{♀},\mathbf{y}_{♂})}\right) + \frac{M(\mathbf{y}^{\mathrm{m}}_{♀})}{\overline{M}(\mathbf{y}_{♀},\mathbf{y}_{♂})}\left(\hat{p}\frac{F(\mathbf{y}_{♀})}{\overline{F}(\mathbf{y}_{♀},\mathbf{y}_{♂})}+\frac{1-\hat{p}}{2}\frac{F(\mathbf{y}_{♂})}{\overline{F}(\mathbf{y}_{♀},\mathbf{y}_{♂})}\right)\Bigg], \tag{B27a}$$

$$w_{12}(\mathbf{y}^{\mathrm{m}}_{♀},\mathbf{y}^{\mathrm{m}}_{♂}|\mathbf{y}_{♀},\mathbf{y}_{♂}) = \frac{\hat{p}}{4}\left(\frac{F(\mathbf{y}^{\mathrm{m}}_{♂})}{\overline{F}(\mathbf{y}_{♀},\mathbf{y}_{♂})}\frac{M(\mathbf{y}_{♀})}{\overline{M}(\mathbf{y}_{♀},\mathbf{y}_{♂})}+\frac{M(\mathbf{y}^{\mathrm{m}}_{♂})}{\overline{M}(\mathbf{y}_{♀},\mathbf{y}_{♂})}\frac{M(\mathbf{y}_{♀})}{\overline{M}(\mathbf{y}_{♀},\mathbf{y}_{♂})}\right) \tag{B27b}$$

$$w_{21}(\mathbf{y}^{\mathrm{m}}_{♀},\mathbf{y}^{\mathrm{m}}_{♂}|\mathbf{y}_{♀},\mathbf{y}_{♂}) = \frac{1-\hat{p}}{4}\left(\frac{F(\mathbf{y}^{\mathrm{m}}_{♀})}{\overline{F}(\mathbf{y}_{♀},\mathbf{y}_{♂})}\frac{M(\mathbf{y}_{♂})}{\overline{M}(\mathbf{y}_{♀},\mathbf{y}_{♂})}+\frac{M(\mathbf{y}^{\mathrm{m}}_{♀})}{\overline{M}(\mathbf{y}_{♀},\mathbf{y}_{♂})}\frac{M(\mathbf{y}_{♂})}{\overline{M}(\mathbf{y}_{♀},\mathbf{y}_{♂})}\right) \tag{B27c}$$

$$w_{22}(\mathbf{y}^{\mathrm{m}}_{♀},\mathbf{y}^{\mathrm{m}}_{♂}|\mathbf{y}_{♀},\mathbf{y}_{♂}) = \frac{1}{2}\Bigg[\frac{F(\mathbf{y}^{\mathrm{m}}_{♂})}{\overline{F}(\mathbf{y}_{♀},\mathbf{y}_{♂})}\left(\frac{\hat{p}}{2}\frac{M(\mathbf{y}_{♀})}{\overline{M}(\mathbf{y}_{♀},\mathbf{y}_{♂})}+(1-\hat{p})\frac{M(\mathbf{y}_{♂})}{\overline{M}(\mathbf{y}_{♀},\mathbf{y}_{♂})}\right) + \frac{M(\mathbf{y}^{\mathrm{m}}_{♂})}{\overline{M}(\mathbf{y}_{♀},\mathbf{y}_{♂})}\left(\frac{\hat{p}}{2}\frac{F(\mathbf{y}_{♀})}{\overline{F}(\mathbf{y}_{♀},\mathbf{y}_{♂})}+(1-\hat{p})\frac{F(\mathbf{y}_{♂})}{\overline{F}(\mathbf{y}_{♀},\mathbf{y}_{♂})}\right)\Bigg]. \tag{B27d}$$

These were constructed following the same logic used to compute allelic frequency change $\Delta p$ above (eq. B22).

**Selection gradient and singular values.** From this matrix, the selection gradient on trait component $\tau \in \{a, b\}$, is given by

$$s_{\tau}(\mathbf{y}_{♀},\mathbf{y}_{♂}) = \mathbf{v}^{\circ}(\mathbf{y}_{♀},\mathbf{y}_{♂})\cdot\mathbf{D}^{\tau}(\mathbf{y}_{♀},\mathbf{y}_{♂})\cdot\mathbf{q}^{\circ}(\mathbf{y}_{♀},\mathbf{y}_{♂}), \tag{B28}$$

where vector $\mathbf{q}^{\circ}(\mathbf{y}_{♀},\mathbf{y}_{♂}) = \left(q^{\circ}_{1}(\mathbf{y}_{♀},\mathbf{y}_{♂}),\ q^{\circ}_{2}(\mathbf{y}_{♀},\mathbf{y}_{♂})\right)$ is the vector of asymptotic class-frequencies (i.e., of frequencies of individuals carrying allele $x_{♀}$ and $x_{♂}$), and is given by the right eigenvector of $\mathbf{W}^{\circ}(\mathbf{y}_{♀},\mathbf{y}_{♂}) = \mathbf{W}(\mathbf{y}_{♀},\mathbf{y}_{♂}|\mathbf{y}_{♀},\mathbf{y}_{♂})$, the invasion matrix under neutrality, normalised such that

$$q^{\circ}_{1}(\mathbf{y}_{♀},\mathbf{y}_{♂}) + q^{\circ}_{2}(\mathbf{y}_{♀},\mathbf{y}_{♂}) = 1; \tag{B29}$$

matrix $\mathbf{D}^{\tau}(\mathbf{y}_{♀}, \mathbf{y}_{♂})$ is a $2 \times 2$ matrix whose $(i,j)$-entry $d_{ij,\tau}(\mathbf{y}_{♀}, \mathbf{y}_{♂})$ is given by

$$d_{ij}^{\tau}(\mathbf{y}_{♀}, \mathbf{y}_{♂}) = \left. \frac{\partial w_{ij}(\mathbf{y}_{♀}^{\mathrm{m}}, \mathbf{y}_{♂}^{\mathrm{m}} | \mathbf{y}_{♀}, \mathbf{y}_{♂})}{\partial \tau_{\mathrm{m}}} \right|_{\substack{\mathbf{y}_{♀}^{\mathrm{m}}=\mathbf{y}_{♀} \\ \mathbf{y}_{♂}^{\mathrm{m}}=\mathbf{y}_{♂}}} \tag{B30}$$

for $\tau_{\mathrm{m}} \in \{a_{\mathrm{m}}, b_{\mathrm{m}}\}$ and measures the effect of a change in trait $\tau$ on the number of successful mutant descendants carrying sex allocation allele $x_i$ produced by a mutant carrying $x_j$; and $\mathbf{v}^{\circ}(\mathbf{y}_{♀}, \mathbf{y}_{♂}) = \big(v_1^{\circ}(\mathbf{y}_{♀}, \mathbf{y}_{♂}),\ v_2^{\circ}(\mathbf{y}_{♀}, \mathbf{y}_{♂})\big)$ is the vector of class reproductive values under neutrality, and is given by the left eigenvector of $\mathbf{W}^{\circ}(\mathbf{y}_{♀}, \mathbf{y}_{♂})$ normalised such that

$$\mathbf{v}^{\circ}(\mathbf{y}_{♀}, \mathbf{y}_{♂}) \cdot \mathbf{q}^{\circ}(\mathbf{y}_{♀}, \mathbf{y}_{♂}) = 1. \tag{B31}$$

Our aim is to compute

$$a^* \quad \text{and} \quad b^* \quad \text{such that} \quad s_a(\mathbf{y}_{♀}^*, \mathbf{y}_{♂}^*) = s_b(\mathbf{y}_{♀}^*, \mathbf{y}_{♂}^*) = 0, \tag{B32}$$

where $\mathbf{y}_i^* = \big(x_i,\ a^*,\ b^*\big)$, $i \in \{1,2\}$, i.e., to find those trait values of $a$ and $b$ for which directional selection vanishes when there is polymorphism for sex allocation. These equilibria are difficult to compute in general, but they are functions of $\Delta x$ and can thus be approximated by a Taylor expansion respect to $\Delta x$ as

$$a^*(\Delta x) = a_{(0)}^* + \Delta x\, a_{(1)}^* + \frac{\Delta x^2}{2} a_{(2)}^* + \mathcal{O}(\Delta x^3) \quad \text{and} \quad b^*(\Delta x) = b_{(0)}^* + \Delta x\, b_{(1)}^* + \frac{\Delta x^2}{2} b_{(2)}^* + \mathcal{O}(\Delta x^3), \tag{B33}$$

where terms denoted $a_{(n)}^*$ and $b_{(n)}^*$, $n \in \{0, 1, 2, ...\}$, are $n^{\text{th}}$ order perturbations of $a^*$ and $b^*$, defined as

$$a_{(n)}^* = \left. \frac{\partial^n a^*(\Delta x)}{\partial \Delta x^n} \right|_{\Delta x = 0} \quad \text{and} \quad b_{(n)}^* = \left. \frac{\partial^n b^*(\Delta x)}{\partial \Delta x^n} \right|_{\Delta x = 0}. \tag{B34}$$

To calculate the perturbations of $a^*$ and $b^*$, we note that the selection gradients on each of those traits can too be expanded with respect to $\Delta x$, which yields

$$
\begin{aligned}
s_\tau(\mathbf{y}_♀, \mathbf{y}_♂) = {} & \mathbf{v}^\circ_{(0)} \cdot \mathbf{D}^\tau_{(0)} \cdot \mathbf{q}^\circ_{(0)} \\
& + \Delta x \left[ \mathbf{v}^\circ_{(1)} \cdot \mathbf{D}^\tau_{(0)} \cdot \mathbf{q}^\circ_{(0)} + \mathbf{v}^\circ_{(0)} \cdot \mathbf{D}^\tau_{(1)} \cdot \mathbf{q}^\circ_{(0)} + \mathbf{v}^\circ_{(0)} \cdot \mathbf{D}^\tau_{(0)} \cdot \mathbf{q}^\circ_{(1)} \right] \\
& + \frac{\Delta x^2}{2} \Big[ \mathbf{v}^\circ_{(2)} \cdot \mathbf{D}^\tau_{(0)} \cdot \mathbf{q}^\circ_{(0)} + \mathbf{v}^\circ_{(0)} \cdot \mathbf{D}^\tau_{(2)} \cdot \mathbf{q}^\circ_{(0)} + \mathbf{v}^\circ_{(0)} \cdot \mathbf{D}^\tau_{(0)} \cdot \mathbf{q}^\circ_{(2)} \\
& \qquad + \mathbf{v}^\circ_{(1)} \cdot \mathbf{D}^\tau_{(1)} \cdot \mathbf{q}^\circ_{(0)} + \mathbf{v}^\circ_{(1)} \cdot \mathbf{D}^\tau_{(0)} \cdot \mathbf{q}^\circ_{(1)} + \mathbf{v}^\circ_{(0)} \cdot \mathbf{D}^\tau_{(1)} \cdot \mathbf{q}^\circ_{(1)} \Big] + \mathcal{O}(\Delta x^3),
\end{aligned} \tag{B35}
$$

where we dropped arguments $(\mathbf{y}_♀, \mathbf{y}_♂)$ for brevity, and elements $\mathbf{v}^\circ_{(n)}$, $\mathbf{D}^\tau_{(n)}$ and $\mathbf{q}^\circ_{(n)}$ are $n^{\text{th}}$ perturbations of $\mathbf{v}^\circ$, $\mathbf{D}^\tau$ and $\mathbf{q}^\circ$, respectively, defined as

$$
\mathbf{C}_{(n)} = \left. \frac{\partial^n \mathbf{C}}{\partial \Delta x^n} \right|_{\Delta x = 0}, \quad \text{with} \quad \mathbf{C} \in \{\mathbf{v}^\circ,\ \mathbf{D}^\tau,\ \mathbf{q}^\circ\}. \tag{B36}
$$

**Zero-th order perturbations.** To compute the selection gradients on $a$ and $b$ to zero-th order in $\Delta x$ (i.e., setting $\Delta x = 0$), we must calculate the zero-th order perturbations $\mathbf{v}^\circ_{(0)}$ and $\mathbf{q}^\circ_{(0)}$ of the left and right eigenvector of $\mathbf{W}^\circ$, the invasion matrix under neutrality. The right eigenvector of $\mathbf{W}^\circ$ satisfies

$$
\mathbf{W}^\circ \cdot \mathbf{q}^\circ = \mathbf{q}^\circ, \tag{B37}
$$

since the leading eigenvalue of $\mathbf{W}^\circ$ is $\lambda^\circ = 1$ by definition. Expanding $\mathbf{W}^\circ$ and $\mathbf{q}^\circ$ with respect to $\Delta x$, i.e., writing

$$
\mathbf{W}^\circ = \mathbf{W}^\circ_{(0)} + \Delta x\, \mathbf{W}^\circ_{(1)} + \frac{\Delta x^2}{2}\, \mathbf{W}^\circ_{(2)} + \mathcal{O}(\Delta x^3), \tag{B38}
$$

and

$$
\mathbf{q}^\circ = \mathbf{q}^\circ_{(0)} + \Delta x\, \mathbf{q}^\circ_{(1)} + \frac{\Delta x^2}{2} \mathbf{q}^\circ_{(2)} + \mathcal{O}(\Delta x^3), \tag{B39}
$$

and inserting these into eq. (B37), we find that $\mathbf{q}^\circ_{(0)}$ satisfies

$$
\mathbf{W}^\circ_{(0)} \cdot \mathbf{q}^\circ_{(0)} = \mathbf{q}^\circ_{(0)}. \tag{B40}
$$

Solving eq. (B40) for $\mathbf{q}^\circ_{(0)}$, with the constraint that its elements must sum to one, i.e., that $q^\circ_{(0),2} = 1 - q^\circ_{(0),1}$, we obtain

$$
\mathbf{q}^\circ_{(0)} = \big(\hat{p},\ 1 - \hat{p}\big). \tag{B41}
$$

Proceeding in a similar manner for $\mathbf{v}^{\circ}_{(0)}$, with the added constraint that its elements must satisfy

$$\mathbf{v}^{\circ}_{(0)} \cdot \mathbf{q}^{\circ}_{(0)} = 1 \quad \Leftrightarrow \quad v^{\circ}_{(0),2} = \frac{1 - \hat{p}\, v^{\circ}_{(0),1}}{1 - \hat{p}}, \tag{B42}$$

we find

$$\mathbf{v}^{\circ}_{(0)} = \big(1,\ 1\big). \tag{B43}$$

Using eqs. (B41) and (B43), the zero-th order selection gradients on $a$ and $b$ are thus given by

$$\begin{aligned} s_a^{(0)} &= \mathbf{v}^{\circ}_{(0)} \cdot \mathbf{D}^{a}_{(0)} \cdot \mathbf{q}^{\circ}_{(0)} \\ &= \frac{\omega_{♀} + \omega_{♂}}{2} \left[ \theta_{♀} \frac{\omega_{♀}}{\omega_{♀} + \omega_{♂}} + \theta_{♂} \frac{\omega_{♂}}{\omega_{♀} + \omega_{♂}} - b_{(0)} - \frac{a_{(0)}}{2} \left( 1 + 8 \frac{c}{\omega_{♀} + \omega_{♂}} \right) \right], \end{aligned} \tag{B44}$$

and

$$s_b^{(0)} = \mathbf{v}^{\circ}_{(0)} \cdot \mathbf{D}^{b}_{(0)} \cdot \mathbf{q}^{\circ}_{(0)} = (\omega_{♀} + \omega_{♂}) \left( \theta_{♀} \frac{\omega_{♀}}{\omega_{♀} + \omega_{♂}} + \theta_{♂} \frac{\omega_{♂}}{\omega_{♀} + \omega_{♂}} - b_{(0)} - \frac{a_{(0)}}{2} \right). \tag{B45}$$

Solving $s_a^{(0)} = s_b^{(0)} = 0$ for $a^*_{(0)}$ and $b^*_{(0)}$, we obtain

$$a^*_{(0)} = 0 \quad \text{and} \quad b^*_{(0)} = \theta_{♀} \frac{\omega_{♀}}{\omega_{♀} + \omega_{♂}} + \theta_{♂} \frac{\omega_{♂}}{\omega_{♀} + \omega_{♂}}. \tag{B46}$$

These are identical to the singular sex allocation-dependent and -independent components of trait $z$, $a^*$ and $b^*$, obtained in the case of monomorphic population (eq. B17), indicating that $a^*$ and $b^*$ in the presence of polymorphism for sex allocation should only deviate weakly from the values favoured in the absence of polymorphism when $\Delta x$ is small.

**First-order perturbation.** Given $a^*_{(0)}$ and $b^*_{(0)}$, we can now compute the selection gradients on $a$ and $b$ to first order in $\Delta x$ in order to compute first-order perturbations $a^*_{(1)}$ and $b^*_{(1)}$. To this end, we first need to compute $\mathbf{q}^{\circ}_{(1)}$ and $\mathbf{v}^{\circ}_{(1)}$, the first-order perturbations of vectors $\mathbf{q}^{\circ}$ and $\mathbf{v}^{\circ}$. Inserting eqs. (B38) and (B39) into eq. (B37) and neglecting terms of order two and higher, we find that $\mathbf{q}^{\circ}_{(1)}$ must satisfy

$$\mathbf{W}^{\circ}_{(0)} \cdot \mathbf{q}^{\circ}_{(1)} + \mathbf{W}^{\circ}_{(1)} \cdot \mathbf{q}^{\circ}_{(0)} = \mathbf{q}^{\circ}_{(1)}, \tag{B47}$$

with the added constraint that its elements sum to zero, i.e., $q^\circ_{(1),2} = -q^\circ_{(1),1}$. Inserting eq. (B46) into eq. (B47) and solving for $\mathbf{q}^\circ_{(1)}$ yields

$$\mathbf{q}^\circ_{(1)} = (0,\ 0). \tag{B48}$$

Similarly for $\mathbf{v}^\circ_{(1)}$, with the constraint that we must have

$$\mathbf{v}^\circ_{(1)} \cdot \mathbf{q}^\circ_{(0)} + \mathbf{v}^\circ_{(0)} \cdot \mathbf{q}^\circ_{(1)} = 0 \quad \Leftrightarrow \quad v^\circ_{(1),2} = \frac{1 - \hat{p}\, v^\circ_{(1),1}}{1 - \hat{p}}, \tag{B49}$$

we obtain

$$\mathbf{v}^\circ_{(1)} = (0,\ 0). \tag{B50}$$

Inserting eq. (B46), (B48) and (B50) into eq. (B35) up to order $\Delta x$ (i.e., ignoring terms of order $\Delta x^2$ and higher), we find that the selection gradients on $a$ and $b$ to first-order in $\Delta x$ are given by

$$s_a^{(1)} = \Delta x\, \mathbf{v}^\circ_{(0)} \cdot \mathbf{D}^a_{(1)} \cdot \mathbf{q}^\circ_{(0)} = -\Delta x \frac{\omega_{♀} + \omega_{♂}}{4} \left[ 2b_{(1)} + a_{(1)} \left( 1 + 8 \frac{c}{\omega_{♀} + \omega_{♂}} \right) \right] \tag{B51}$$

and

$$s_b^{(1)} = \Delta x\, \mathbf{v}^\circ_{(0)} \cdot \mathbf{D}^b_{(1)} \cdot \mathbf{q}^\circ_{(0)} = -\Delta x \frac{\omega_{♀} + \omega_{♂}}{2} \big( a_{(1)} + 2b_{(1)} \big). \tag{B52}$$

Solving $s_a^{(1)} = s_b^{(1)} = 0$ for $a^*_{(1)}$ and $b^*_{(1)}$ then yields

$$a^*_{(1)} = b^*_{(1)} = 0. \tag{B53}$$

**Second-order perturbation.** We now compute the second-order perturbations $a^*_{(2)}$ and $b^*_{(2)}$. This would normally require computing second-order perturbations of $\mathbf{q}^\circ$ and $\mathbf{v}^\circ$, $\mathbf{q}^\circ_{(2)}$ and $\mathbf{v}^\circ_{(2)}$ (eq. (B35)). However, since

$$\mathbf{v}^\circ_{(0)} \cdot \mathbf{D}^\tau_{(0)} = \mathbf{D}^\tau_{(0)} \cdot \mathbf{q}^\circ_{(0)} = (0,\ 0), \tag{B54}$$

when evaluated at $a_{(0)} = a^*_{(0)}$ and $b_{(0)} = b^*_{(0)}$ (eq. B46), any term involving $\mathbf{q}^\circ_{(2)}$ and $\mathbf{v}^\circ_{(2)}$ will be zero to second order in $\Delta x$ (eq. B35). In fact, because we also have

$$\mathbf{v}^\circ_{(1)} = \mathbf{q}^\circ_{(1)} = (0,\ 0), \tag{B55}$$

the selection gradient on $a$ and $b$ to second order in $\Delta x$ simplifies to

$$\begin{aligned} s_a^{(2)} &= \frac{\Delta x^2}{2}\,\mathbf{v}^{\circ}_{(0)}\cdot\mathbf{D}^{a}_{(2)}\cdot\mathbf{q}^{\circ}_{(0)} \\ &= \Delta x^2\left\{16\omega_{♀}\omega_{♂}\frac{\theta_{♀}-\theta_{♂}}{\omega_{♀}+\omega_{♂}}\hat{p}(1-\hat{p})-\frac{\omega_{♀}+\omega_{♂}}{4}\left[b_{(2)}+\frac{a_{(2)}}{2}\left(1+8\frac{c}{\omega_{♀}+\omega_{♂}}\right)\right]\right\}. \end{aligned} \tag{B56}$$

and

$$s_b^{(2)} = -\Delta x^2\frac{\omega_{♀}+\omega_{♂}}{4}\left(a_{(2)}+2b_{(2)}\right). \tag{B57}$$

Until now, none of our calculations required further consideration of $\hat{p}$, as it was absent from zero-th and first order terms. The equilibrium frequency $\hat{p}$ can also be written as a Taylor expansion around $\Delta x$,

$$\hat{p} = \hat{p}_{(0)} + \Delta x\,\hat{p}_{(1)} + \frac{\Delta x^2}{2}\,\hat{p}_{(2)} + \mathcal{O}(\Delta x^3), \tag{B58}$$

which, using eqs. (B46) and (B53), yields

$$\hat{p} = \frac{1}{2} + \mathcal{O}(\Delta x^3). \tag{B59}$$

Inserting eq. (B59) into eq. (B56), ignoring terms of order $\Delta x^3$ and above, and solving for

$$a^*_{(2)} \quad \text{and} \quad b^*_{(2)} \quad \text{such that} \quad s_a^{(2)} = s_b^{(2)} = 0, \tag{B60}$$

we obtain

$$a^*_{(2)} = 4\frac{\theta_{♀}-\theta_{♂}}{c(\omega_{♀}+\omega_{♂})}\omega_{♀}\omega_{♂} \quad \text{and} \quad b^*_{(2)} = -2\frac{\theta_{♀}-\theta_{♂}}{c(\omega_{♀}+\omega_{♂})}\omega_{♀}\omega_{♂}. \tag{B61}$$

Thus, the singular components of trait $z$ are given by

$$a^* = 4\Delta x^2\frac{\theta_{♀}-\theta_{♂}}{c(\omega_{♀}+\omega_{♂})}\omega_{♀}\omega_{♂} + \mathcal{O}(\Delta x^3), \tag{B62a}$$

and

$$b^* = \theta_{♀}\frac{\omega_{♀}}{\omega_{♀}+\omega_{♂}} + \theta_{♂}\frac{\omega_{♂}}{\omega_{♀}+\omega_{♂}} - 2\Delta x^2\frac{\theta_{♀}-\theta_{♂}}{c(\omega_{♀}+\omega_{♂})}\omega_{♀}\omega_{♂} + \mathcal{O}(\Delta x^3), \tag{B62b}$$

to second order in $\Delta x$. Eq. (B62) corresponds to eq. (11) in the main text; see Section 3.2 for interpretation.

**Consequences of evolved conditional trait expression for selection on sex allocation.** We now consider a population fixed for $a^*$ and $b^*$ (eq. B62) in which sex allocation alleles $x_{♀}$ and $x_{♂}$ segregate ($x_{♂} < x^* < x_{♀}$), and investigate the nature of selection weighing on those alleles given the phenotypic correlation favoured by selection. To this end, we study the fate of a mutant allele $x_{\mathrm{m}}$ at the sex allocation locus. Denoting the phenotype expressed by an individual carrying this allele as $\mathbf{y}_{\mathrm{m}} = (x_{\mathrm{m}},\ a^*,\ b^*)$, the invasion fitness of the mutant is given by

$$W(\mathbf{y}_{\mathrm{m}}, \mathbf{y}_{♀}, \mathbf{y}_{♂}) = \frac{1}{2}\left(\frac{F(\mathbf{y}_{\mathrm{m}})}{\overline{F}(\mathbf{y}_{♀}, \mathbf{y}_{♂})} + \frac{M(\mathbf{y}_{\mathrm{m}})}{\overline{M}(\mathbf{y}_{♀}, \mathbf{y}_{♂})}\right), \tag{B63}$$

where

$$\overline{F}(\mathbf{y}_{♀}, \mathbf{y}_{♂}) = \hat{p}F(\mathbf{y}_{♀}) + (1-\hat{p})F(\mathbf{y}_{♂}) \quad \text{and} \quad \overline{M}(\mathbf{y}_{♀}, \mathbf{y}_{♂}) = \hat{p}M(\mathbf{y}_{♀}) + (1-\hat{p})M(\mathbf{y}_{♂}) \tag{B64}$$

are the average female and male fecundities in the resident population. From eq. (B63), the selection gradient on allele $x_u$, $u \in \{♀, ♂\}$, can be obtained as

$$s_u(\mathbf{y}_{♀}, \mathbf{y}_{♂}) = \left.\frac{\partial W(\mathbf{y}_{\mathrm{m}}, \mathbf{y}_{♀}, \mathbf{y}_{♂})}{\partial \mathbf{y}_{\mathrm{m}}}\right|_{\mathbf{y}_{\mathrm{m}}=\mathbf{y}_u}. \tag{B65}$$

The selection gradients obtained from applying eq. (B65) to eq. (B63) are complicated, and we thus refrain from giving their full expression here. They can however be greatly simplified under the assumption that $\Delta x$ is small. Expressing $a^*$, $b^*$ and $\hat{p}$ to third-order in $\Delta x$, that is writing

$$\hat{p} = \frac{1}{2} + \frac{\Delta x^3}{6}\hat{p}_{(3)} + \mathcal{O}(\Delta x^4), \tag{B66a}$$

$$a^* = 4\Delta x^2 \frac{\theta_{♀} - \theta_{♂}}{c(\omega_{♀} + \omega_{♂})}\omega_{♀}\omega_{♂} + \frac{\Delta x^3}{6}a^*_{(3)} + \mathcal{O}(\Delta x^4), \tag{B66b}$$

and

$$b^* = \theta_{♀}\frac{\omega_{♀}}{\omega_{♀} + \omega_{♂}} + \theta_{♂}\frac{\omega_{♂}}{\omega_{♀} + \omega_{♂}} - 2\Delta x^2 \frac{\theta_{♀} - \theta_{♂}}{c(\omega_{♀} + \omega_{♂})}\omega_{♀}\omega_{♂} + \frac{\Delta x^3}{6}b^*_{(3)} + \mathcal{O}(\Delta x^4), \tag{B66c}$$

where $\hat{p}_{(3)}$, $a^*_{(3)}$ and $b^*_{(3)}$ denote third-order perturbation terms, we insert these into the selection gradients $s_{♀}(\mathbf{y}_{♀}, \mathbf{y}_{♂})$ and $s_{♂}(\mathbf{y}_{♀}, \mathbf{y}_{♂})$, and we find that, to third-order in $\Delta x$,

$$s_{♀}(\mathbf{y}_{♀}, \mathbf{y}_{♂}) = \frac{\Delta x^3}{c}\left(4\omega_{♀}\omega_{♂}\frac{\theta_{♀} - \theta_{♂}}{\omega_{♀} + \omega_{♂}}\right)^2 + \mathcal{O}(\Delta x^4), \tag{B67a}$$

and

$$s_{♂}(\mathbf{y}_{♀}, \mathbf{y}_{♂}) = -\frac{\Delta x^3}{c}\left(4\omega_{♀}\omega_{♂}\frac{\theta_{♀} - \theta_{♂}}{\omega_{♀} + \omega_{♂}}\right)^2 + \mathcal{O}(\Delta x^4), \tag{B67b}$$

where $\hat{p}_{(3)}$, $a^*_{(3)}$ and $b^*_{(3)}$ all vanish. Eq. (B67) corresponds to eq. (12) in Section 3.2 of the main text; see there for interpretation.

##### B.2.3 Individual-based simulations

We ran individual-based simulations coded in C++. The program simulates a population of $N$ haploid individuals characterised by their genotype at the loci encoding $x$, $a$ and $b$, which we denote as $\mathbf{y}_i = (x_i, a_i, b_i)$ for individual $i$. The population is initially fixed for an arbitrary sex allocation $x_0 \in (0, 1)$, and sex allocation-dependent and -independent trait components $a_0 \in \mathbb{R}$ and $b_0 \in \mathbb{R}$. At each time step, we begin by determining the female and male fecundities of individuals, which are given by

$$F(\mathbf{y}_i) = x_i V_{♀}(a_i x_i + b_i) \exp\left(-c{a_i}^2\right) \quad \text{and} \quad M(\mathbf{y}_i) = (1 - x_i) V_{♂}(a_i x_i + b_i) \exp\left(-c{a_i}^2\right), \tag{B68}$$

for individual $i$, where $c > 0$ measures the cost of plasticity. To create the next generation, we generate $N$ haploid offspring. For each offspring, a mother and a father are sampled with replacement from the parental population with a probability proportional to their female and male fecundities, respectively. We assume that the loci encoding the three traits are unlinked, so that the resulting diploid zygote undergoes recombination between each pair of locus with probability $1/2$. Following recombination, one haplotype of the zygote is chosen at random to be inherited by the haploid offspring, and the other one is discarded. Alleles carried by the haploid offspring undergo mutation with probability $\mu$ and the mutated value is sampled in a truncated Gaussian distribution centred on parental value with standard deviation $\sigma$ (as before). We let simulations run for $t_{\text{max}}$ generations. Every $t_{\text{mes}}$ generations, we record the sex allocation $x$ and trait $z$ components $a$ and $b$ of $n_{\text{mes}}$ randomly sampled individuals in the population.

Figure 4 in the main text was generated using this program.

### Appendix C

### The effects of limited dispersal and kin competition

#### C.1 Selection in the island model of dispersal

We first briefly review current theory on how to characterise directional, correlational and quadratic selection in the island model of dispersal.

##### C.1.1 Directional selection

Following established theory (Frank, 1998; Rousset, 2004 for textbooks), the selection gradient on a continuous trait $\tau \in \{x, z\}$ is given by the sum of direct and relatedness-weighted indirect fitness effects of a change in trait $\tau$, i.e., by

$$s_\tau(\mathbf{y}) = \left.\frac{\partial \mathrm{w}(\mathbf{y}_i, \mathbf{y}_{-i}, \mathbf{y})}{\partial \tau_i}\right|_{\mathbf{y}} + R_2^\circ(N-1) \left.\frac{\partial \mathrm{w}(\mathbf{y}_i, \mathbf{y}_{-i}, \mathbf{y})}{\partial \tau_{j\neq i}}\right|_{\mathbf{y}}, \tag{C1}$$

where $\mathrm{w}(\mathbf{y}_i, \mathbf{y}_{-i}, \mathbf{y})$ is the individual fitness, that is the expected number of successful offspring (i.e., that establish as adults), of a focal individual $i$ expressing phenotype $\mathbf{y}_i = (x_i, z_i)$ in a patch where other individuals express $\mathbf{y}_{-i} = (\mathbf{y}_1, \ldots, \mathbf{y}_{i-1}, \mathbf{y}_{i+1}, \ldots, \mathbf{y}_N)$, with $\mathbf{y}_j = (x_j, z_j)$ as the phenotype of individual $j$ in the patch, and the rest of the population expresses $\mathbf{y} = (x, z)$. All derivatives in eq. (C1) are evaluated at $\mathbf{y}_k = \mathbf{y}$ for all $k \in \{1, \ldots, N\}$, which we denote as $\cdot|_{\mathbf{y}}$ for short. $\tau_{j\neq i}$ denotes the value of $\tau$ expressed by a neighbour $j$ of the focal individual $i$. The quantity $R_2^\circ$ is pairwise relatedness under neutrality, that is the probability that two randomly sampled individuals from the same patch are identical-by-descent (IBD) in a monomorphic population. We derive individual fitness and relatedness for our model in Section C.2. The selection gradient eq. (C1) can then be used as explained in Section A.1.1 to determine the convergence stable equilibrium $\mathbf{y}^* = (x^*, z^*)$.

##### C.1.2 Correlational and quadratic selection

Once the population expresses $\mathbf{y}^*$, selection is given by the Hessian matrix $\mathbf{H}(\mathbf{y}^*)$ (eq. A12), whose element $h_{\tau_1\tau_2}(\mathbf{y}^*)$, with $\tau_1 \in \{x, z\}$ and $\tau_2 \in \{x, z\}$, is given by the sum of two terms in the island model,

$$h_{\tau_1\tau_2}(\mathbf{y}^*) = h_{\mathrm{w},\tau_1\tau_2}(\mathbf{y}^*) + h_{\mathrm{r},\tau_1\tau_2}(\mathbf{y}^*) \tag{C2a}$$

(Mullon et al., 2016). The first term is given by

$$\begin{aligned} h_{\mathrm{w},\tau_1\tau_2}(\mathbf{y}^*) = {} & \left.\frac{\partial^2 \mathrm{w}(\mathbf{y}_i, \mathbf{y}_{-i}, \mathbf{y})}{\partial \tau_{1,i} \partial \tau_{2,i}}\right|_{\mathbf{y}^*} + R_2^\circ (N-1) \left( \left.\frac{\partial^2 \mathrm{w}(\mathbf{y}_i, \mathbf{y}_{-i}, \mathbf{y})}{\partial \tau_{1,i} \partial \tau_{2,j\neq i}}\right|_{\mathbf{y}^*} + \left.\frac{\partial^2 \mathrm{w}(\mathbf{y}_i, \mathbf{y}_{-i}, \mathbf{y})}{\partial \tau_{1,j\neq i} \partial \tau_{2,i}}\right|_{\mathbf{y}^*} \right) \\ & + R_3^\circ (N-1)(N-2) \left.\frac{\partial^2 \mathrm{w}(\mathbf{y}_i, \mathbf{y}_{-i}, \mathbf{y})}{\partial \tau_{1,j\neq i} \partial \tau_{2,k\neq\{i,j\}}}\right|_{\mathbf{y}^*}, \end{aligned} \tag{C2b}$$

where $\tau_{n,i}$ denotes the value of trait $\tau_n$ ($n = 1, 2$) expressed by individual $i$, $\tau_{n,j\neq i}$ denotes the value of $\tau_n$ expressed by individual $j$ that is different from $i$, and $\tau_{n,k\neq\{i,j\}}$ denotes the value of $\tau_n$ expressed by yet another neighbour $k$ (that is neither $i$ nor $j$), and $R_3^\circ$ is three-way relatedness under neutrality, i.e., the probability that three randomly sampled individuals from the same patch are IBD in a monomorphic population. The second term in eq. (C2), $h_{\mathrm{r},\tau_1\tau_2}(\mathbf{y}^*)$, is given by

$$\begin{aligned} h_{\mathrm{r},\tau_1\tau_2}(\mathbf{y}^*) = (N-1) \Bigg( & \left.\frac{\partial R_2(\mathbf{y}_i, \mathbf{y}_{-i}, \mathbf{y})}{\partial \tau_{1,i}}\right|_{\mathbf{y}^*} \times \left.\frac{\partial \mathrm{w}(\mathbf{y}_i, \mathbf{y}_{-i}, \mathbf{y})}{\partial \tau_{2,i}}\right|_{\mathbf{y}^*} \\ & + \left.\frac{\partial R_2(\mathbf{y}_i, \mathbf{y}_{-i}, \mathbf{y})}{\partial \tau_{2,i}}\right|_{\mathbf{y}^*} \times \left.\frac{\partial \mathrm{w}(\mathbf{y}_i, \mathbf{y}_{-i}, \mathbf{y})}{\partial \tau_{1,i}}\right|_{\mathbf{y}^*} \Bigg), \end{aligned} \tag{C2c}$$

where

$$\begin{aligned} \left.\frac{\partial R_2(\mathbf{y}_i, \mathbf{y}_{-i}, \mathbf{y})}{\partial \tau_{n,i}}\right|_{\mathbf{y}^*} = 2\frac{R_2^\circ}{1-d_\mathrm{s}} \Bigg[ & \left(1 + (N-1)R_2^\circ\right) \left.\frac{\mathrm{w_p}(\mathbf{y}_i, \mathbf{y}_{-i}, \mathbf{y})}{\tau_{n,i}}\right|_{\mathbf{y}^*} \\ & + \left(2(N-1)R_2^\circ + (N-1)(N-2)R_3^\circ\right) \left.\frac{\mathrm{w_p}(\mathbf{y}_i, \mathbf{y}_{-i}, \mathbf{y})}{\tau_{n,j\neq i}}\right|_{\mathbf{y}^*} \Bigg] \end{aligned} \tag{C2d}$$

is the the effect of trait $\tau_n$ on pairwise relatedness among mutant, in which $\mathrm{w_p}(\mathbf{y}_i, \mathbf{y}_{-i}, \mathbf{y})$ the philopatric fitness of the focal individual, i.e., its expected number of offspring that establish in their natal patch when the focal expresses phenotype $\mathbf{y}_i = (x_i, z_i)$ in a patch where other individuals express $\mathbf{y}_{-i} =$

$(\mathbf{y}_1, \ldots, \mathbf{y}_{i-1}, \mathbf{y}_{i+1}, \ldots, \mathbf{y}_N)$, and the rest of the population expresses $\mathbf{y} = (x, z)$. See Mullon et al. (2016) or Mullon and Lehmann (2019) for a detailed interpretation of eq. (C2).

#### C.2 Individual fitness and relatedness

##### C.2.1 Individual fitness

According to our assumptions given in Section 4.1 of the main text, individual fitness can be written as,

$$\mathrm{w}(\mathbf{y}_i, \mathbf{y}_{-i}, \mathbf{y}) = \mathrm{w}_{♀}(\mathbf{y}_i, \mathbf{y}_{-i}, \mathbf{y}) + \mathrm{w}_{♂}(\mathbf{y}_i, \mathbf{y}_{-i}, \mathbf{y}). \tag{C3a}$$

where $\mathrm{w}_{♀}(\mathbf{y}_i, \mathbf{y}_{-i}, \mathbf{y})$ is the number offspring produced by the focal individual through its own ovules, and $\mathrm{w}_{♂}(\mathbf{y}_i, \mathbf{y}_{-i}, \mathbf{y})$ is that through fertilising of the ovules of other individuals in its patch. The former number is given by

$$\begin{aligned}\mathrm{w}_{♀}(\mathbf{y}_i, \mathbf{y}_{-i}, \mathbf{y}) =& F(\mathbf{y}_i)\left(\underbrace{\frac{M(\mathbf{y}_i)}{\sum_{j=1}^{N} M(\mathbf{y}_j)}}_{\substack{\text{Probability of being}\\ \text{fertilised by its own pollen.}}} + \frac{1}{2}\underbrace{\frac{\sum_{j\neq i} M(\mathbf{y}_j)}{\sum_{j=1}^{N} M(\mathbf{y}_j)}}_{\substack{\text{Probability of being}\\ \text{by the pollen of others.}}}\right) \\ &\times\left(\underbrace{\frac{d_{\mathrm{s}}}{F(\mathbf{y})}}_{\substack{\text{Seeds dispersed}\\ \text{to other patches.}}} + \underbrace{\frac{N(1-d_{\mathrm{s}})}{d_{\mathrm{s}} N F(\mathbf{y}) + (1-d_{\mathrm{s}})\sum_{j=1}^{N} F(\mathbf{y}_j)}}_{\text{Non-dispersed seeds competing locally.}}\right),\end{aligned} \tag{C3b}$$

which can be understood from the labels below each relevant term. Meanwhile, fitness through fertilisation of others' ovules is given by

$$\mathrm{w}_{♂}(\mathbf{y}_i, \mathbf{y}_{-i}, \mathbf{y}) = \frac{1}{2}\left(\sum_{j\neq i} F(\mathbf{y}_j)\right)\frac{M(\mathbf{y}_i)}{\sum_{j=1}^{N} M(\mathbf{y}_j)} \times \left(\frac{d_{\mathrm{s}}}{F(\mathbf{y})} + \frac{N(1-d_{\mathrm{s}})}{d_{\mathrm{s}} N F(\mathbf{y}) + (1-d_{\mathrm{s}})\sum_{j=1}^{N} F(\mathbf{y}_j)}\right), \tag{C3c}$$

where $\sum_{j\neq i} F(\mathbf{y}_j)$ is the number of others' ovules in the patch. The focal individual fertilises a fraction $M(\mathbf{y}_i)/\left(\sum_{j=1}^{N} M(\mathbf{y}_j)\right)$ of these to produce seeds, that then either disperse or compete in their natal patch, as before.

From eq. (C3), philopatric fitness is then given by

$$\begin{aligned}\mathrm{w}_{\mathrm{p}}(\mathbf{y}_i, \mathbf{y}_{-i}, \mathbf{y}) = & \frac{N(1-d_{\mathrm{s}})}{d_{\mathrm{s}} N F(\mathbf{y}) + (1-d_{\mathrm{s}}) \sum_{j=1}^{N} F(\mathbf{y}_j)} \\ & \times \left[ F(\mathbf{y}_i) \left( \frac{M(\mathbf{y}_i)}{\sum_{j=1}^{N} M(\mathbf{y}_j)} + \frac{1}{2} \frac{\sum_{j\neq i}^{N} M(\mathbf{y}_j)}{\sum_{j=1}^{N} M(\mathbf{y}_j)} \right) + \frac{1}{2} \frac{M(\mathbf{y}_i)}{\sum_{j=1}^{N} M(\mathbf{y}_j)} \sum_{j\neq i} F(\mathbf{y}_j) \right].\end{aligned} \tag{C4}$$

##### C.2.2 Relatedness

The approach described in Section C.1 also relies on the pairwise $R_2^{\circ}$ and three-way $R_3^{\circ}$ relatedness coefficients. Under neutrality, our model is equivalent to a standard haploid Wright-Fisher model, such that we can use previous results and have

$$\begin{aligned} R_2^{\circ} &= \frac{(1-d_{\mathrm{s}})^2}{1 + d_{\mathrm{s}}(2-d_{\mathrm{s}})(N-1)} \\ R_3^{\circ} &= \frac{(1-d_{\mathrm{s}})^3 \left\{3N - 2\left[1 + d_{\mathrm{s}}(2-d_{\mathrm{s}})(N-1)\right]\right\}}{\left[1 + d_{\mathrm{s}}(2-d_{\mathrm{s}})(N-1)\right]\left[N^2 - (1-d_{\mathrm{s}})^3(N-2)(N-1)\right]}, \end{aligned} \tag{C5}$$

(e.g., Ohtsuki, 2010).

#### C.3 Analyses

##### C.3.1 Evolutionary attractor

Inserting eqs. (C3) and eq. (C5) into eq. (C1), we obtain

$$s_x(\mathbf{y}) = \frac{d_{\mathrm{s}}(2-d_{\mathrm{s}})[1 + N(1-2x)]}{2x(1-x)[1 + d_{\mathrm{s}}(2-d_{\mathrm{s}})(N-1)]} \tag{C6a}$$

$$s_z(\mathbf{y}) = \frac{d_{\mathrm{s}}(2-d_{\mathrm{s}})}{2\left[N - (1-d_{\mathrm{s}})^2(N-1)\right]} \left[(N+1)\frac{V'_{♂}(z)}{V_{♂}(z)} + (N-1)\frac{V'_{♂}(z)}{V_{♂}(z)}\right], \tag{C6b}$$

for the selection gradients on sex allocation $x$ and trait $z$, respectively. Solving those for zero, we obtain the equilibrium trait values

$$x^* = \frac{1}{2} + \frac{1}{2N}, \tag{C7}$$

for sex allocation, and that $z^*$ must be such that

$$\frac{V'_{♀}(z)}{V_{♀}(z)} = -\frac{N-1}{N+1}\frac{V'_{♂}(z)}{V_{♂}(z)}, \tag{C8}$$

which are given in eq. (15) of the main text. Note that with Gaussian functions (eq. 2), we have

$$z^* = \theta_{♀}\frac{\omega_{♀}(N+1)}{\omega_{♀}(N+1)+\omega_{♂}(N-1)} + \theta_{♂}\frac{\omega_{♂}(N-1)}{\omega_{♀}(N+1)+\omega_{♂}(N-1)}, \tag{C9}$$

which becomes increasingly biased towards the female optimum $\theta_{♀}$ as $N$ decreases, as expected from eq. (C8) (see Fig. 5B for illustration).

As per usual, the equilibrium $\mathbf{y}^* = (x^*, z^*)$ is convergence stable if the leading eigenvalue of $\mathbf{J}(\mathbf{y}^*)$, the Jacobian matrix (eq. A8), is negative. Using eqs. (C6)-(C8), this condition is equivalent to

$$\frac{d_{\mathrm{s}}(2-d_{\mathrm{s}})}{2\left[N-(N-1)(1-d_{\mathrm{s}})^2\right]}\left[(N+1)\frac{V''_{♀}(z^*)}{V_{♀}(z^*)} + (N-1)\frac{V''_{♂}(z^*)}{V_{♂}(z^*)} + 2N\frac{V'_{♀}(z^*)}{V_{♀}(z^*)}\frac{V'_{♀}(z^*)}{V_{♀}(z^*)}\right] < 0, \tag{C10}$$

which is always the case with Gaussian functions, as eq. (C10) then reduces to

$$-\frac{d_{\mathrm{s}}(2-d_{\mathrm{s}})}{N-(N-1)(1-d_{\mathrm{s}})^2}\left[\omega_{♀}(N+1)+\omega_{♂}(N-1)\right] < 0, \tag{C11}$$

which is always true.

##### C.3.2 Evolutionary branching

To compute the Hessian matrix for our model, we insert eqs. (C3) to (C5) into eq. (C2). The resulting matrix is complicated. We thus refrain from providing it here, but it is available in the *Mathematica* Notebook “AppendixC-Limited_Dispersal.nb” provided in Supplementary Material. To investigate the influence of limited dispersal in the emergence of polymorphism, we consider the large patches, low seed dispersal limit that yields simpler mathematical expressions (see eq. (16) in the main text) and explore the conditions for polymorphism numerically (see Fig. 5 in the main text).

#### Appendix D

### The effects of self-fertilisation and inbreeding depression

Following the assumptions detailed in Section 4.2 of the main text, the invasion fitness $W(\mathbf{y}_{\mathrm{m}}, \mathbf{y})$ of a rare mutant $\mathbf{y}_{\mathrm{m}} = (x_{\mathrm{m}}, z_{\mathrm{m}})$ in a resident population fixed for $\mathbf{y} = (x, z)$ is given by

$$W(\mathbf{y}_{\mathrm{m}}, \mathbf{y}) = \frac{1}{F(\mathbf{y})(1 - A(\mathbf{y}, \mathbf{y})\delta)} \left[ F(\mathbf{y}_{\mathrm{m}}) \left( A(\mathbf{y}_{\mathrm{m}}, \mathbf{y})(1 - \delta) + \frac{1 - A(\mathbf{y}_{\mathrm{m}}, \mathbf{y})}{2} \right) + F(\mathbf{y}) \frac{1 - A(\mathbf{y}, \mathbf{y})}{2} \frac{M(\mathbf{y}_{\mathrm{m}})}{M(\mathbf{y})} \right], \tag{D1a}$$

where $A(\mathbf{y}_i, \mathbf{y}_{-i})$ is the selfing rate the $i^{\mathrm{th}}$ individual in the population, and is given by

$$A(\mathbf{y}_i, \mathbf{y}_{-i}) = \frac{\alpha M(\mathbf{y}_i)}{\alpha M(\mathbf{y}_i) + \frac{1 - \alpha}{N - 1} \sum_{j \neq i} M(\mathbf{y}_j)}, \tag{D1b}$$

where $\mathbf{y}_{-i}$ denotes the strategies of the $N - 1$ other individuals in the population, so that, for a rare mutant, we have

$$A(\mathbf{y}_{\mathrm{m}}, \mathbf{y}) = \frac{\alpha M(\mathbf{y}_{\mathrm{m}})}{\alpha M(\mathbf{y}_{\mathrm{m}}) + (1 - \alpha)\mathbf{y}}. \tag{D1c}$$

Eq. (D1a) can be understood as follows. A mutant gains fitness through female function by producing $F(\mathbf{y}_{\mathrm{m}})$ ovules. A fraction $A(\mathbf{y}_{\mathrm{m}}, \mathbf{y})$ of these ovules gets self-fertilised, leading to self-fertilised zygotes that survive with probability $1 - \delta$. The surviving selfed zygotes then divide to produce seeds that all carry the mutant haplotype. The remaining fraction of ovules $1 - A(\mathbf{y}_{\mathrm{m}}, \mathbf{y})$ gets outcrossed, giving rise to outcrossed zygotes that survive with probability 1, and then divide to produce seeds that carry the mutant haplotype with probability $1/2$. The mutant also gains fitness through its male function by fertilising resident individuals' ovules $F(\mathbf{y})$ that have not been self-fertilised, which it does in proportion $M(\mathbf{y}_{\mathrm{m}})/M(\mathbf{y})$. Interestingly, eq. (D1a) is equivalent to individual fitness under limited dispersal (eq. C3) when $\delta = 0$, $d_{\mathrm{s}} = 1$ and $\alpha = 1/N$.

#### D.1 Directional selection

**Selection gradients and singular strategies.** As before, the selection gradients $s_x(\mathbf{y})$ and $s_z(\mathbf{y})$ acting on $x$ and $z$ can be computed using eq. (A2). We find that the selection gradient on sex allocation is given by

$$s_x(\mathbf{y}) = \frac{1+\alpha(1-2\delta)}{2x(1-x)(1-\alpha\delta)}\big[1+x(2-\alpha)\big], \tag{D2}$$

which, solving for the singular sex allocation strategy $x^*$ (eq. A3), yields

$$x^* = \frac{1}{2-\alpha}. \tag{D3}$$

Eq. (D3) corresponds to eq. (18a) in the main text (see also Fig. 6A). The selection gradient on trait $z$, meanwhile, is

$$s_z(\mathbf{y}) = \left.\frac{\partial W}{\partial z_{\mathrm{m}}}\right|_{\mathbf{y}_{\mathrm{m}}=\mathbf{y}} = \frac{1+\alpha(1-2\delta)}{2(1-\alpha\delta)}\left[\frac{V'_{♀}(z)}{V_{♀}(z)} + (1-\alpha)\frac{V'_{♂}(z)}{V_{♂}(z)}\right]. \tag{D4}$$

Eq. (D4) shows that selfing interferes with directional selection on $z$ through a similar effect to the one selecting for female-biased sex allocation (eq. D3), i.e., by reducing the contribution to fitness of male function relative to female function, so that the singular trait value $z^*$ (eq. A3) now satisfies

$$\frac{V'_{♀}(z^*)}{V_{♀}(z^*)} = -(1-\alpha)\frac{V'_{♂}(z^*)}{V_{♂}(z^*)}, \tag{D5}$$

which corresponds to eq. (18b) in the main text. With Gaussian $V_u(z)$ functions (eq. 2), this yields

$$z^* = \theta_{♂}\frac{(1-\alpha)\omega_{♀}}{\omega_{♂}+(1-\alpha)\omega_{♀}} + \theta_{♀}\frac{\omega_{♂}}{\omega_{♂}+(1-\alpha)\omega_{♀}}, \tag{D6}$$

which shows that increasing $\alpha$ moves $z^*$ closer to the female optimum $\theta_{♀}$, as shown in Figure 6B in the main text.

**Convergence stability.** The Jacobian matrix evaluated at $\mathbf{y} = \mathbf{y}^*$ (eq. A8) is given by

$$\mathbf{J}(\mathbf{y}^*) = \begin{pmatrix} -\dfrac{(2-\alpha)^3[1+\alpha(1-2\delta)]}{2(1-\alpha)(1-\alpha\delta)} & 0 \\ 0 & \dfrac{1+\alpha(1-2\delta)}{2(1-\alpha\delta)}\left(\dfrac{V''_{♀}(z^*)}{V_{♀}(z^*)} + (1-\alpha)\dfrac{V''_{♂}(z^*)}{V_{♂}(z^*)} + (2-\alpha)\dfrac{V'_{♀}(z^*)}{V_{♀}(z^*)}\dfrac{V'_{♂}(z^*)}{V_{♂}(z^*)}\right) \end{pmatrix}. \tag{D7}$$

Because the matrix is diagonal (i.e., only elements on its diagonal differ from zero), its eigenvalues are given by its diagonal elements, which implies that $\mathbf{y}^*$ is convergence stable if these elements are both negative. Similar to the complete outcrossing case, $\mathbf{y}^*$ is always convergence stable provided that functions $V_{♀}(z)$ and $V_{♂}(z)$ are not too accelerating relative to their rate of increase close to $z^*$. This is always the case with Gaussian functions (eq. 2), as we have

$$\left.\frac{\partial s_z(\mathbf{y})}{\partial z}\right|_{\mathbf{y}=\mathbf{y}^*} = -\frac{[\omega_{♀} + \omega_{♂}(1-\alpha)][1+\alpha(1-2\delta)]}{1-\alpha\delta}. \tag{D8}$$

#### D.2 Disruptive selection

To study selection on $x$ and $z$ once the population expresses $\mathbf{y}^*$, we apply eq. (A12) to eq. (D1a), and obtain the Hessian matrix presented in main text equation (19) in Section 4.2.

#### Appendix E

### Diploidy and the evolution of dominance

In this Appendix, we explore the consequences of diploidy on our results. We first reanalyse our baseline model in diploids analytically (section E.1.1), and then present simulations where dominance is allowed to evolve at the loci encoding sex allocation and trait $z$ (section E.1.2). Next, we extend our model of conditional trait expression evolution (section B.2) to investigate the joint evolution of dominance and conditional expression of trait $z$ when the loci encoding these traits are unlinked (section E.2) . Finally, we show that the stabilising effect of selfing on hermaphroditism is unaffected by ploidy in our model (section E.3).

#### E.1 Baseline model under diploidy

In this section, we extend our baseline model to include diploidy. We first derive the conditions for correlational selection to favour the joint emergence of separate sexes and sexual dimorphism, and we then use computer simulations to study the evolution of dominance when these conditions are met.

##### E.1.1 Invasion analysis

The life cycle is identical to Section 2.1 in the main text, except that we assume individuals to be diploid, which entails two modifications. First, during *(i) Sexual development*, diploid individuals produce a large number of haploid female and male gametes through meiosis. Second, *(ii) Mating* gives rise to a large number of diploid zygotes that mature into diploid seeds instead of dividing into haploid ones.

As before, we assume that sex allocation $x$ and trait $z$ are each determined by a quantitative trait locus evolving under a 'continuum-of-alleles' model, and that these loci are fully linked. Alleles at each locus are additive, meaning that the sex allocation $x$ and conflict trait $z$ of an individual with genotype $x_1/x_2$ and $z_1/z_2$ at encoding loci are given by

$$x = \frac{x_1 + x_2}{2} \quad \text{and} \quad z = \frac{z_1 + z_2}{2}. \tag{E1}$$

We later allow for dominance evolution.

**Invasion fitness.** Under the assumptions described above, the invasion fitness $W(\mathbf{y}_\mathrm{m}, \mathbf{y})$ of a rare mutant haplotype encoding $\mathbf{y}_\mathrm{m} = (x_\mathrm{m}, z_\mathrm{m})$ in a resident population otherwise fixed for $\mathbf{y} = (x, z)$ is simply given by

$$W(\mathbf{y}_\mathrm{m}, \mathbf{y}) = \frac{1}{2}\left(\frac{\frac{x + x_\mathrm{m}}{2} V_{♀}\left(\frac{z + z_\mathrm{m}}{2}\right)}{x\, V_{♀}(z)} + \frac{\left(1 - \frac{x + x_\mathrm{m}}{2}\right) V_{♂}\left(\frac{z + z_\mathrm{m}}{2}\right)}{(1 - x) V_{♂}(z)}\right). \tag{E2}$$

This is because, under random mating in a large population, a rare mutant only ever occurs in heterozygous state.

**Directional selection.** Using eq. (A2), we find that the selection gradient on $x$ and $z$, $s_x(\mathbf{y})$ and $s_z(\mathbf{y})$, are given by

$$s_x(\mathbf{y}) = \frac{1 - 2x}{4x(1 - x)} \quad \text{and} \quad s_z(\mathbf{y}) = \frac{1}{4}\left(\frac{V'_{♀}(z)}{V_{♀}(z)} + \frac{V'_{♂}(z)}{V_{♂}(z)}\right), \tag{E3}$$

respectively, which correspond to the selection gradients obtained in the haploid case (eq. A4), multiplied by $1/2$. This $1/2$ factor reflects the fact that the mutant's effect on phenotype is halved in heterozygotes, which reduces the efficacy of selection on it. Because the gradients in the diploid case only differ by a positive constant from the haploid case, the singular sex allocation $x^*$ and trait value $z^*$ are unchanged, and convergence stability is guaranteed by the same arguments as in Appendix A.1.1.

**Disruptive selection.** Once an haplotype encoding the singular phenotype $\mathbf{y}^* = (x^*, z^*)$ fixes in the population, whether or not selection favours the emergence of polymorphism is determined by the leading eigenvalue of the Hessian matrix. Using eq. (A12), this matrix is given by

$$\mathbf{H}(\mathbf{y}^*) = \begin{pmatrix} 0 & \frac{1}{4}\left(\frac{V'_{♀}(z^*)}{V_{♀}(z^*)} - \frac{V'_{♂}(z^*)}{V_{♂}(z^*)}\right) \\ \frac{1}{4}\left(\frac{V'_{♀}(z^*)}{V_{♀}(z^*)} - \frac{V'_{♂}(z^*)}{V_{♂}(z^*)}\right) & \frac{1}{8}\left(\frac{V''_{♀}(z^*)}{V_{♀}(z^*)} + \frac{V''_{♂}(z^*)}{V_{♂}(z^*)}\right) \end{pmatrix}, \tag{E4}$$

which corresponds to the Hessian matrix under haploidy (eq. A13) multiplied by $1/4$. It follows immediately that the results obtained on the conditions for the emergence of polymorphism are identical to the haploid case.

##### E.1.2 The evolution of dominance

In this section, we investigate the evolution of dominance at the loci encoding sex allocation $x$ and trait $z$ using computer simulations.

###### E.1.2.1 Simulation program

The simulation program is coded in C++. It simulates a population of diploids with a constant size $N$, in which each individual is characterised by its diploid genotype at the sex allocation and the trait locus. To include dominance evolution, we assume that each locus contains two linked components, a promoter sequence and a gene. At the gene, alleles can encode any sex allocation $x \in [0, 1]$ or trait value $z \in \mathbb{R}$, as before. The promoter sequence, meanwhile, encodes dominance relationships between alleles at the gene following an 'affinity' model (Van Dooren, 1999; Lesaffre et al., 2024). This model assumes that promoters at each locus are characterised by their affinity with transcription factors, which we denote as $\phi_\tau \in ]0, +\infty)$, $\tau \in \{x, z\}$. Transcription factors must bind to the promoter linked to an allele for this allele to be expressed, and are assumed to do so proportionately to promoter affinity. Therefore, the phenotype $\tau \in \{x, z\}$ of an individual with genotype $\{\phi_{\tau,1}, \tau_1\}/\{\phi_{\tau,2}, \tau_2\}$ at the corresponding locus is given by

$$\tau = \tau_1 \frac{\phi_{\tau,1}}{\phi_{\tau,1} + \phi_{\tau,2}} + \tau_2 \frac{\phi_{\tau,2}}{\phi_{\tau,1} + \phi_{\tau,2}}, \tag{E5}$$

so that alleles associated with higher affinity promoters tend to be over-expressed in heterozygotes, i.e., to be dominant, and alleles associated with lower affinity promoters tend to be recessive.

The population is initialised as monomorphic, with promoters at both loci characterised by an affinity $\phi_0 = 1$, and alleles at the sex allocation and trait genes encoding values $x_0 \in (0, 1)$ and $z_0 \in \mathbb{R}$, respectively. At each time step, we create the next generation by generating $N$ diploid offspring. For each offspring, we sample a maternal and a paternal parent with replacement from the previous generation. Parents are sampled from the female or male gamete pool in proportion to their sex-specific fecundity, and they each randomly transmit one of their chromosomes. Each time an offspring is produced, the alleles it carries at the genes and the promoters each undergo mutation with probability $\mu$, in which case the new value encoded by the mutated allele is sampled in a Gaussian distribution centred on the parental value with standard deviation $\sigma$, truncated such that allelic values are kept within bounds (i.e., between zero and one for sex allocation, and above zero for promoter affinities). We let simulations run for $t_{\max}$ generations. Every $t_{\text{mes}}$ generations, we record the genotype of $n_{\text{mes}}$ randomly sampled individuals in the population at the sex allocation and conflict trait loci.

###### E.1.2.2 Dominance evolution leads to the emergence of a sex-determining locus

We ran simulations for several combinations of $\omega_{♀}$ and $\omega_{♂}$. Out of a thousand simulation runs total (200 per parameter set), we found that polymorphism and dominance evolved in all of them, at both loci. In these simulations, evolutionary branching leads to the emergence of two haplotypes: one specialised into female function which encodes complete femaleness ($x = 1$) and a female-biased trait value; and another specialised into male function, encoding complete maleness ($x = 0$) and a male-biased trait value.

At the locus encoding trait $z$, one allele ends up encoding the optimal value for one sex (typically the allele beneficial to the sex where sex-specific selection was weakest), such that homozygotes for that allele expressed the optimal trait value for that sex. The other allele 'overshoots' the optimum for the other sex, and dominance evolves such that heterozygotes express the optimal trait value for that other sex. At the sex allocation locus, meanwhile, alleles encoding complete femaleness ($x = 1$) and complete maleness ($x = 0$) emerge, and one becomes fully dominant over the other, so that either an XY (male dominant) or a ZW (female dominant) sex-determining locus always evolves. Figure S2 shows an example where selection is stronger through female than male function $\omega_{♀} > \omega_{♂}$. Here, the female-beneficial allele overshoots the female optimum at the conflict trait locus and dominance of the female allele evolves at the sex allocation locus, leading to a ZW system.

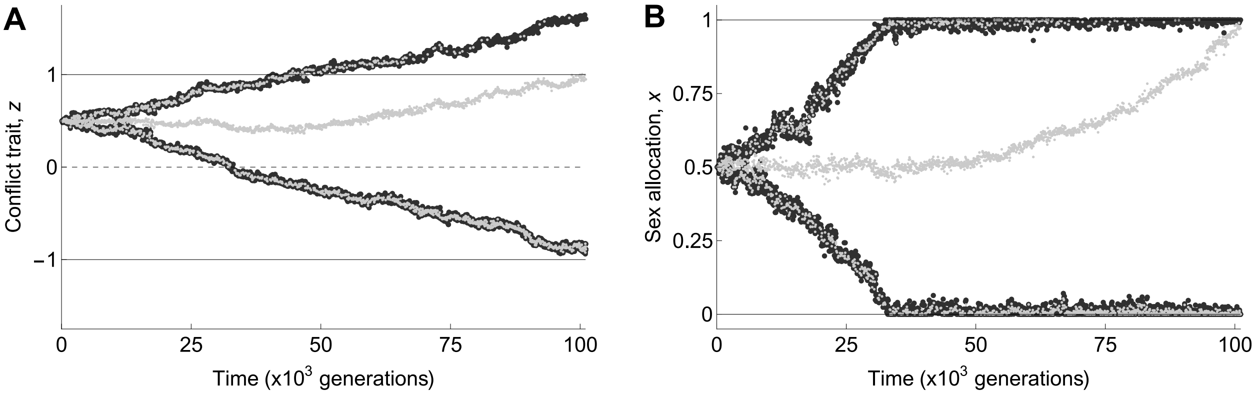

Figure S2: Evolution of dominance in diploids. In each plot, large black dots indicate the values encoded by the alleles borne by 50 randomly sampled individuals in the population, and smaller grey dots indicate the phenotype expressed by these same 50 individuals. **A** Evolution of dominance at the conflict trait locus. Solid lines indicate sex-specific optima $\theta_{♀} = 1$ and $\theta_{♂} = -1$. Following evolutionary branching, the allele encoding a male-beneficial trait value stabilises close to the male optimum, whereas the allele for a more female-beneficial trait value overshoots the female optimum and stabilises at a higher value. Dominance then evolves such that the heterozygote genotype expresses the female optimum $\theta_{♀} = 1$. **B** Evolution of dominance at the sex allocation locus. Alleles for complete femaleness ($x = 1$) and complete maleness ($x = 0$) emerge through evolutionary branching, and dominance of the female allele subsequently evolves. Parameters used in this simulation: $\omega_{♀} = 0.6$, $\omega_{♂} = 0.2$, $\theta_{♀} = 1$, $\theta_{♂} = -1$, $N = 10^3$, $\mu = 10^{-3}$ and $\sigma = 10^{-2}$.

#### E.2 Evolution of conditional trait expression in diploids

We now extend our model of conditional trait expression evolution to the diploid case (Appendix B.2) and allow dominance to evolve using computer simulations.

##### E.2.1 Simulation program

The simulation program is coded in C++. It is modified from the program described in Appendix B.2.3. It simulates a population of $N$ diploid individuals. Each individual is characterised by its diploid genotype at three unlinked loci encoding its sex allocation $x \in [0, 1]$, its sex-dependent trait component $a \in \mathbb{R}$, and its independent trait component $b \in \mathbb{R}$, respectively. Given $x$, $a$ and $b$, the trait $z$ value expressed by an individual is given by

$$z = a \cdot x + b. \tag{E6}$$

Each locus contains a gene and a promoter. The promoters determines dominance relationships within each locus following the same affinity model as described in Appendix E.1.2.1 (eq. E5; Van Dooren, 1999; Lesaffre et al., 2024).

The population is monomorphic at the beginning of the simulation, with promoters characterised by an affinity $\phi_0 = 1$ and genes fixed for alleles $x_0 \in (0, 1)$, $a_0 \in \mathbb{R}$ and $b_0 \in \mathbb{R}$, respectively. Each new generation is formed by creating $N$ diploid offspring. For each of them, we sample maternal and paternal parents with replacement in proportion to their sex-specific fecundities. As we assume free recombination, each parent randomly transmits one of its alleles at each locus. These alleles then each undergo mutation with probability $\mu$, and their new value is sampled in a Gaussian distribution centred on the parental trait with standard deviation $\sigma$, truncated such that allelic values remain within the definition domain of the corresponding trait. We let simulations run for $t_{\mathrm{max}}$ generations. We record the genotype of $n_{\mathrm{mes}}$ randomly sampled individuals in the population every $t_{\mathrm{mes}}$ generations at all loci.

##### E.2.2 Conditional expression and dominance readily evolve in diploids

We ran many replicates for various strengths of sex-specific selection ($\omega_{♀}$ and $\omega_{♂}$). We found that sex allocation-dependent expression of trait $z$ began to evolve in response to standing variation for $x$, which then triggered evolutionary branching at the sex allocation locus and the emergence of alleles encoding pure femaleness ($x = 1$) and pure maleness ($x = 0$). Once these alleles were sufficiently diverged, complete dominance of one allele over the other, leading to the emergence of a heterogametic sex-determining locus. The loci encoding $a$ and $b$, meanwhile, remained monomorphic and gradually converged to allelic values that allowed female and male individuals to express their sex-specific optima, i.e.,

$$a^* = \theta_{♀} - \theta_{♂} \quad \text{and} \quad b^* = \theta_{♂}, \tag{E7}$$

as in the haploid case. Figure S3 shows a typical simulation run where an XY system and sex-dependent expression of trait $z$ evolve simultaneously.

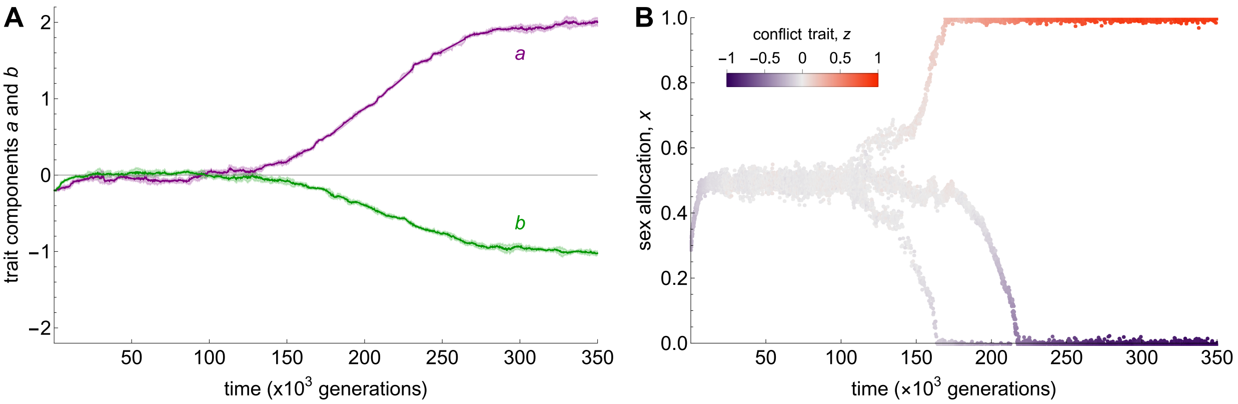

Figure S3: Evolution of conditional trait expression and dominance in diploids. **A** Evolution of conditional trait expression. The purple and green lines show the average value of $a$ and $b$ expressed by individuals in the population as a function of time. The shaded areas indicate two standard deviations around these means. **B** Evolution of sex allocation as a function of time. Each dot represents an individual, and is coloured according to its conflict trait value.

#### E.3 The interplay between selfing and diploidy

Here we extend our model to investigate the joint evolution of sex allocation and trait $z$ under partial selfing in diploids. The life cycle is identical to the one described in Section 4.2 of the main text and analysed in Appendix D, except that we assume individuals to be diploid, so that gamete fusion results in the production of diploid zygotes that suffer from inbreeding depression if they are self-fertilised, and mature into diploid seeds directly instead of dividing into two haploid seeds.

We assume that sex allocation $x$ and trait $z$ are each encoded by a quantitative trait locus following a 'continuum-of-alleles' model. Alleles are additive at each locus, meaning that the maternally and paternally inherited copies contribute equally to phenotype. The two loci are assumed to be in complete linkage, as in our baseline model.

##### E.3.1 Invasion matrix and invasion fitness

In a partially selfing diploid population, a rare mutant haplotype $\mathbf{y}_{\mathrm{m}}$ can be present in heterozygous state with the resident allele $\mathbf{y}$, but also in homozygous state, because it is now likely to mate with itself even when the population is very large owing to selfing. Therefore, the dynamics of the mutant sub-population are described by the recursion

$$\mathbf{n}_{t+1} = \mathbf{W}(\mathbf{y}_{\mathrm{m}}, \mathbf{y}) \cdot \mathbf{n}_t, \tag{E8}$$

where $\mathbf{n}_t = (n_{1,t}, n_{2,t})$ is a vector that gives the number of mutant heterozygotes ($n_{1,t}$) and mutant homozygotes ($n_{2,t}$) in the population at time $t$, and $\mathbf{W}(\mathbf{y}_{\mathrm{m}}, \mathbf{y})$ is a $2 \times 2$ matrix, the $(i, j)$-entry of which $w_{ij}(\mathbf{y}_{\mathrm{m}}, y)$ gives the expected number of successful mutant offspring of class $i$ produced by a mutant of class $j$.

Analogously to eq. (1) of the main text, we define the female and male fecundities of individual $i$ carrying haplotypes $\mathbf{y}_{i,1} = (x_{i,1}, z_{i,1})$ and $\mathbf{y}_{i,2} = (x_{i,2}, z_{i,2})$ in the population as

$$F(\mathbf{y}_{i,1}, \mathbf{y}_{i,2}) = \frac{x_{i,1} + x_{i,2}}{2} V_{♀}\left(\frac{z_{i,1} + z_{i,2}}{2}\right) \quad \text{and} \quad M(\mathbf{y}_{i,1}, \mathbf{y}_{i,2}) = \left(1 - \frac{x_{i,1} + x_{i,2}}{2}\right) V_{♂}\left(\frac{z_{i,1} + z_{i,2}}{2}\right), \tag{E9}$$

and the selfing rate of this individual analogously to eq. (D1b) in Appendix D, as

$$A(\mathbf{y}_{i,1}, \mathbf{y}_{i,2} | \mathbf{y}_{-i}) = \frac{\alpha M(\mathbf{y}_{i,1}, \mathbf{y}_{i,2})}{\alpha M(\mathbf{y}_{i,1}, \mathbf{y}_{i,2}) + \frac{1-\alpha}{N-1} \sum_{j \neq i} M(\mathbf{y}_{j,1}, \mathbf{y}_{j,2})}, \tag{E10}$$

where $\mathbf{y}_{-i}$ denotes the genotype of the $N - 1$ other individuals in the population.

Using these notations, the entries of $\mathbf{W}(\mathbf{y}_{\mathrm{m}}, \mathbf{y})$ are given by

$$w_{11}(\mathbf{y}_{\mathrm{m}}, \mathbf{y}) = F(\mathbf{y}_{\mathrm{m}}, \mathbf{y}) \left[A(\mathbf{y}_{\mathrm{m}}, \mathbf{y}|\mathbf{y}) \frac{1-\delta}{2} + \left[1 - A(\mathbf{y}_{\mathrm{m}}, \mathbf{y}|\mathbf{y})\right] \frac{1}{2}\right] + F(\mathbf{y}, \mathbf{y}) \left[1 - A(\mathbf{y}, \mathbf{y}|\mathbf{y})\right] \frac{M(\mathbf{y}_{\mathrm{m}}, \mathbf{y})}{M(\mathbf{y}, \mathbf{y})} \frac{1}{2}, \tag{E11a}$$

$$w_{12}(\mathbf{y}_{\mathrm{m}}, \mathbf{y}) = F(\mathbf{y}_{\mathrm{m}}, \mathbf{y}_{\mathrm{m}}) \left[1 - A(\mathbf{y}_{\mathrm{m}}, \mathbf{y}_{\mathrm{m}}|\mathbf{y})\right] + F(\mathbf{y}, \mathbf{y}) \left[1 - A(\mathbf{y}, \mathbf{y}|\mathbf{y})\right] \frac{M(\mathbf{y}_{\mathrm{m}}, \mathbf{y}_{\mathrm{m}})}{M(\mathbf{y}, \mathbf{y})}, \tag{E11b}$$

$$w_{21}(\mathbf{y}_{\mathrm{m}}, \mathbf{y}) = F(\mathbf{y}_{\mathrm{m}}, \mathbf{y}) A(\mathbf{y}_{\mathrm{m}}, \mathbf{y}|\mathbf{y}) \frac{1-\delta}{4}, \tag{E11c}$$

$$w_{22}(\mathbf{y}_{\mathrm{m}}, \mathbf{y}) = F(\mathbf{y}_{\mathrm{m}}, \mathbf{y}_{\mathrm{m}}) A(\mathbf{y}_{\mathrm{m}}, \mathbf{y}_{\mathrm{m}}|\mathbf{y})(1-\delta). \tag{E11d}$$

To understand how these entries were obtained, let us consider eq. (E11a), which gives the expected number of successful heterozygous mutant offspring produced by a focal heterozygous mutant. Such a mutant can produce heterozygous offspring in three distinct way. First, it self-fertilises a fraction $A(\mathbf{y}_{\mathrm{m}}, \mathbf{y}|\mathbf{y})$ of its ovules, and the resulting offspring then have a 50% chance of being heterozygous themselves (hence the $1/2$ factor) but may die due to inbreeding depression $(1-\delta)$. Second, their remaining ovules $(1 - A(\mathbf{y}_{\mathrm{m}}, \mathbf{y}|\mathbf{y}))$ are fertilised by resident pollen, in which case the mutant allele is transmitted $1/2$ of the time. Third, they may fertilise available resident ovules, in which the mutant allele is again transmitted to a fraction $1/2$ of the resulting offspring.

The leading eigenvalue of $\mathbf{W}(\mathbf{y}_{\mathrm{m}}, \mathbf{y})$, which we denote as $\rho(\mathbf{y}_{\mathrm{m}}, \mathbf{y})$, is the invasion fitness of the mutant. Its expression can be computed explicitly, but it is complicated. We therefore refrain from giving it here, but it is available in the accompanying Mathematica Notebook "AppendixE3-Selfing_Diploids.nb"

##### E.3.2 Directional selection

We compute the selection gradients on $x$ and $z$ by applying eq. (A2) to $\rho(\mathbf{y}_{\mathrm{m}}, \mathbf{y})$, which yields

$$s_x(\mathbf{y}) = \frac{1+\alpha(1-2\delta)}{2x(1-x)[2-\alpha(1+\delta)]}[1-x(2-\alpha)] \tag{E12a}$$

and

$$s_z(\mathbf{y}) = \frac{1+\alpha(1-2\delta)}{2[2-\alpha(1+\delta)]}\left(\frac{V'_{♀}(z)}{V_{♀}(z)} + (1-\alpha)\frac{V'_{♂}(z)}{V_{♂}(z)}\right). \tag{E12b}$$

Eqs. (E12a) and (E12b) only differ from the haploid case (eqs. D2 and D4 in Appendix D) by a positive constant. As a result, they lead to the same convergence stable singular sex allocation $x^*$ and trait value $z^*$, which are given by eqs. (D3) and (D5) in Appendix D.

##### E.3.3 Disruptive selection

Using eq. (A12), we find that the elements of the Hessian matrix are given by

$$
\begin{aligned}
h_{xx}(x^*) &= -\alpha\frac{(2-\alpha)^2(1-2\delta)}{(1-\alpha)(1-\alpha\delta)} \times \frac{1+\alpha(1-2\delta)}{2[2-\alpha(1+\delta)]} \\
h_{xz}(x^*) &= \left\{-\alpha^2\frac{(2-\alpha)(1-2\delta)}{2(1-\alpha)(1-\alpha\delta)}\frac{V'_{♀}(z^*)}{V_{♀}(z^*)} + \frac{2-\alpha}{2(1-\alpha\delta)}\left(\frac{V'_{♀}(z^*)}{V_{♀}(z^*)} - \frac{V'_{♂}(z^*)}{V_{♂}(z^*)}\right)\right\} \times \frac{1+\alpha(1-2\delta)}{2[2-\alpha(1+\delta)]} \\
h_{zz}(x^*) &= \left\{\alpha\frac{1-2\delta}{1-\alpha\delta}\frac{V'_{♀}(z^*)}{V_{♀}(z^*)}\frac{V'_{♂}(z^*)}{V_{♂}(z^*)} + \frac{1+\alpha(1-2\delta)}{2(1-\alpha\delta)}\left(\frac{V''_{♀}(z^*)}{V_{♀}(z^*)} + (1-\alpha)\frac{V''_{♂}(z^*)}{V_{♂}(z^*)}\right)\right\} \times \frac{1+\alpha(1-2\delta)}{2[2-\alpha(1+\delta)]},
\end{aligned}
\tag{E13}
$$

which correspond to the elements of the Hessian matrix under partial selfing in a population of haploid individuals presented in eq. (19) in the main text, scaled by a positive constant. Therefore, the conditions for selection to favour polymorphism are identical under haploidy and diploidy in our model.
